# Breast Cancer Macrophage Heterogeneity and Self-renewal are Determined by Spatial Localization

**DOI:** 10.1101/2023.10.24.563749

**Authors:** Nir Ben-Chetrit, Xiang Niu, Jesus Sotelo, Ariel D. Swett, Vinagolu K. Rajasekhar, Maria S. Jiao, Caitlin M. Stewart, Priya Bhardwaj, Sanjay Kottapalli, Saravanan Ganesan, Pierre-Louis Loyher, Catherine Potenski, Assaf Hannuna, Kristy A. Brown, Neil M. Iyengar, Dilip D. Giri, Scott W. Lowe, John H. Healey, Frederic Geissmann, Irit Sagi, Johanna A. Joyce, Dan A. Landau

**Author notes:** **Correspondence:** Dan Avi Landau and Nir Ben-Chetrit. (D.A.L), (N.B.-C). These authors contributed equally (Nir Ben-Chetrit, Xiang Niu).

## Abstract

Tumor-infiltrating macrophages support critical steps in tumor progression, and their accumulation in the tumor microenvironment (TME) is associated with adverse outcomes and therapeutic resistance across human cancers. In the TME, macrophages adopt diverse phenotypic alterations, giving rise to heterogeneous immune activation states and induction of cell cycle. While the transcriptional profiles of these activation states are well-annotated across human cancers, the underlying signals that regulate macrophage heterogeneity and accumulation remain incompletely understood. Here, we leveraged a novel *ex vivo* organotypic TME (oTME) model of breast cancer, *in vivo* murine models, and human samples to map the determinants of functional heterogeneity of TME macrophages. We identified a subset of F4/80^high^Sca-1+ self-renewing macrophages maintained by type-I interferon (IFN) signaling and requiring physical contact with cancer-associated fibroblasts. We discovered that the contact-dependent self-renewal of TME macrophages is mediated via Notch4, and its inhibition abrogated tumor growth of breast and ovarian carcinomas *in vivo*, as well as lung dissemination in a PDX model of triple-negative breast cancer (TNBC). Through spatial multi-omic profiling of protein markers and transcriptomes, we found that the localization of macrophages further dictates functionally distinct but reversible phenotypes, regardless of their ontogeny. Whereas immune-stimulatory macrophages (CD11C+CD86+) populated the tumor epithelial nests, the stroma-associated macrophages (SAMs) were proliferative, immunosuppressive (Sca-1+CD206+PD-L1+), resistant to CSF-1R depletion, and associated with worse patient outcomes. Notably, following cessation of CSF-1R depletion, macrophages rebounded primarily to the SAM phenotype, which was associated with accelerated growth of mammary tumors. Our work reveals the spatial determinants of macrophage heterogeneity in breast cancer and highlights the disruption of macrophage self-renewal as a potential new therapeutic strategy.

## Introduction

Solid tumors evolve in complex ecosystems that promote consistent growth, neovascularization, and immune evasion. These processes are orchestrated through diverse dialogues between heterogeneous tumor cells and their supporting stroma, including fibroblasts and immune cells ^1,2^. Across human cancers, macrophages comprise the major immune cell component of the tumor microenvironment (TME) ^3^ and enhance cellular responses that promote invasive growth, angiogenesis, matrix remodeling, and immune tolerance ^4,5^. Consistent with their profound impact on tumor growth and dissemination ^6^, the accumulation of tumor-infiltrating macrophages is associated with therapeutic resistance and worse prognosis for cancer patients ^7–12^. These properties have highlighted macrophages as an attractive target for cancer therapeutics ^13^. While single-cell RNA-seq (scRNA-seq) studies have provided important in-depth transcriptional analyses of macrophage phenotypes and subpopulations with relevance to clinical outcomes and response to diverse therapies ^9,10,14–18^, mechanistic understanding of how macrophage phenotypes are determined in solid tumors remains incomplete. Therefore, deciphering the underlying mechanisms and cell-cell interactions that regulate macrophage phenotypes in tumors is critical for developing novel and effective interventions.

Macrophages encompass both recruited and tissue-resident myeloid cells that play central roles in immune defense and maintaining homeostasis during organogenesis and tissue remodeling ^19^. In different organs, macrophages display tissue-specific functions ^20–24^ which are mediated via unique transcriptional and epigenetic programs ^25–27^. Macrophage plasticity, a hallmark characteristic of the monocyte-macrophage lineage, mediates their functional diversification in steady-state, but also enables their rapid response to various insults such as pathogen infection ^28^ and tissue injury ^29,30^. Indeed, solid tumors are enriched in signals that are associated with tissue injury and wound healing responses, rendering the TME as ‘a wound that never heals’ ^31^. Consequently, tumor-infiltrating macrophages can be co-opted by these signals and undergo phenotypic alterations that resemble their phenotypes in injured tissues ^32–35^. This includes dampening inflammation to prevent excessive tissue damage ^29^, clearing cell debris, triggering matrix deposition by fibroblasts, and secreting growth factors that promote growth of endothelial and epithelial cells.

Tissue-resident macrophages arise during early developmental stages and self-renew locally with minimal contributions from adult hematopoiesis ^36–42^. Similarly, tissue-resident macrophages of mammary glands arise primarily from yolk-sac and fetal liver progenitors ^43,44^. However, under inflammatory conditions such as within mammary tumors, a substantial proportion of macrophages were found to originate from mobilization of bone marrow-derived monocytes via the CCL2-CCR2 axis ^6,45–47^. Nonetheless, whereas blocking monocyte recruitment (*Ccr2*-deficient mice) effectively reduced monocyte and macrophage numbers in healthy mammary glands ^44^, the abundance of macrophages in mammary tumors was not affected by this perturbation (*Ccr2-KO* MMTV-PyMT) ^45^. These data suggest that the long-term accumulation of macrophages in mammary tumors is primarily dependent on local self-renewal.

Macrophage proliferation, differentiation, and survival in healthy tissues rely on CSF-1R signaling through tissue-specific ligands IL-34 or CSF-1 ^48,49^. Specifically, mammary gland macrophages are dependent on CSF-1, as demonstrated using *Csf1*-deficient mice ^50^, while microglia and Langerhans cells can additionally rely on IL-34 ^51^. Upon differentiation, macrophages can lose their ability to proliferate in response to CSF-1R ligands ^52,53^. Indeed, in contrast to TME macrophages, tissue-resident macrophages in healthy mammary glands are predominantly quiescent ^44,54^. However, since CSF-1 is abundantly expressed in both healthy and malignant mammary tissues ^19,50,55^, it is unclear what specifically allows TME macrophages to regain their proliferation capability in response to the same mitogenic signal. These context-dependent responses imply that additional microenvironmental mechanisms (beyond CSF-1 sensing) are needed to license macrophage self-renewal in tumors and may serve as therapeutic vulnerabilities.

To interrogate the signals that shape breast cancer macrophage phenotypes and their ability to self-renew, we developed an *ex vivo* organotypic TME (oTME) model from murine mammary tumors ^56^. This model contains the major mammary TME constituents, including macrophages, tumor epithelial cells, and their supporting fibroblasts, while preserving the functions, transcriptional programs, cell-cell interactions, and typical spatial organization of mammary tumors. oTME interrogation of macrophage education and proliferation revealed that type-I interferon (IFN) signaling triggered macrophage self-renewal in a specific subset of *Ly6a*^+^F4/80^high^ macrophages. Surprisingly, and regardless of their ontogeny (tissue-resident vs. monocyte-derived), we found that macrophage self-renewal required cell contact, specifically with tumor-activated fibroblasts (cancer-associated fibroblasts; CAFs) but not with tumor epithelial cells, despite their comparable levels of CSF-1 expression. We found that Notch signaling, and specifically Notch4, was key to breast TME macrophage proliferation. Antibody-based neutralization of Notch4 inhibited macrophage proliferation and restrained tumor growth in an immunocompetent murine model, as well as lung metastasis from a patient-derived xenograft (PDX) model of triple-negative breast cancer (TNBC). We further characterized the spatial dependencies of macrophage heterogeneity using multi-omic spatial profiling ^57^ and found that spatial TME localization was the critical determinant of macrophage immunophenotypes, regardless of ontogeny. Specifically, whereas macrophage co-localization with tumor epithelial cells (Tumor-Nest Macrophages, TNMs) resulted in non-proliferative and pro-inflammatory phenotypes, interactions with stromal fibroblasts (Stroma-Associated Macrophages, SAMs) gave rise to proliferative, phagocytic, and immunosuppressive macrophages, which were resistant to CSF-1R depletion, and associated with poor clinical outcomes. Collectively our data showed that while ontogeny determines macrophage phenotypes under steady state, spatial signals in the TME override the ontogeny-derived differences between tissue-resident vs. monocyte-derived macrophages, leading to the accumulation of tumor-promoting, immunosuppressive macrophages.

## Results

### Organotypic TME (oTME) modeling of breast cancer macrophages identifies locally renewing populations

The phenotypic alterations in TME macrophages rely on dynamic interactions between macrophages and TME constituents, including tumor cells and fibroblasts ^1,58^. To enable comprehensive interrogations of the signals and interactions that fuel this transition, we developed an *ex vivo* TME platform from the MMTV-PyMT transgenic mouse model of mammary cancer ^59^ (hereafter referred to as organotypic TME; oTME) (**Fig. 1a**). The oTME model contains the major non-immune TME constituents of breast cancer, including tumor epithelial cells and fibroblasts, and preserves the typical structure of mammary tumors *in vivo* (**Fig. 1b and Extended Data Fig. 1a**).

**Figure 1:**
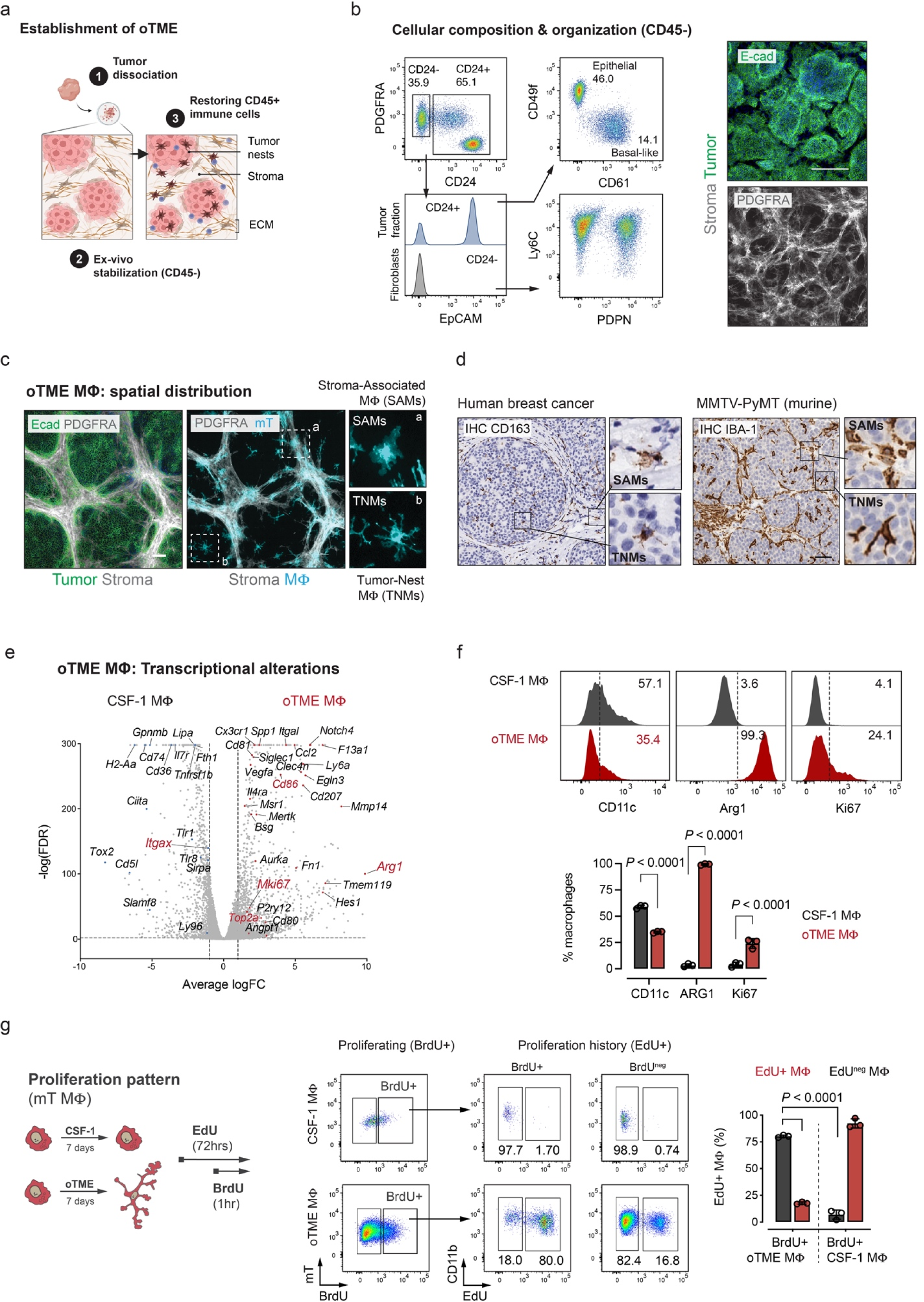
Breast organotypic tumor microenvironment (TME) recapitulates the cell-cell interactions and phenotypic alterations in breast cancer macrophages. (**a**) establishment of organotypic TME (oTME) from MMTV-PyMT breast tumor. ECM; extracellular matrix. (**b**) Left, the cellular composition and structure of oTME cultures. Flow cytometry and immunofluorescence analyses demonstrating the cell identities (CD45-) and proportions of oTME cells using mammary gland markers. Right, immunofluorescence for E-Cadherin (tumor) and PDGFRA (stroma) in two-week-old oTME cultures demonstrating the oTME cellular organization. Data are representative of at least 5 independent experiments. Scale bars, 100μm. (**c**) Spatial distribution of oTME Macrophages. Bone Marrow monocytes were purified from *Rosa26^mTmG^* mice (n=3 replicates) and plated with oTME cultures for 7 days and immunostained for E-Cadherin and PDGFRA. Scale bars, 100μm. (**d**) Immunohistochemistry (IHC) of breast cancer macrophages using CD163 (human) and IBA1 (murine) antibodies, demonstrating their typical spatial distribution in the TME. Scale bar, 100μm. (**e**) Volcano plot showing fold change (FC; x axis, log scale) and the FDR corrected *P*-value (y axis, -log10 scale) of expressed genes between CSF-1-treated BMDMs and BMDMs cultured with oTME cells for 7 days (n=3 replicates). Genes with an average count value above 3 in CSF-1- treated BMDMs are presented. Horizontal dotted line represents a *P*-value of 0.01 (-log10 scale). Selected genes are indicated by text. *P*-values of zero were set to the smallest non-zero *P*-value for visualization. (**f**) Immunophenotyping of oTME macrophages. BMDMs were cultured with oTME cells or CSF- 1 as control for 7 days and analyzed by flow cytometry with the indicated markers. Data shown as mean±SD, 2-tailed paired Student’s t-test. (**g**) EdU/BrdU dual pulse-chase labeling in CSF-1-treated and oTME macrophages. BMDMs cultured with oTME cells for 7 days, labeled with EdU for 72hrs, and then pulsed with BrdU for another 1hr. Macrophages were analyzed by flow cytometry for EdU/BrdU and quantified for single and double positivity (n=3 replicates). Data are shown as mean±SD, 2-way ANOVA test, Bonferroni-corrected.

First, we characterized the cellular identities of the oTME components. In murine mammary tissue, CD24 expression marks the luminal (CD24^high^) and basal (CD24^int^) epithelial cells, while CD24^neg^ cells are identified as the stromal fraction in healthy and malignant tissues ^60,61^. Relying on these established markers, we identified two populations of CD24+ tumor cells; a dominant population of tumor epithelial cells (CD24^high^EpCAM^+^) that express luminal markers ^62,63^, and a smaller population of EpCAM^neg^ tumor cells with basal-like markers (CD49f^low^CD61^+^) ^64,65^ (**Fig. 1b**). Within the CD24^neg^ stromal fraction, we identified the extracellular matrix (ECM)-depositing PDPN^+^ subset observed in murine and human breast cancer ^58,66,67^ (**Extended Data Fig. 1b,c**). Notably, we verified that only EpCAM^+^ tumor cells, but not the CD24^neg^ stromal cells, displayed characteristics of transformed cells (**Extended Data Fig. 1d,e**).

Next, we introduced primary macrophages into the system and examined their spatial organization and the associated transcriptional alterations. To enable visualization, we utilized bone marrow (BM) Ly6C^high^ monocytes purified from *Rosa26^mTmG^*mice ^68^ that express membrane-tagged tdTomato (mT). Flow cytometry of monocyte-derived oTME macrophages confirmed the conversion of Ly6C^high^F4/80^neg^ monocytes into Ly6C^neg^F4/80^+^ macrophages (**Extended Data Fig. 1f, Supplementary Movie 1**), indicating that monocytes in this model differentiate into mature macrophages rather than into Ly6C^+^F4/80^+^ monocytic myeloid-derived suppressor cells (M-MDSCs) ^69^. Immunofluorescence staining of oTME macrophages (mT^+^) along with epithelial (E-Cadherin) and stromal (PDGFRA) markers (**Fig. 1c**) revealed two major distributions; a dominant subset that co-localized with PDGFRA^+^ stromal regions (Stroma-Associated Macrophages; SAMs) and a smaller subset that localized within the tumor nests (Tumor-Nest Macrophages; TNMs). Notably, oTME macrophage localization recapitulated the typical spatial distributions of macrophages in murine and human breast cancer (**Fig. 1d**) ^70^.

We then assessed the transcriptional alterations (TME-education) in oTME macrophages via bulk RNA-seq of bone marrow-derived macrophages (BMDMs) cultured for ten days in oTME by comparison to CSF-1-treated BMDMs as control. Differential expression analysis (**Fig. 1e; Supplementary Table 1**) revealed downregulation of immune-stimulatory mediators (*Tnfrsf1b, Tlr1, Tlr8, Tlr7*) ^71^, coupled with robust induction of genes associated with wound healing responses ^35^ including angiogenesis (*Vefga, Angpt1, Egln3*), Notch signaling (*Notch4, Hes1*), immune modulation (*Arg1, Il4ra, Ccl2, Cd86, Cd80, Cd207*), cell cycle (*Mki67*), scavenging (*MerTK, Msr1, Tyro3*), and tissue remodeling (*Spp1, Fn1, Bsg, Bgn, Mmp14, Timp3*). The transcriptional alterations were functionally validated by flow cytometry, T cell suppression assay, and LPS/IFNγ challenge (**Fig. 1f and Extended Data Fig. 1g,h**). Following 10 days of TME education, macrophages transitioned from a quiescent (Ki67^neg^CD11c^high^Arg1^neg^) state into proliferative and immunosuppressive Ki67^+^CD11c^low^Arg1^+^ phenotype associated with wound healing response ^32,72,73^. Notably, while a major proportion of oTME macrophages became proliferative (24.1%, *P*= 0.0072; **Fig. 1f**), the vast majority (>95%) of CSF-1-treated macrophages remained predominantly quiescent despite excessive exposure to CSF-1 (20ng/mL every other day), the key mitogenic signal of macrophages ^23,74^.

To distinguish between broad proliferative capacity vs. persistent proliferation within a specific subset, we tracked the proliferation history of macrophages during the education process. We utilized a dual pulse-chase labeling strategy with EdU and BrdU that enables cell cycle scoring of the same cells at two different time points (**Fig. 1g**). In agreement with the quiescent phenotype in terminally-differentiated macrophages ^52,75^, the majority of the BrdU^+^ control macrophages displayed a low proliferative history (EdU^+^; 6.9%±4.3% mean±SD). In contrast, the BrdU^+^ cycling subset of oTME macrophages displayed a substantial proliferative history (EdU^+^; 80.4%±1.1%), while the non-cycling subset (BrdU^neg^) showed lower proliferative history (EdU^+^; 17.7%±1.0%) (**Fig. 1g**). Thus, the formation of continuously proliferating and non-proliferating subsets indicates that macrophage proliferation in tumors has characteristics of self-renewal. Importantly, the signals that were required for macrophage self-renewal (beyond CSF-1 availability) were successfully captured by the oTME model, enabling mechanistic interrogation of this clinically-relevant phenotype ^12^.

### scRNA-seq time course of macrophage education revealed a self-renewing subset instigated by type-I interferons

To characterize the establishment of proliferative macrophages in tumors, we performed an education time-course experiment using single-cell RNA-seq (scRNA-seq). We compared macrophages educated in the oTME for two days (“Early”) and ten days (“Late”) along with CSF- 1-treated macrophages as controls (“CSF-1 MΦ”) (**Fig. 2a, Extended Data Fig. 2a; Supplementary Table 2**). To survey the transcriptional dynamics of macrophage education we reconstructed an ‘education trajectory’ by overlaying the single-cell macrophage transcriptomes of the CSF-1-treated (at day 2 and day 10), “Early”, and “Late” groups onto a pseudo-temporal axis (**Fig. 2b, Extended Data Fig. 2b; Supplementary Table 4, Methods**). This analysis revealed a dynamic process consisting of heterogeneous activation states (**Fig. 2c**). The education process at day 2 was characterized by down-regulation of genes that mediate immunostimulatory responses, including GM-CSF/IL-5 and TNF signaling (*Csf2rb, Tnfrsf1b, Tnfrsf4*) ^71,76^ and negative regulators of phagocytosis (*Sirpa*). Interestingly, these changes were coupled with a robust induction of type-I interferon genes (*Irf7, Isg15, Rtp4, Ifit1, Ccl2, Cxcl10, Ccl9*) ^77,78^ in conjunction with cell cycle activation (*Mki67, Top2a*) (**Fig. 2d**). As the education process continued (at day 10), macrophages transitioned into the wound-healing phenotype, upregulating genes associated with tissue injury inflammation (*Bsg, Bgn, Ninj1, Mmp14, Fn1, Ecm1*), angiogenesis (*Egln3, Vegfa, Eno1*), and immune modulation (*Arg1, Thbs1, Mrc1, Apoe, Ptgs1*) (**Fig. 2d**).

**Figure 2:**
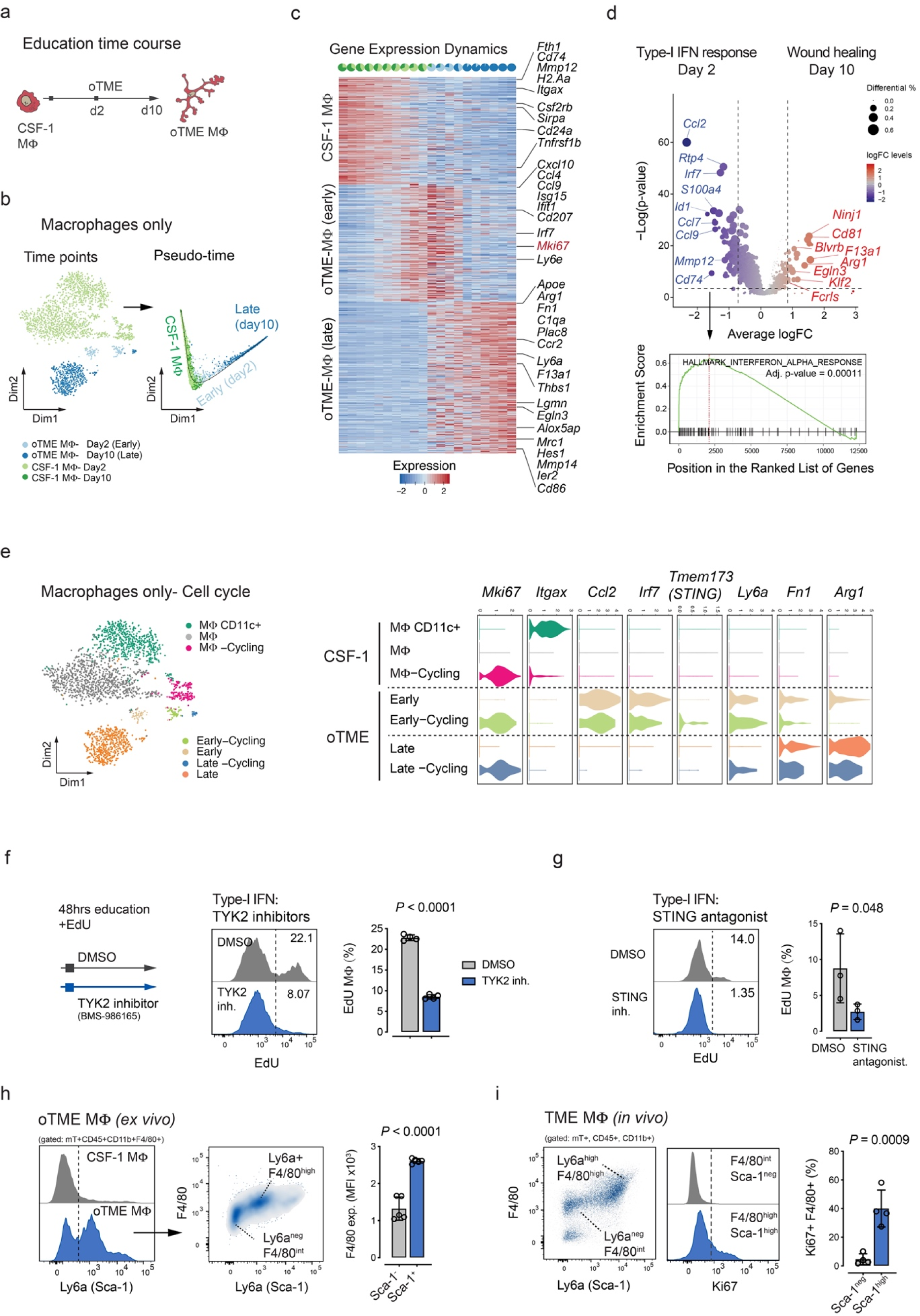
Single-cell RNA-seq education time course revealed a subset of self-renewing macrophages triggered by type-I IFNs. (**a**) Study design: BMDMs were cultured with oTME cells for 2 and 10 days or left unperturbed with CSF-1 as control. Cells were then collected and profiled using the 10x Chromium single-cell RNA-seq platform. (**b**) Two-dimensional t-SNE plot (left; Dim1 and Dim2) of single-cell macrophage transcriptomes (excluding CSF-1-treated day 10) and two-dimensional education trajectory (right; Dim1 and Dim2) with inferred pseudo-time axis (black line). (**c**) Heatmap showing macrophage education expression dynamics of CSF-1-treated (n=214), early (day 2, n=235), late (day 10, n=301) genes. The cells are sorted and split into 20 equal-width bins by the education pseudo-time, average expression level (row-wise Z-score; color scale) of education signature genes (rows) across each bin (columns) is shown. (**d**) Volcano plot showing log fold change (logFC) of genes (n=12,322) between day 2 (early) vs. day 10 educated macrophages (late) and their significance (y axis; -log10 scale). Genes are dotted and colored by logFC levels (color scale). The size of each dot represents the difference in the fraction of detection between the two groups. *P*-values were determined by Wilcoxon Rank Sum test. Vertical dotted lines represent ±0.75 logFC. Horizontal dotted lines represent a *P*-value of 0.01 (-log10 scale). Raw and FDR corrected *P*-values and logFC values are listed in Supplementary **Table 5**. Gene Set Enrichment Analysis in day 2 macrophages shows a significant enrichment of Interferon alpha response (FDR < 0.05). (**e**) Left, two-dimensional t-SNE plots (Dim1 and Dim2) of annotated single-cell transcriptome clusters colored by cell cycle states. Right, violin plots showing selected genes from single-cell transcriptomes of CSF-1-treated, early, and late educated macrophages. (**f**) BMDMs were plated with oTME cells for 48hrs in the presence of EdU (10µM) and TYK2 inhibitor (BMS-986165; 10µM) or DMSO as control. EdU incorporation was analyzed by flow cytometry (n=4 replicates). Data are shown as mean±SD, 2-tailed unpaired Student’s t-test. (**g**) BMDMs were plated with oTME cells for 7 days, and then treated for 48hrs with EdU and STING inhibitor (C-157; 1µM) or DMSO as control. EdU incorporation was analyzed by flow cytometry (n=4 replicates). Data are shown as mean±SD, 2-tailed unpaired Student’s t-test. (**h**) Induction and expression of *Ly6a* (Sca-1) in oTME macrophages. Flow cytometry for *Ly6a* (Sca-1) and F4/80 in BMDMs treated either with CSF-1 or cultured with oTME cells for 10 days (n=5 replicates). Data are shown as mean±SD, 2-tailed unpaired Student’s t-test. MFI; mean fluorescence intensity. (**i**) Relative expression of *Ly6a* (Sca-1) and Ki67 in TME macrophages. Flow cytometry plots quantifying the expressions of *Ly6a* (Sca-1) in F4/80^high^ vs. F4/80^int^ macrophages (left) and Ki67 Sca-1^high^ vs. Sca-1^neg^ macrophages from PyMT tumor transplants (n=4 replicates). Data are shown as mean±SD, 2-tailed unpaired Student’s t-test.

To specifically focus on the proliferative subsets, we re-annotated the macrophage transcriptomes according to their (i) education state and (ii) cell cycle status (**Fig. 2e**). Consistent with the nonproliferative state in terminally differentiated macrophages ^52^, tumor-naive macrophages remained refractory to CSF-1 signals and showed neither induction nor changes in cycling proportions at either time-point (**Extended Data Fig. 2c**). In contrast, nearly half (42.8%) of the oTME macrophages at day 2 engaged the cell cycle (*Mki67*+) and a subset (4.7%) remained proliferative at day 10, which was further verified by EdU labeling (**Extended Data Fig. 2d**). To uncover potential cell-cycle activation mechanisms, we identified genes that were uniquely or highly expressed in the cycling subsets. This includes *Ly6a* (Stem Cell Antigen-1; Sca-1) and *Tmem173* (STING) (**Fig. 2e**), a sensor of tumor-derived cGAMP/dsDNA and a potent stimulator of type-I IFNs in macrophages ^79,80^. Notably, acute activation of type-I IFNs and *Ly6a* (Sca-1) induction were previously shown to mediate activation of self-renewal in dormant hematopoietic stem cells (HSCs) ^81–83^. To examine whether induction of type-I IFN and cell cycle activation are also functionally-dependent processes in breast cancer macrophages, we measured EdU incorporation in oTME macrophages during the early stages of education (the first 48 hrs) while blocking type-I IFN signaling using TYK2 inhibitors (BMS-986165; pan inhibition) ^84,85^ or STING-specific (C-157) inhibitors at later stages of education (**Fig. 2f,g**). The reduction in EdU+ incorporation suggests that type-I IFN signaling and cell cycle were indeed functionally-dependent processes in TME macrophages.

To characterize Sca-1 expression in breast cancer macrophages, we used the oTME *(ex vivo)* and tumor transplants *(in vivo)* models. We noted that Sca-1 consistently marked a subpopulation of F4/80^high^ macrophages and was specifically enriched in the Ki67+ signal (**Fig. 2h,i**). These results corroborated the scRNAseq results and confirmed that F4/80^high^Sca-1^+^ expression highlights the proliferating subset in breast cancer macrophages in response to type-I IFN signaling activation. Notably, stimulation of tumor-naive macrophages with IFNα and CSF-1 did not lead to robust proliferation (**Extended Data Fig. 2e**), suggesting that type-I IFN signaling was necessary but not sufficient to license macrophage proliferation, and additional oTME signals were required.

### Macrophage proliferation in mammary tumors is mediated by fibroblasts activated by tumor cells

Next, we leveraged our platform to identify the specific interactions that promoted the Sca-1+F4/80^high^ proliferative phenotype. Immunostaining for F4/80 in oTME macrophages (mT) along with FN1 (**Fig. 3a**) revealed two populations with intermediate (F4/80^int^) and high (F4/80^high^) expression with distinct spatial contexts. Specifically, while F4/80^int^ macrophages localized inside tumor epithelial nests (TNMs), the F4/80^high^ macrophages were localized exclusively in FN1^+^ stromal regions (SAMs) (**Fig. 3a**), indicating that the self-renewing subset resides in tumor stroma.

**Figure 3:**
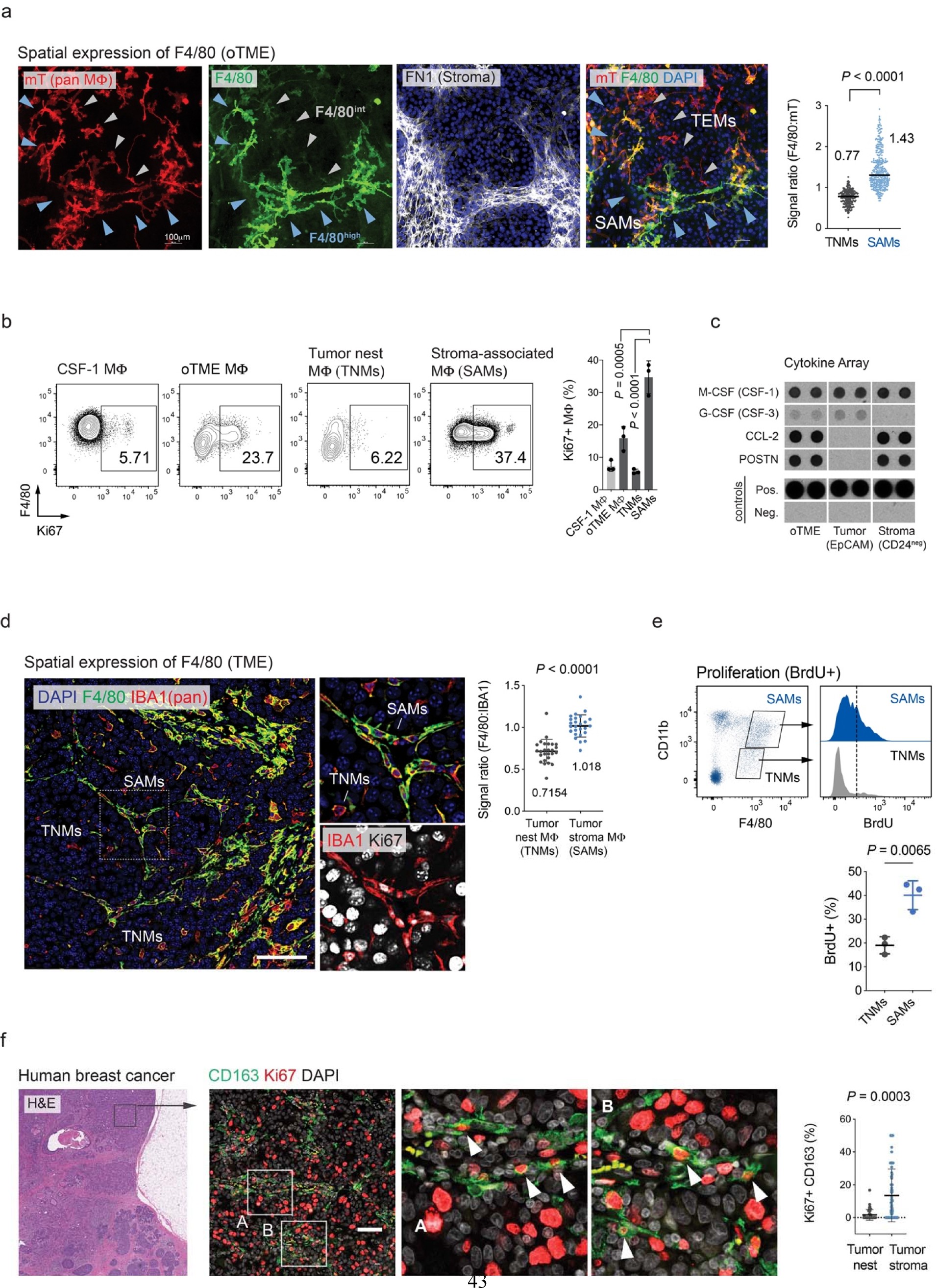
Stroma-associated macrophages are the self-renewing subset in breast cancer. (**a**) Immunofluorescence of endogenous tdTomato (mT), F4/80, and Fibronectin (FN1) in *Rosa26^mTmG^* monocyte-derived macrophages that were cultured with oTME cells for 10 days and immunostained as indicated (n=3 replicates). Scale bars, 100μm. Fluorescent signal ratios of F4/80 over tdTomato were plotted as a function of physical localization with FN1+ cells (n=282 cells). Data are shown as mean±SD, Mann-Whitney test. (**b**) Flow cytometry of Ki67 in BMDMs cultured either with oTME, tumor epithelial, or stromal cells, or CSF-1 as control cells (10ng/mL every other day). Data quantification as % for Ki67 are presented (n=3 replicates). Data are shown as mean ±SD, one-way ANOVA test, Bonferroni-corrected. (**c**) Cytokine array analysis of supernatants collected from purified tumor epithelial, stromal or unsorted oTME cells. Growth media were conditioned for 5 days from confluent cultures and probed for 111 cytokines. Selected cytokines are presented along with positive and negative spots (see **Extended Data Fig. 4** for full array). (**d**) IF for IBA1 (red), F4/80 (green), and Ki67 (white) in mammary tumors. Dashed lines mark tumor nests. Fluorescent signal ratios of F4/80 over IBA1 were recorded and plotted as a function of their spatial localization (tumor stroma vs. tumor nests). Dots represent fields of view (n=24) from 4 tumor macrophages. F4/80^int^ TNMs and F4/80^high^ SAMs are labeled. Scale bar, 100μm. Data are shown as mean±SD, two-tailed paired non-parametric Wilcoxon test. Scale bar, 50μm. (**e**) *In vivo* BrdU incorporation experiment in tumor-infiltrated macrophages. Tumor-bearing mice (n=3) were pulsed with BrdU (2mg/mouse/intraperitoneally) 24hrs before tumor harvesting and analyzed by flow cytometry. Data are shown as mean±SD, 2-tailed unpaired Student’s t-test. (**f**) IF for CD163 and Ki67 in human breast tumors. Scale bar, 100μm. Arrows indicate Ki67 positive SAMs. Data (fields of view; tumor nests n=39, tumor stroma n=88) are shown as mean±SD, Mann–Whitney U test. Dots represent fields of view from 5 cases.

To directly connect macrophage self-renewal with stromal interaction, we measured Ki67 in macrophages exposed to either tumor epithelial-enriched (EpCAM^+^) or stroma-enriched (CD24^neg^PDGFRA^+^) cultures. To address the generalizability of these interaction-specific responses, we challenged macrophages from various sources, including tissue-resident macrophages harvested from healthy mammary glands, mammary tumor macrophages, and BMDMs (**Fig. 3b, Extended Data Fig. 3a**). We found that regardless of their origin, macrophage proliferation was significantly enhanced by the presence of stromal cells (CD24^neg^PDGFRA^+^) and not by tumor epithelial cells, or soluble CSF-1. Since potential differences in CSF-1 expression levels in stromal vs. tumor epithelial cells could underlie these differences, we also compared secretomes from purified EpCAM+ and CD24^neg^PDGFRA+ cultures (**Fig. 3c** and **Extended Data Fig. 3b**). Interestingly, while several cytokines were lineage-specific, we found that both cell types expressed comparable levels of CSF-1, the key mitogenic signal of macrophages. These results indicate that even within the TME context, specific signals from stromal cells were needed to license proliferation in macrophages.

Since macrophages in healthy mammary glands are predominantly quiescent despite their close interactions with fibroblasts, we further examined the interplay between mammary gland fibroblasts and macrophages (both harvested from healthy glands) (**Extended Data Fig. 3c**). Notably, while tumor-naive fibroblasts supported macrophage survival ^74^, they failed to promote macrophage proliferation, unless activated by the presence of tumor cells which promote their transition into CAFs (POSTN+, IL-6) (**Extended Data Fig. 3d,e**), and despite comparable levels of CSF-1 (**Extended Data Fig. 3c,4**). This interplay portrays a three-way interaction, where fibroblasts can license macrophage proliferation only following their activation by tumor cells.

To experimentally test the ability of fibroblasts to drive macrophage self-renewal *in vivo*, we employed orthotopic transplantations of tumor epithelial cells alone or enriched with PDGFRA^+^CD24^neg^ stromal cells. We confirmed the stromal enrichment by IHC (Vimentin) and compared the myeloid landscapes in the resulting tumors by flow cytometry (**Extended Data Fig. 5a,b**). Beyond the increase in macrophage infiltration (IBA1, IHC staining), higher fibroblast abundance in tumors also triggered a prominent shift in macrophage immunophenotype, without affecting the M-MDSC population (**Extended Data Fig. 5c**). Specifically, as predicted by our oTME results, the F4/80^high^CD11b^high^ subset became the dominant population in fibroblast-enriched tumors (expanding from 42.3% to 74.9%, *P*= 0.0005) while the F4/80^int^CD11b^low^ population was markedly reduced from 41.9% to 3.6% (*P*= 0.0023). These results further validate that the F4/80^high^CD11b^high^ and F4/80^int^CD11b^low^ immunophenotypes correspond to specific interactions with fibroblast (SAMs) and tumor epithelial cells (TNMs), respectively. We also confirmed these proliferative biases in mammary tumors by BrdU labeling and IHC for Ki67 (**Fig. 3d**). Consistent with the *ex vivo* oTME data, we found higher rates of BrdU incorporation in the SAM subset (F4/80^high^CD11b^high^), as well as higher F4/80 expression and Ki67 positivity in stromal macrophages by IHC (**Fig. 3e**). Of note, macrophage proliferation in the MMTV-PyMT model was detected in both dysplasia and proliferating adenocarcinoma regions (**Extended Data Fig. 6a,b**), suggesting that macrophage self-renewal occurs throughout the different stages of neoplastic growth of mammary tumors.

Finally, we validated these findings in primary human samples from grade 3 invasive ductal carcinomas across HER2^+^, ER/PR^+^, and triple-negative (TNBC) molecular subtypes (**Supplementary Table 5;** n=15 patients). In agreement with the murine models, we found that the Ki67^+^ CD163^+^ macrophages were preferentially detected in the stromal regions in comparison to tumor nests (**Fig. 3f**). Collectively, these results demonstrate that macrophage accumulation in breast cancer is a spatially-determined process, driven by interaction with CAFs.

### Macrophage self-renewal is contact-dependent and mediated by Notch4

The ability of TME macrophages to self-renew suggests that rather than blocking monocyte recruitment, specific targeting of macrophage proliferation is required to limit their long-term accumulation ^6^. Using the oTME, *in vivo,* and human data, we demonstrated that macrophage proliferation is supported by close interaction with tumor-activated fibroblasts (CAFs), while soluble CSF-1 was not a differentiating factor. To determine whether cell contact is required, we compared EdU incorporation in macrophages that were plated either directly with oTME cultures or physically separated by transwells but permit cytokine diffusion. We found that macrophage proliferation was enhanced only when physical contact was allowed (**Fig. 4a**), highlighting pathways that rely on cell contact as potential regulators.

**Figure 4:**
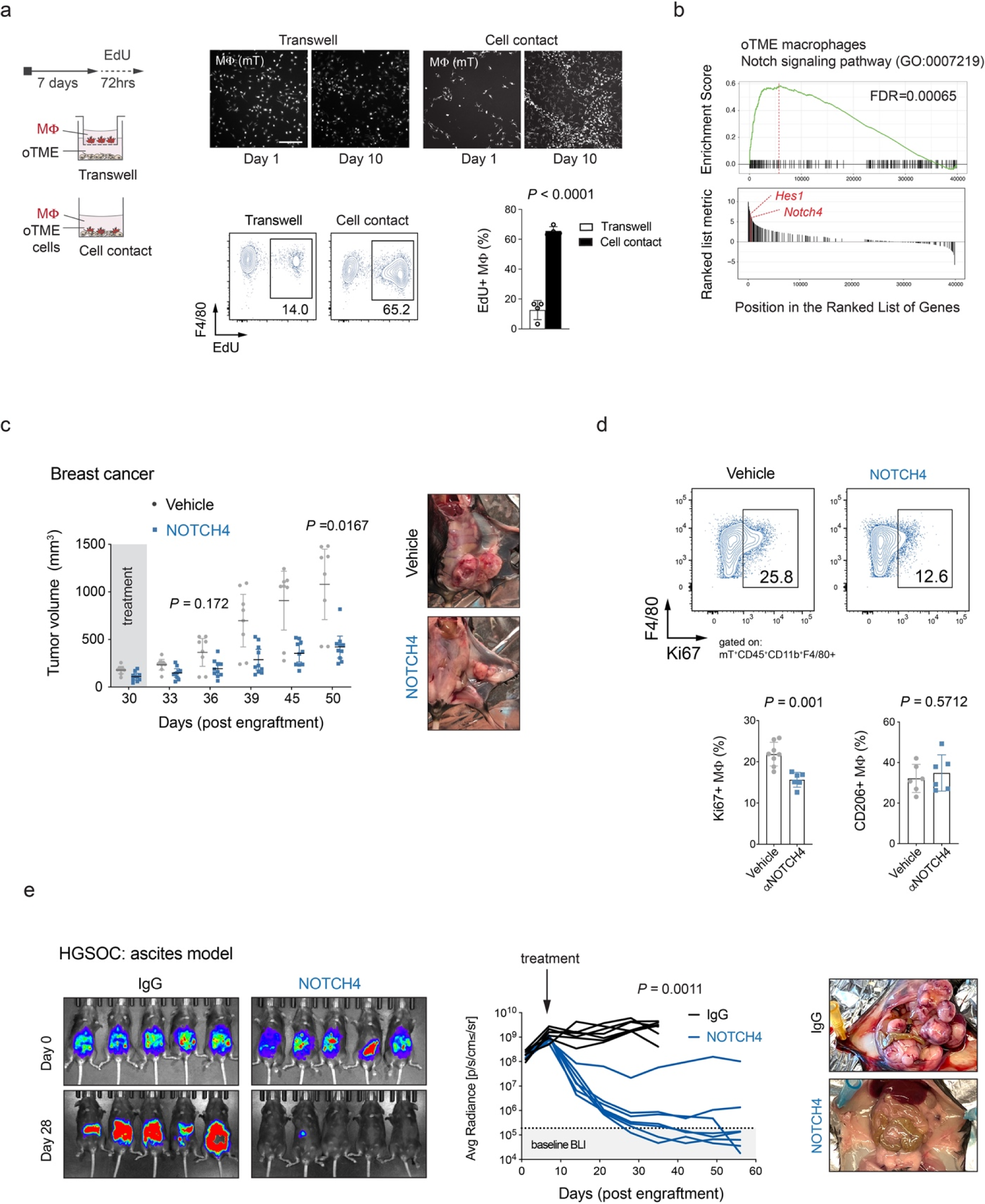
Macrophage self-renewal in mammary tumors is contact-dependent and essential for tumor growth. (**a**) Macrophages plated either directly with oTME cells (cell contact) or separated by transwells for 7 days, followed by 72 hrs of EdU labeling. Macrophages were imaged (mT) at plating and prior to flow cytometry analysis of EdU. Individual dots represent individual fields of view from 3 independent experiments. Data are shown as mean±SD, 2-tailed t-test. Scale bar, 100μm. (**b**) Gene Set Enrichment Analysis of Gene Ontology showing a significant enrichment of Notch signaling pathway in oTME macrophages (GO:0007219; FDR=0.00065; top, enrichment scores; bottom, rank positions of member genes). (**c**) *Rosa26^mTmG^* mice (n=8) with established mammary tumors were treated with Notch4 monoclonal antibodies (15μg/kg body weight, dosed every 3 days, intraperitoneally) or vehicle (PBS) as a control for 20 days. Tumor sizes were recorded by a caliper and shown as mean ±SD, 2-way ANOVA test, Bonferroni-corrected. At the trial endpoint, mammary tumors were imaged (right). (**d**) Macrophage proliferation analysis in mammary tumors from (**c**) following Notch4 treatment (n=6) at study endpoint. Tumors were harvested and scored for Ki67 and CD206 expression in macrophages by flow cytometry. Data are shown as mean±SD, 2-way ANOVA test, Bonferroni-corrected. Macrophage abundance quantified as percent of live CD45+ cells. Data are shown as mean±SD, Mann-Whitney test. (**e**) C57BL/6 female mice were injected with luciferase-expressing HGSOC cells (1 x 10^6^, intraperitoneally) and treated 7 days after tumor BLI signals were established with NOTCH4 or isotype control IgG antibodies (15μg/kg body weight, dosed every 3 days, intraperitoneally). BLI measurements were recorded every week and representative images are presented.

Gene set enrichment analysis of RNA-seq data from oTME macrophages showed significant enrichment of the Notch pathway (GO:0007219, FDR=0.00065) and upregulation of several Notch signaling mediators, including *Notch4*, *Dll1*, *Hes1*, *Rbpj*, *Adam17*, and *Adam10* (**Fig. 4b and 1e; bulk RNAseq**). A previous study has shown that pan-ablation of Notch signaling in breast cancer macrophages (*Cd11c*^Cre^*Rbpj*^flox^ MMTV-PyMT) increased the tumoricidal activity of tumor-infiltrating T cells ^45^. To connect Notch signaling directly with macrophage proliferation in breast cancer, we first confirmed activated Notch signaling in cell lysates from sorted oTME macrophages by probing for the cleaved portion of Notch receptor (NICD) (**Extended Data Fig. 7a**). Then, we examined the impact of Notch inhibition on macrophage proliferation following treatments with two protease inhibitors that regulate Notch activation in receiver cells: (i) γ- secretase inhibitors (Compound-E) that inhibit the intracellular cleavage of activated Notch receptors (**Extended Data Fig. 7b**) and (ii) Adam17 inhibitors (A17Pro) (**Extended Data Fig. 7c**) that inhibits the extracellular cleavage of ligand-activated Notch receptors ^86,87^. Notably, both treatments resulted in effective inhibition of macrophage proliferation, indicating that Notch signaling was required to maintain the proliferative state in TME macrophages.

Among the upregulated Notch-related genes in oTME macrophages, we identified *Notch4* receptor among the top induced genes (**Fig. 4b and 1f; bulk RNAseq**). Therefore, we examined the therapeutic potential of Notch4 neutralization on macrophage proliferation and tumor growth *in vivo*. Tumor cells were engrafted in the mammary gland of *Rosa26^mTmG^* mice to allow clear identification of host cells (mT-labeled), and tumors were allowed to establish (∼100-150 mm^3^) before treatment was initiated (**Fig. 4c**). Treatment with anti-Notch4 monoclonal antibodies (15mg/kg body weight, every 72 hrs, intraperitoneally) resulted in a substantial reduction in tumor growth (**Fig. 4c**), reflected in a 77.3% decrease in tumor volume over the experimental trial period (*P*= 0.0167, 2-way ANOVA test). Importantly, the impact on tumor growth was coupled with a significant reduction in Ki67+ macrophages (*P*= 0.001, Mann-Whitney test) (**Fig. 4d),** but without alteration of tumor vascularization (IF for CD31), or depletion of stromal cells (host; mT^+^CD45^neg^) (**Extended Data Fig. 7d,e**). Of note, Notch4 inhibitor-treated mice did not show adverse effects such as weight loss, as typically observed with γ-secretase inhibitor regimens ^88,89^. To further explore the potential macrophage-specific effect and translatability to human TNBC, we utilized patient-derived breast cancer cells from a TNBC patient (BR18 cells). We implanted luciferase-expressing BR18 cells in immunodeficient (NSG) mice and initiated treatment with anti-Notch4 or isotype control IgG antibodies following the same dosing regimen as in the murine breast cancer model. Despite the lack of functional T/B/NK cells in this model, we noted an attenuated growth of primary tumors at earlier time points and significant effect on lung dissemination at experimental endpoint (**Extended Data Fig. 7f**), indicating a direct effect on TME macrophages by Notch4 targeting. Moreover, the use of human cancer cells and anti-mouse Notch4 monoclonal antibodies excluded the possibility of a direct tumor targeting. To examine the generalizability of Notch4 blockade, we leveraged an aggressive model of high grade serous ovarian carcinoma (HGSOC; *Myc*^amp^:*Trp53*^KO^; termed MP cells) ^90^. Mice were challenged intraperitoneally with luciferase-expressing MP cells and treated when tumors were established. As for the breast cancer models, we observed a significant reduction of tumor growth (BLI signal) in the anti-Notch4-treated mice in comparison to the isotype control IgG-treated group (**Fig. 4e**). Collectively, these results demonstrate that macrophage proliferation in tumors is targetable by Notch4 inhibition.

### TME spatial localization determines the phenotypic heterogeneity in breast cancer macrophages

The close association between macrophage proliferation and spatial distribution suggests that local interactions may also manifest in distinct macrophage phenotypes. To address this potential impact, we differentiated monocytes in the presence of tumor nest epithelial cells (TNMs) or stromal cells (SAMs) and examined the morphological, functional, and immunomodulatory features in macrophages. *Ex vivo* imaging and immunophenotyping analyses revealed a significant functional diversification under these distinct conditions. Specifically, while TNMs were characterized by extensive branched dendrites (**Supplementary Movie 2**), immune stimulatory phenotype (CD11c) and lower scavenging potential (TIM-4^low^)^91^, SAMs were loaded with large phagosomes (**Supplementary Movie 3**), expressed immunosuppressive (CD206, PD-L1) markers, and displayed higher scavenging potential (TIM-4^high^) that was further validated following exposure to GFP-labeled T cells (**Fig. 5a,b, Extended Data Fig. 8a,b**). Importantly, similar phenotypic compartmentalization was detected in macrophages from murine tumors *in vivo* (**Fig. 5c,b**) and human breast tissues (**Fig. 5e**), demonstrating higher expression of immunosuppressive (PD-L1, CD206) mediators in SAMs vs. TNMs.

**Figure 5:**
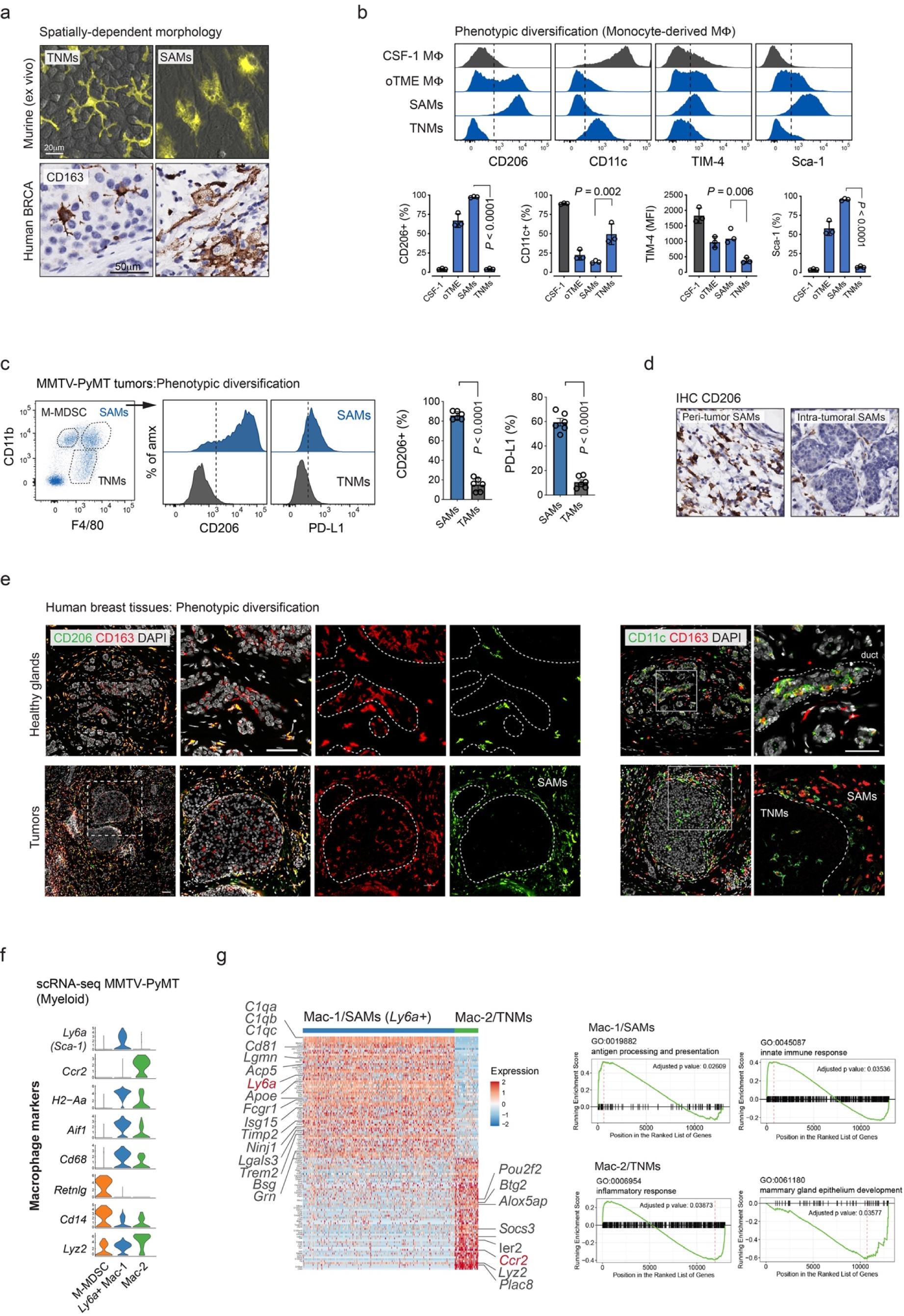
Macrophage heterogeneity is determined by spatial interactions with tumor epithelial cells and stromal fibroblasts. (**a**) Top panels: representative fluorescent images (mT in yellow) of BM monocytes from *Rosa26^mTmG^* mice (n=4; biological replicates) that were differentiated to macrophages for 7 days in the presence of either EpCAM+ tumor epithelial cells or PDGFRA+CD24^neg^ stromal cells. Scale bars, 50μm. **Bottom panels**: IHC images of CD163 macrophages in tumor nests (TNMs) vs. tumor stroma (SAMs), demonstrating the distinct morphology in human breast cancer macrophages as a function of their spatial localization. Scale bars, 20μm. (**b**) Immunophenotyping of monocyte-derived macrophages differentiated in the presence of the indicated cell type or with CSF-1 as control. BM monocytes were purified from *Rosa26^mTmG^* mice (n=3), cultured as indicated for seven days and analyzed by flow cytometry with the indicated surface markers. Data are shown as mean±SD, 2-way ANOVA test, Bonferroni-corrected. (**c**) Flow cytometry analysis of CD206 and PD-L1 expression in SAMs and TNMs from MMTV- PyMT tumors (n=6; biological replicates). Data are shown as mean±SD, 2-way ANOVA test, Bonferroni-corrected. (**d**) Tissue sections from (c) were analyzed by IHC for CD206, showing CD206+ macrophages only in peritumoral stroma and intratumoral stroma. (**e**) IF analysis for CD163/CD206 and CD163/CD11c in adjacent healthy and malignant mammary human tissues. Dashed lines highlight the normal (top) or malignant (bottom) epithelial cells (n=4 replicates). Scale bar, 100μm. (**f**) Violin plots showing expression of key marker genes of annotated myeloid cells from scRNA- seq analysis of MMTV-PyMT tumor fragments. (**g**) Differentially expressed genes in tumor infiltrating Mac-1 (*Ly6a*+ SAMs) and Mac-2 (*Ly6a*^neg^ TNMs). The expression level (row-wise Z-score; color scale) of signature genes (rows) is shown in the heatmap and their associated gene ontology (right).

Next, we asked whether the functional diversification of macrophages can be directly modulated by altering the cell-cell interactions. Monocyte-derived SAMs (F4/80^high^/CD206^+^) and TNMs (F4/80^int^/CD206^neg^) were sorted from oTME cultures and re-plated with the opposite microenvironments; SAMs with tumor cells, and TNMs with stromal cells (**Extended Data Fig. 8c**). Consistent with macrophage plasticity, the original recorded morphologies and immunophenotypes (PD-L1, CD206, F4/80) were overwritten according to the new microenvironment. To study this plasticity *in vivo*, we performed a macrophage re-population experiment in mammary tumors following macrophage depletion (anti-CSF-1R antibody) (**Extended Data Fig. 9a,b**). Tumor-bearing mice (*Rosa26^mTmG^*) were treated with anti-CSF-1R or IgG control for two weeks and then allowed to reconstitute the myeloid landscape for another two weeks, relying on the host’s monocytes and adoptively transferred GFP-labeled monocytes. Flow cytometry analysis of IgG-treated tumors showed that GFP+ monocytes were able to recapitulate similar phenotypes and subpopulations, indistinguishable from the host macrophages (**Extended Data Fig. 9c**). However, following macrophage depletion, both the host and GFP+ recruited monocytes gave rise primarily to the suppressive “SAM” phenotype (*P*= 0.0008, 2-way ANOVA test) that is associated with faster tumor growth (*P*= 0.003, 2-way ANOVA test) and substantial influx of tumor-infiltrating granulocytes (*P*= 0.0001, 2-way ANOVA test) (**Extended Data Fig. 9d**).

Lastly, to define the transcriptional signatures of these macrophage subsets *in vivo*, we performed single-cell RNA-seq on tumor fragments from the MMTV-PyMT model. Unsupervised clustering of CD45+ transcriptomes identified three distinct myeloid subpopulations (**Fig. 5f**), including M-MDSCs (*S100a8*, *S100a9*, *Cd14*^high^, *Retnlg*, *H2-Aa*^neg^) ^92,93^ and two macrophage subsets (*Cd68, Aif1, Lyz2, Cd14, H2-Aa*^+^) that we termed Mac-1 and Mac-2. Differential expression analysis (**Fig. 5g, Extended Data Fig. 8d, Supplementary Table 6**) revealed that Mac-1 macrophages expressed the SAM marker *Ly6a* (Sca-1) and genes that associated with complement activation (*C1qa, C1qb, C1qc*), active type-I IFNs signaling (*Isg15*), and cell scavenging (*Fcgr1, Trem2*). In contrast, the Mac-2 subset (*Ly6a*^neg^) was marked by expression of *Ccr2*, *Plac8*, *Pou2f2*, and *F13a1* and enriched for genes typical of pro-inflammatory responses (*Tlr2*, *Csf2ra*, *Csf2rb*, *Alox5ap*, *Ccl9*), and mammary gland epithelium development (**GSEA**). Furthermore, the proliferating subset (*Mki67*+) was observed with SAMs with *Ly6a* expression (**Extended Data Fig. 8e**) consistent with the Ki67^+^Sca-1^+^ self-renewing population.

### Integration of spatial transcriptomics and protein modalities validates the spatially-determined macrophage heterogeneity in breast cancer

To validate *in situ* the molecular signals underlying macrophage heterogeneity in their native environments, we performed spatial transcriptomics on murine breast cancer tissues integrated with protein markers (**Fig. 6a; SPOTS** ^57^). We designed a panel of 32 antibody-derived tags (ADTs; **Supplementary Table 7**) spanning key lineage and immune activation states including mammary tumor (EpCAM, KIT) and endothelial (CD31, TIE2) cells, fibroblasts (PDPN, PDGFRA), myeloid (CD45 CD11b, CD1d, F4/80, MHC-II, Ly-6G, Ly-6C), and lymphoid cells (CD19, B220, CD4, CD8a, NK1.1). The activation panel consisted of surface markers that are associated with immune stimulation (CD86, CD11c, CD27, CCR2, CD38, IgD, P2X7R, CD11a) and immune suppression (PD-1, PD-L1, CD25, TIM-3).

**Figure 6:**
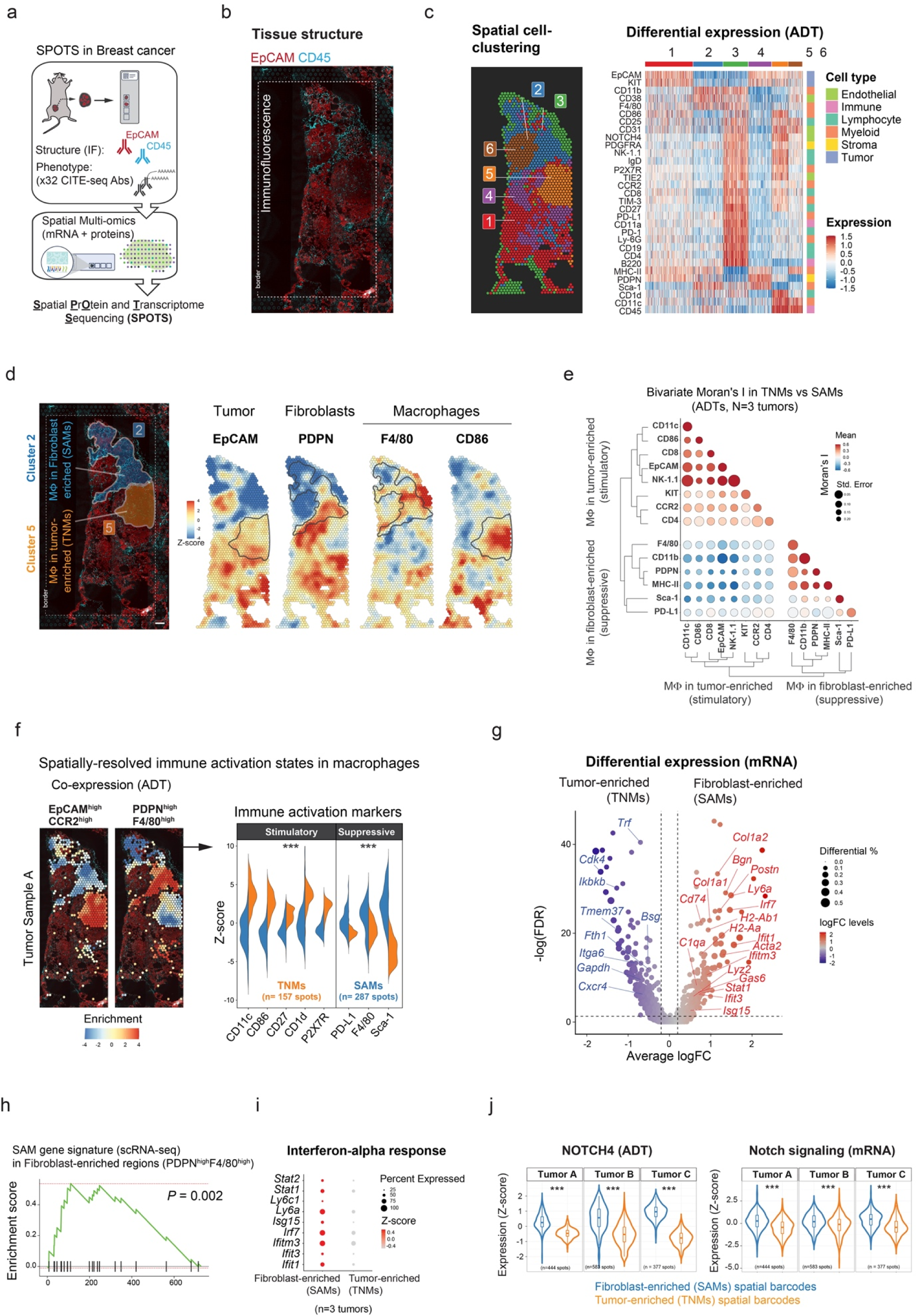
Transcriptional and immunological spatial profiling of macrophages in fibroblast-enriched vs. tumor-enriched regions in intact breast cancer tissue. (**a**) <u>S</u>patial <u>P</u>r<u>O</u>tein and <u>T</u>ranscriptome <u>S</u>equencing (SPOTS) on tissue from MMTV-PyMT model (n=3 mice). (**b**) Tissue structure. IF staining with EpCAM-PE and CD45-APC to reveal the tissue architecture. Scale bar, 200μm. (**c**) Spatial clustering and ADT signatures of each cluster. Left panel: spatially-informed clusters (1-6) overlaid onto tissue spatial barcodes. Right panel: heatmap of ADT expression for each cluster, where the right colorbar represents the cell-type annotation of each ADT. (**d**) Spatial ADT expression levels of key surface markers for tumor (EpCAM), fibroblasts (PDPN), and macrophages (F4/80, CD86) in fibroblast-enriched region (cluster 2) and adenocarcinoma (cluster 5). Note enrichment of EpCAM^high^CD86^high^ expression in adenocarcinoma region (cluster 5; consistent with TNMs) vs. enrichment of PDPN^high^F4/80^high^ expression in fibroblasts-enriched regions (cluster 2; consistent with SAMs). (**e**) Spatial correlation of lineage (NK1.1, CD4, CD8, CD11b, F4/80, MHC-II, EpCAM) immunostimulatory (CD27, CD86, CCR2, CD11c), and immunosuppressive (PD-L1, Sca-1) ADTs in fibroblast-enriched region (cluster 2) and adenocarcinoma region (cluster 5). ADTs are colored by bivariate Moran’s I (color scale). The size of each dot represents the inverse of standard error of the mean (S.E.M; n=3 mice). Dendrograms indicate the hierarchical clustering of the ADTs. (**f**) EpCAM and CCR2 vs. PDPN and F4/80 ADTs co-expression levels overlaid onto tumor sample A tissue (Methods) and immunophenotyping of SAMs and TNMs. Middle panel: boxplots of EpCAM and CCR2 vs. PDPN and F4/80 co-expression levels (addition of EpCAM and CCR2, or PDPN and F4/80 expression values). Right panel: violin plots of immune-stimulatory and suppressive ADT expression levels. Kolmogorov–Smirnov test, \**P* < 0.05, \*\**P* < 0.01, \*\*\**P* < 0.001, otherwise not significant (n.s). Each boxplot ranges from the first and third quartiles with median values shown as middle lines, and the whiskers represent 1.5 times the interquartile range. (**g**) Volcano plot showing log fold changes (logFC) of top 5,000 most variable genes between fibroblast-enriched region (cluster 2) vs. adenocarcinoma region (cluster 5) and their significance (y axis; -log10 scale). Genes are dotted and colored by logFC levels (color scale). The size of each dot represents the difference in the fraction of detection between the two groups. Macrophage-related genes are annotated. *P*-values were determined by Wilcoxon Rank Sum test. Vertical dotted lines represent ±0.2 logFC. Horizontal dotted lines represent FDR of 0.05 (-log10 scale). Raw, FDR corrected *P*-values, and logFC values are listed in Supplementary Table 10. (**h**) Gene Set Enrichment Analysis of scRNA-seq *Ly6a*+SAM gene signature (Supplementary Table 7; Fig. 5g) in the fibroblast-enriched region (cluster 2) (*P*= 0.002). (**i**) Gene expression of IFNα response genes (*Ly6a, Ly6c1, Ifit1, Ifit3, Ifitm3, Irf7, Isg15, Stat1, Stat2*) in SAMs (cluster 2) and TNMs (cluster 5). Genes are dotted and colored by expression levels. The size of each dot represents the percentage of expression in the tissue area. (**j**) Notch signaling pathway activity in fibroblast-enriched regions (SAMs) and adenocarcinoma regions (TNMs) across all three biological replicates. Left panel: violin plots of NOTCH4 ADT expression levels. Right panel: violin plots of Notch signaling pathway (GO:0007219; n=179 genes) transcriptional expression levels. Kolmogorov–Smirnov test, \**P* < 0.05, \*\**P* < 0.01, \*\*\**P* < 0.001, otherwise not significant (n.s).

We applied a spatially-informed clustering approach ^94,95^ (**Supplementary Table 8**) and identified six spatial regions with distinct histopathological, transcriptional and protein expression characteristics (**Fig. 6b,c, Extended Data Fig. 10a-c; Supplementary Table 9**). The peri-tumor regions (cluster 3) were enriched for adipocytes (*Adipoq*, *Cd36*), endothelial (CD31), and immune cells (CD8, CD4, B220, NK1.1, F4/80). Phenotypically, these regions co-expressed immune activation (CD38, CD27, CCR2, CD1d, CD11a) and immune exhaustion (PD-L1, PD1, CD25, TIM-3) markers, reflecting the immunosuppressive barrier formed around solid tumors ^57^. The remaining cluster regions (clusters 1, 2, 4, 5, and 6) corresponded to the varying histopathological stages of mammary gland transformation ^56^ (**Extended Data Fig. 10d**).

We focused on regions 2 and 5 as they displayed opposing cellular characteristics with regards to fibroblast infiltration (**Fig. 6d, Extended Data Fig. 11**). Region 2 was enriched with fibroblasts (PDPN, PDGFRA) and myeloid (Sca-1^+^, MHC-II, CD11b^high^, F4/80^high^) markers consistent with the *Ly6a^+^*SAMs immunophenotype. In contrast, region 5 displayed low fibroblast abundance and alternative expression of myeloid markers consistent with *Ccr2*^+^ TNMs (CCR2^+^, MHC-II, CD11b^low^, F4/80^low^). Furthermore, while tumor epithelial-enriched spatial barcodes positively correlated with myeloid immune-stimulatory markers (CD11c^high^, CD86^high^, CD1d) and higher lymphocyte infiltration (CD8, CD4, NK1.1), the fibroblast-enriched spatial barcodes expressed immune-suppressive markers of macrophages (PD-L1, Sca-1) and were almost devoid of lymphocyte-related genes (**Fig. 6e,f, Extended Data Fig. 12a**). These inverse correlations were consistent with TNMs and SAMs representing immune activating and suppressive macrophage subsets, respectively. Finally, differential expression analysis between SAM and TNM-enriched spatial barcodes (**Fig. 6g; Supplementary Table 9, Methods**) confirmed the upregulation of fibroblast-specific gene markers (*Postn*, *Acta2*, *Col1a1*, *Col1a2*) and specific enrichment of the SAM gene signature (**Fig. 5g**; *Ly6a*, *C1qa*, *Irf7*, *Lgals3*, *Bgn*, *Isg15*, *H2-Ab1*, *H2-Aa*). We further confirmed that the fibroblast-enriched spatial barcodes were significantly enriched with active IFNα-related signaling (**Fig. 6h,i**; GSEA, *P*-value = 0.002), and showed upregulation of expression of NOTCH4 (ADT) and Notch-related genes (**Fig. 6j**). These results validate our oTME findings via direct spatial genomics and confirm that the SAM phenotype is driven by active type-I IFNs and NOTCH4 signaling.

### Tumor infiltration of SAMs is associated with adverse outcomes in cancer patients

Our data from breast cancer models and human specimens have identified spatially-driven macrophage phenotypes with opposing immune activation states. To evaluate the potential prognostic impact of SAM vs. TNM infiltration on breast cancer outcomes, we first identified human macrophage populations with corresponding gene signatures. We projected the macrophage scRNA-seq signatures from oTME and *in vivo* mouse tumors onto a human breast cancer scRNA-seq dataset ^16^(**Extended Data Fig. 13a,b**). We observed good concordance between the *ex vivo* and *in vivo* gene signatures that recapitulated the transcriptional signatures and activation markers of human macrophage subsets reflecting the murine *Ly6a*+SAM (*MRC1*, *C1QA/B/C*, *SPP1*, *GPNMB*, *LGALS3*, *MARCO*, *APOE*, *FN1*, *MMP14*) and TNM (*IL1B, PLAC8, F13A1, ALOX5A, IER2*) subsets (**Supplementary Table 3**). Through these analyses, we defined the matching (Pearson’s correlation = 0.71, *P*-value < 2.2*⨉*10^16^) SAM and TNM signatures (n=38 and n=71 genes, respectively) in human breast cancer macrophages (**Extended Data Fig. 13b, Methods; Supplementary Table 10**).

To ensure the fidelity of these human SAM vs. TNM gene signatures, we cross-validated their specificity leveraging scRNA-seq data obtained from two fully humanized oTME models composed of primary mammary gland fibroblasts, immune cells (PBMC), and TNBC cells originating from tumor epithelial (BR18) or mesenchymal (BR22) primary samples (**Extended Data Fig. 13c, Methods**). To capture a broad spectrum of macrophage activation states (as in **Extended Data Fig. 3c**), we compared the single-cell transcriptomes of tumor-naive stroma/PBMC with oTME cultures (stroma/PBMC/BR18 or BR22) (**Fig. 7a**) following unbiased cell clustering (**Fig. 7b; middle U-MAP**). Consistent with our murine models, the human SAMs were specifically enriched for type-I IFN signals (**GSEA; Extended Data Fig. 13d**) and marked by enhanced expression of immunosuppressive/wound-healing genes including *C1QA/B/C*, *SPP1*, *CD274, LILRB3, TMEM176B, JAML, LGMN, GPNMB*, *APOE, ITGB5, CCL8, ACP5* (**Fig. 7b,c, Supplementary Table 10**) ^96–99^. To confirm the physical location of the C1QA-expressing macrophages in breast cancer tissues, we immunostained (IF) for C1QA and IBA-1 in a cohort of breast cancer tissues. We found that the spatial expression of C1QA was restricted to macrophages that reside in stromal regions of malignant tissues (**Fig. 7d**), corroborating the murine *Ly6a*+SAM signatures (**Fig. 5g**).

**Figure 7:**
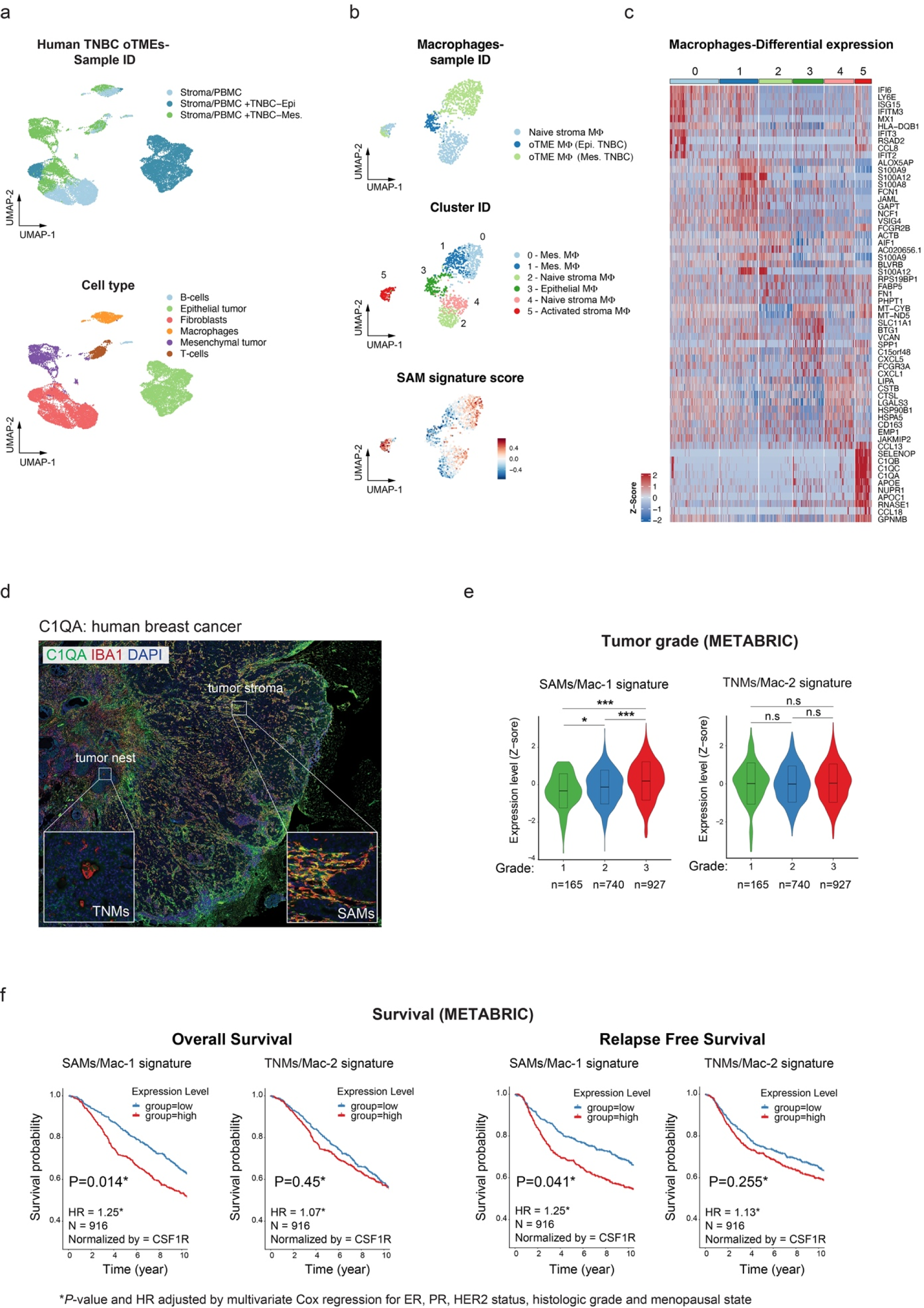
Infiltration of stroma-associated macrophages (SAMs) predicts poor outcomes in breast cancer patients and is associated with aggressive molecular subtypes of breast cancer. (**a**) Two-dimensional UMAP plots of annotated single-cell transcriptome of human oTME cells. Clusters are colored by sample ID (top) and cell type (bottom). (**b**) Two-dimensional UMAP plots of annotated single-cell transcriptome clusters colored by sample ID (top), cluster ID (middle), and SAM gene signature score (bottom). (**c**) Heatmap showing differentially expressed genes in human oTME macrophages. Average expression level (Z-score; color scale) of signature genes (rows) across each subset (columns) are shown. (**d**) IF analysis for IBA1/C1QA in human breast tumors, mapping the spatial localization of C1QA+ macrophages to stromal regions of the TME. Scale bar, 100μm. (**e**) Expression levels of SAM/Mac-1 and TNM/Mac-2 gene signatures in different neoplasm histologic grades of breast cancer (METABRIC). Wilcoxon Rank Sum test, \**P* < 0.05, \*\**P* < 0.01, \*\*\**P* < 0.001, otherwise not significant (n.s). (**f**) Kaplan-Meier plots showing overall survival (OS) and relapse-free survival (RFS) in breast cancer patients (n=916) with higher and lower expression of CSF1R-normalized SAM (31 genes) and TNM (71 genes) gene signatures in the METABRIC cohort. *P*-value and hazard ratio (HR) were calculated by multivariate Cox regression adjusting for ER, PR, HER2 status, tumor histologic grade, and menopause status (**Methods**). The 10-year survival is shown (x-axis), but the analyses were conducted using all of the data.

To determine the prognostic impact of SAM and TNM gene sets in patients, we applied these signatures to the Molecular Taxonomy of Breast Cancer International Consortium (METABRIC) cohort ^100^. A higher correlation with the SAM signature was detected in more aggressive and mesenchymal molecular subtypes, such as Claudin-low and Basal, compared to the normal-like and luminal A/B subtypes (**Extended Data Fig. 13e**). In addition, the SAM signature was significantly associated with higher tumor histologic grade while the TNM signature was not (**Fig. 7e**). To ensure that our survival analysis specifically evaluated the impact of SAM vs. TNM phenotypes (rather than changes in macrophage tumor infiltration), we normalized the gene signatures by *CSF1R* expression to account for the variation in total macrophage infiltration across different tumors ^14^. In agreement with the proposed proliferative and immunosuppressive phenotype of SAMs, we observed that higher expression of SAM genes (upper vs. lower quartile) was significantly associated with shorter overall survival (OS; hazard ratio (HR) = 1.24, 95% CI 1.05-1.42, N=916, *P*-value = 0.02), disease-specific survival (DSS; HR = 1.29, 95% CI 1.04-1.54, N=914, *P*-value = 0.04), and relapse-free survival (RFS; HR = 1.25, 95% CI 1.03-1.47, N=916, *P*- value = 0.04) in breast cancer patients (**Fig. 7f and Extended Data Fig. 13f**), even after accounting for ER/PR and HER2 status, menopausal state, and tumor grade in a multivariate Cox analysis (**Supplementary Table 10**). Conversely, higher expression of TNM genes was not associated with OS (HR = 1.07, 95% CI 0.90-1.25, *P*-value = 0.44), DSS (HR = 1.12, 95% CI 0.88-1.35, *P*-value = 0.35) or disease recurrence (HR = 1.13, 95% CI 0.93-1.35, *P*-value = 0.23). These data show that the spatially-defined SAM phenotype, rather than overall macrophage abundance or ontogeny, is associated with adverse prognostic impact in breast cancer, across molecular subtypes.

## Discussion

Tumor-infiltrating macrophages play a critical role in promoting tumor growth, immune evasion, and therapeutic resistance ^5,101^ highlighting macrophages as a promising target for therapeutic intervention ^10,102–108^. In solid tumors, including breast cancer, tumor-infiltrating macrophages undergo two fundamental phenotypic alterations; (i) they adopt wound healing-associated activities in response to complex microenvironmental signals ^16,99^, and (ii) regain their proliferative capacity in response to mitogenic signals to accumulate *in situ* ^12,45^. While the transcriptional and proteomic details of these alterations were extensively annotated with relevance to patient survival ^16,70,109^, the specific TME interactions that shape these phenotypic alterations remain incompletely understood.

To address this challenge, we developed scalable organotypic models of murine and human breast TME (oTME) that recapitulate with high phenotypic fidelity the functions, heterogeneity, spatial organization, and cell-cell interactions of macrophages with tumor epithelial cells and stromal fibroblasts. We harnessed these models to define the dynamics, spatial, and molecular mechanisms underpinning macrophage heterogeneity and self-renewal. We found that regardless of their ontogeny (monocyte-derived vs. resident), the common denominator of macrophage functional diversification in breast cancer was their spatial localization in tumors.

By exploring the transcriptional dynamics during macrophage education, we revealed a subpopulation of Sca-1^+^F4/80^high^CD11b^high^ self-renewing macrophages that was instigated by type-I IFN signaling. The faithful recapitulation of macrophage proliferation in mammary tumors allowed us to mechanistically address the central question of what drives the long-term accumulation of macrophages during tumor growth. Our finding that TME macrophages can self-renew helps explain why macrophage abundance in mammary tumors was unaffected by ablation of monocyte recruitment (*Ccr2*-KO MMTV-PyMT) ^45^. This finding is of particular interest, as it is well accepted that upon differentiation in healthy tissues, macrophages become predominantly quiescent despite active CSF-1R/CSF-1 signaling ^44,52^. However, in pathological conditions, including cancer, macrophages regain their proliferative capacities in response to CSF-1R signaling ^32,45,110–112^. The dependency of CSF-1R has led to the development of multiple CSF1R targeting agents that are under clinical investigation in various cancer types ^113–115^. However, the abundant expression of CSF-1 both in healthy and malignant mammary tissues ^50^ also raises the question of why only TME macrophages can proliferate in response to CSF-1, while macrophages in healthy tissues remain refractory to the same mitogenic signals.

To address this context-dependent response, we explored the molecular pathways that co-existed in the Sca-1^+^F4/80^high^ proliferating subset (**Fig. 2**). We found that active type-I IFN (via STING) was essential to shift the response toward CSF-1 from survival to proliferation. In addition, by exploring the spatial distribution and immunophenotyping of breast cancer macrophages, we identify a spatial determinant of macrophage proliferation in mammary tumors (**Fig. 3, Extended Data Fig. 5-6**). Specifically, we found that the Sca-1+F4/80^high^CD11b^high^ subset in murine or the Ki67^+^CD163^high^ macrophages in human, share a common spatial distribution, and are found in close proximity to cancer-associated fibroblasts (CAFs). Conversely, macrophages that reside in tumor epithelial nests (tumor nest macrophages; TNMs) were Sca-1^neg^F4/80^int^CD11b^low^ and largely Ki67 negative despite comparable CSF-1 expression. Importantly, we found that fibroblast-induced proliferation was consistent irrespective of macrophage origin, suggesting that local spatial signals override ontogeny in dictating macrophage behavior. Our finding that macrophage proliferation was facilitated only by cell contact with tumor-activated fibroblasts and not tumor-naive fibroblasts, revealed a three-way interaction where tumor cells activate fibroblasts, which in turn induce macrophage proliferation.

The contact dependency of macrophage proliferation revealed Notch signaling, and specifically Notch4 as a key driver. Previous reports in breast cancer have shown that CAFs upregulate several Notch-activating ligands ^24,116^ and enable Notch activation in neighboring cells, including macrophages. Although Notch signaling in breast cancer macrophages was shown to mediate their immunosuppressive phenotype in tumors ^24,45,117^, the critical link between Notch4 and macrophage self-renewal was previously unknown. Using syngeneic breast cancer models (MMTV-PyMT) and TNBC PDX models, we demonstrated that treatment with anti-Notch4 antibodies resulted in significant inhibition of tumor growth and metastasis, coupled with a selective reduction in Ki67^+^ macrophages. Collectively, our results revealed macrophage self-renewal as a novel targetable cancer vulnerability.

Importantly, the spatial localization of TME macrophages impacted not only their self-renewal capacity, but also their broader phenotype. Our results across mouse models and human specimens provided compelling evidence that immunomodulatory phenotypes in breast cancer macrophages are regulated by their local spatial interactions. Here too, independent of macrophage ontogeny, SAM macrophages interacting with CAFs displayed granular morphology, were highly phagocytic (TIM-4^high^), and expressed immune-suppressive markers such as CD206 and PD-L1 ^91,118,119^. In contrast, TNMs exhibited dendritic-like morphology, lower scavenging potential, and an immune-stimulatory (CD86^high^ CD11c^high^) phenotype (**Fig. 5, 6**), with higher lymphocyte infiltration. Importantly, our *ex vivo* replating (**Extended Data Fig. 8c**) and *in vivo* repopulating (**Extended Data Fig. 9**) experiments demonstrated that: (**i**) TME interactions effectively override pre-existing signatures of macrophages (SAM or TNM), (**ii**) monocytes can adopt the same functions/phenotypes as tissue-resident macrophages (unlike in healthy organs ^25,41^), and (**iii**) SAMs are more resistant to CSF-1R depletion treatment.

Furthermore, the enhanced sensitivity of TNMs towards CSF-1R mAb targeting (**Extended Data Fig. 9**) has unexpectedly highlighted a major safety concern as TNMs represent the pro-inflammatory subset that supports lymphocyte infiltration which predicts favorable outcomes in breast cancer patients ^11^. Moreover, the cessation of CSF-1R treatment was associated with accelerated tumor growth, infiltration of immunosuppressive SAM, and a substantial influx of granulocytes into treated tumors. Of note, the protective role of macrophages in injured tissues was previously reported in a wound-healing model ^29^, specifically by limiting granulocyte “swarming” and excessive tissue damage. As macrophage depletion (via CSF1R targeting) is being clinically evaluated across human cancers ^13,115^, this unexpected adverse response requires further mechanistic exploration.

Collectively, these data further support the notion that spatial signals, rather than macrophage ontogeny ^25,41^, function as a key driver of macrophage phenotypic heterogeneity and self-renewal in mammary tumors. We validated the importance of these findings to human breast cancer biology by demonstrating that the phenotypic compartmentalization of TME macrophages was highly conserved in human breast tumors and conferred prognostic relevance. Altogether, these results define SAMs as a primary and replenishing source of immunosuppressive signals and highlight the clinical potential for therapeutic interventions geared at limiting the proliferation of tumor-infiltrating macrophages.

## Author contribution

N.B.-C. and D.A.L. conceived the project. N.B.-C., J.S., M.S.J., P.L.L., A.D.S., V.K.R., C.M.S., and S.G. performed experiments and with the help of N.B.-C., D.A.L., X.N., J.A.J., and F.G., interpreted results. X.N. and S.K., performed the bioinformatics analyses. N.M.I., D.D.D., and K.A.B. provided human breast cancer specimens and interpreted clinical data. I.S. and A.H. provided the Adam17 inhibitors. S.W.L and J.H.H., provided mouse and human models. N.B.-C., X.N., and D.A.L. wrote the manuscript with inputs from all authors. D.A.L supervised the study. All authors reviewed and edited the manuscript.

### Acknowledgments

We thank the members of the Landau laboratory and Robert L. Bowman for their advice and insightful discussions. This work was supported by the Daedalus Award (fund 55000263), Weill Cornell Medicine. D.A.L. is supported by the Burroughs Wellcome Fund Career Award for Medical Scientists, the Vallee Scholar’s Award, the Leukemia Lymphoma Scholar Award, the Mark Foundation Emerging Leader Award, and the NIH Director’s New Innovator Award 11 (DP2-CA239065).

## RESOURCE AVAILABILITY

### Code Availability

The pipelines for scRNAseq analyses are provided as Supplementary Notes (Methods). The oTME cells used in this study will be provided under a material transfer agreement.

### Data Availability

The bulk, single-cell RNA sequencing and SPOTS data reported in this paper can be accessed under the GEO SuperSeries GSE171463 with access token *uzoxuiqejvebrub*. R scripts for the single cell RNA-seq analyses are included in the **Supplementary Notes**. Other materials used in this work will be provided upon request by Dan A. Landau or Nir Ben-Chetrit

## EXPERIMENTAL MODEL AND SUBJECT DETAILS

### RNA isolation for qRT–PCR

RNA from cells was extracted using TRIzol™ Reagent (Thermo Fisher Scientific; 15596026) and 1-5ug RNA were used to prepare cDNA using RNA to cDNA EcoDry™ kit (Clontech; 639549). RNA from FACS-sorted cells was extracted using TRIzol™ LS Reagent (Thermo Fisher Scientific; 10296010) and processed according to manufacture’s protocol. For qRT-PCR reactions, mouse Taqman probes (Applied Biosystems) were used for quantifying expression of *Cdh1* (Mm01247357_m1), *Csf1* (Mm00432686_m1), *Csf2* (Mm01290062_m1), *Csf3* (Mm00438334_m1), and *Vim* (Mm01333430_m1). Expression values were normalized to both *Hprt* (Mm03024075_m1) and *Gapdh* (Mm99999915_g1) as housekeeping genes.

### Human breast tissue samples

Informed consent was provided by patients who underwent mastectomy for breast cancer treatment at Memorial Sloan Kettering Cancer Center (MSKCC). Institutional Review Board approval was obtained from MSKCC and Weill Cornell Medicine (New York, NY), under IRB protocol 10-040. Archived formalin fixed paraffin embedded (FFPE) breast tissue specimens were obtained under a standard tissue acquisition protocol.

### Human oTME of TNBC

data were processed with CellRanger as previously described before being analyzed with Seurat. Cells with over 15% mitochondrial gene expression by counts were removed, and doublets were detected per sample and removed using DoubletFinder^120^. Cells were then normalized with scTransform v2 using 3000 highly variable features^121^. The top 20 principal components were then computed and used for clustering using the FindClusters function, with a resolution of 0.2. Cells were annotated per-cluster according to marker gene expression, as shown in Supplementary Figure X. Differentially expressed genes were computed with the Wilcoxon rank sum test as previously described, and the data was visualized using UMAP. Next, in order to characterize the macrophage compartment, the macrophages were subsetted and re-normalized using the log-TPM method. Genes were z-scored using ScaleData, 2000 highly variable features were selected, and the top 20 principal components were selected for clustering and visualization with UMAP. Differential expression was performed in the same way, and the stroma-associated macrophage (SAM) and tumor nest macrophage (TNM) human signatures were derived through a pairwise differential expression expression test between the C1Q, APOE-high expressing cluster and the cluster of macrophages enriched for cells from the tumor epithelial sample. The expression of the top 10 differentially expressed genes per cluster (one vs. rest) were visualized as a heatmap using the ComplexHeatmap package in R^122^.

### Isolation of stromal fraction from human breast tissue

Breast tissue was obtained from women undergoing reduction mammoplasty surgery at Weill Cornell Medicine in accordance with approved Institutional Review Board protocols 20-01021391 and 1510016712. All patients from whom tissues were collected provided written informed consent. Approximately 25 ml of tissue from each patient was finely minced and digested for 1 hour at 37°C in Ham’s F-12 media (Corning) supplemented with 10% fetal bovine serum (FBS) and 1% penicillin/streptomycin with 10 mg/ml collagenase type 1 and 10 µg/mL hyaluronidase (Sigma Aldrich) in a final volume of 50 ml. After digestion, samples were pelleted to separate the stromal vascular fraction (SVF) by centrifugation at 500g. The pellet was washed, resuspended in Ham’s F-12 media, and passed through a 100 µM filter followed by a 40 µM filter. The final filtrate was centrifuged at 500g and the pellet was treated with 1 ml of Red Cell Lysis Buffer (Sigma Aldrich) following the manufacturer’s protocol. After treatment, the samples were re-pelleted, washed, and resuspended in Ham’s F-12 media for experimentation.

### Mouse work and strains

Animal procedures were approved by the Institutional Animal Care and Use Committee (IACUC) of the Research Animal Resource Center (RARC) at Weill Cornell Medicine (Protocol: 2016- 0058). MMTV-PyMT (Stock No: 022974), Arg1-EYFP (Stock No: 015857), and wild-type C57BL/6J (Stock No: 000664) were purchased from The Jackson Laboratory (USA). *Rosa26^mTmG^* (Stock No: 007676, mTmG) mice were a gift from Dr. Johanna Joyce (MSKCC).

### Tumor engraftment and processing

Females of the MMTV-PyMT model developed spontaneous tumors after 100-120 days post birth. To generate orthotopic tumor transplants, tumor cells (1.0 × 10^6^) were injected in 50µL of 50% Matrigel (Corning) in serum-free DMEM, into the 4th mammary gland of 7-10 week-old wild-type (WT) C57BL/6 female mice. Prior to injection, mice were anesthetized with 2% isoflurane and cells were injected under the nipple into the fat pad. Tumor dimensions (width and length) were measured using a digital caliper and tumor volumes were calculated as V = (L × W^2^)/2.

### Neutralization of Notch4 *in vivo*

To target Notch4 in mammary tumors, wild-type C57BL/6 female mice were engrafted with tumor epithelial cells in the 4th mammary glands and allowed to establish palpable tumors. Then, mice were pooled and randomized into two arms: vehicle-treated (PBS) or Notch4-treated with anti-Notch4 monoclonal antibodies (BioXcell, clone HMN4-14). Mice were dosed every 3 days intraperitoneally with 15μg/kg body weight.

### Generation of mouse bone marrow-derived macrophages (BMDMs)

Mouse BMDMs were obtained from femurs and tibias of 6-8-week-old B6 mice. Bone marrow cells were flushed onto a 40µM strainer using a 25G x 5/8 needle washed with RPMI. Bone marrow cells were gently meshed through the 40µm strainer using a 1ml plunger. After centrifugation at 300g 4°C, for 5 minutes, cells were resuspended in DMEM medium, 10% FBS and 1% Pen-strep supplemented with 10ng/mL recombinant murine CSF-1. The growth medium was replenished every other day with fresh 10ng/mL CSF-1. On day-7, >95% of the cells were CD11b+ Ly6C/Ly6G- (Gr1-) F4/80+ macrophages.

### Mouse bone marrow monocyte isolation

Mouse bone marrow was obtained from femurs and tibias as described above for differentiation of BMDMs. Bone marrow cells were resuspended with 5 mL of RBC Lysis Buffer (ThermoFisher Scientific) for five minutes incubation on ice to remove red blood cells, washed with serum-free RPMI, and monocytes were purified using negative selection monocyte isolation kit (130-100- 629; Miltenyi).

### Chemicals and Biological reagents

99LN parental (oTME) cells were generated in the Johanna Joyce lab as previously described ^76^. Isolations of tumor epithelial or stromal cells were performed by FACS-sort using EpCAM and PDGFRA antibodies. Cells were maintained in DMEM-GlutaMAX (Gibco; 10566016) supplemented with 10% FBS (Gibco), and 1% Pen-strep (ThermoFisher Scientific). Cells were routinely verified to be mycoplasma-free using a commercial kit (Lonza). CSF-1, IL-4, and IL-13 recombinant proteins were from Peprotech.

### EdU incorporation

Cells were incubated with 10µM EdU (5-ethynyl-2’-deoxyuridine) without changing the conditioned growth medium for the required duration and analyzed by flow cytometry according to the manufacturer’s instructions (A10202; Thermo Fisher Scientific). For EdU imaging, cells were plated on coverslips, incubated with EdU (10µM), fixed with 4% paraformaldehyde (PFA), and immunostained with desired antibodies prior to the EdU staining protocol (C10337).

### BrdU incorporation *in vivo*

Tumor-bearing mice were injected with BrdU (2mg/mouse/intraperitoneally) 24hrs before tumor harvesting. Single cell suspensions were stained with anti-BrdU APC antibodies following the manufacturer’s protocol (BD biosciences; APC BrdU Flow Kit 552598) and analyzed by flow cytometry.

### Immune cell isolation for flow cytometry from mammary gland or mammary tumors

MMTV-PyMT tumors were collected from euthanized female mice, washed in cold PBS and digested by mechanical dissociation, using gentleMACS™ Dissociator (Miltenyi Biotec) and the mouse tumor dissociation kit (Miltenyi Biotec) according the manufacturer’s instructions. To remove cell clumps and undigested tissues, cell suspensions were passed through 70μm and then 40μm filters, and equal cell numbers were analyzed by flow cytometry. To isolate cells from murine mammary glands, the 4th and 5th glands were collected and digested with 4mg/ml collagenase type 4 (porcine origin, Sigma), 4 μg/ml DNase I (Sigma) at 37°C with periodic vortexing. Cells were further mashed through 100μm filters, and then passed through 40μm filters. Cells were collected, counted, and analyzed by flow cytometry.

### Chromium 10x single-cell RNA-seq

*Ex vivo*: Single cell solutions from oTME cultures were harvested using trypsin, clumps were removed through a 40μm filter, and single cells (n=4,000 per condition) were loaded on the Chromium platform to generate cDNA following the manufacturer’s protocol. Single-cell cDNA libraries were prepared using the Chromium Single Cell 3′ Library & Gel Bead Kits v2 (PN- 120237, PN-120236, PN-120262) according to the manufacturer’s instructions. Samples were sequenced at an average of 50,000 reads per cell.

*In vivo*: undigested tumor fragments from spontaneous mammary tumor (MMTV-PyMT) were collected, washed twice with cold PBS, and digested. Single cells were collected and processed as above.

### Whole mount imaging

For whole-mount immunofluorescence imaging, approximately 3mm^3^ pieces of inguinal fat-pad from 12-week-old PyMT-MMTV females were incubated in 4% PFA (Electron Microscopy) diluted in PBS for 30 min at room temperature with agitation and permeabilized with PBS Triton 0.3% (Sigma T8787) 4% BSA for 1 hour at room temperature. Samples were then stained with directly conjugated antibodies in PBS 0.03% Triton for 1 hour. Samples were rinsed with PBS 3X and mounted on cavity slides (Sigma) with Fluoromount G (eBioscience). The antibodies used are the following: Anti-mouse F4/80 eFluor570 (eBioscience, clone BM8 (1/200), anti-mouse EpCAM Alexa Fluor 488 (Biolegend clone G8.8, 1/100), Anti-mouse CD140a Alexa Fluor 647 (eBioscience: APA5; PDGFR-a, 1/100). Z-stacks of 30μm to 60μm with 0.8μm consecutive intervals and tile scans were acquired using LSM880 Zeiss microscope with 40x/1.3 objective (oil).

### Flow cytometry and fluorescence-activated cell sorting (FACS)

Flow cytometry data were collected using BD LSRFortessa, BD FACSCanto II, and BD FACSAria III was used for FACS-sort. FlowJo X was used for data analysis and generation of flow plots for figures. For analyzing live cells from tissues by flow cytometry, mice were anesthetized with a ketamine/xylazine cocktail and perfused with 30mL cold PBS using cardiac puncture. Cells from dissociated tissues were filtered through 70μm then 40μm filters to generate a single-cell suspension. Cells (1-2 X10^6^) were then incubated with 2X Fc Block solution (1:50, CD16/32 BD Biosciences) in FACS-buffer (2% FBS-PBS, 2mM EDTA) for 20 min at 8°C. Conjugated antibodies were added to cells and incubated for 30 min at 8°C in the presence of the Fc Block solution. Stained cells were washed twice and resuspended with a FACS-buffer containing DAPI (1μg/mL) to exclude dead/compromised cells. Mammary gland Eosinophils were defined as CD45+CD11b+F4/80^low^SiglecF+ and excluded together with DAPI+ dead cells by staining with BV421-SiglecF.

### CFSE T-cell proliferation assay

T-cell proliferation was measured using CFSE assay (ThermoFisher Scientific). CD3+ T-cells were negatively isolated (EasySep Mouse T Cell Isolation Kit) from the spleen of a WT mouse, labeled with 5µM CSFE and stimulated with CD3e (1:100) and CD28 (1:500) activating antibodies (ThermoFisher Scientific) in serum-free RPMI for 20 minutes, 37°C. Activated CFSE-labeled T cells were seeded either alone or with: (i) CSF-1-treated BMDMs, (ii) oTME CM-educate BMDMs, or (iii) co-cultured with oTME/BMDMs in DMEM growth media contained 10% FBS (Gibco), 1% P/S. Five days later, CD8+ and CD4+ cell divisions were analyzed by flow cytometry by quantifying the CFSE dye dilutions.

### Immunohistochemistry

Mice were anesthetized with xylazine/ketamine and transcardially perfused with 30mL cold PBS, followed by 10mL cold 4% PFA. Tissues were collected and fixed in 4% PFA overnight, washed, transferred to 70% Ethanol, and embedded in paraffin blocks. Paraffin sections (10-12μm) were mounted on plus slides, de-waxed in xylene and hydrated into graded alcohol solutions. Endogenous peroxidase activity was quenched by immersing the slides in 1% hydrogen peroxide in PBS for 15 minutes, room temp. Antigen retrieval pretreatment was performed in a steamer using the appropriate buffer for 30 minutes. Sections were incubated overnight with primary antibody at 4°C. For DAB staining, sections were washed with PBS and incubated with the appropriate secondary antibody followed by avidin-biotin complexes (Vector Laboratories, Burlingame, CA, Cat. No. PK-6100). Antibody reaction was visualized with 3-3’ Diaminobenzidine (Sigma, Cat. No. D8001) followed by counterstaining with hematoxylin. Tissue sections were dehydrated in graded alcohols, cleared in xylene, and sealed with coverslips. For immunofluorescence staining, slides were stained fluorescently-labeled secondary antibodies (Invitrogen) for 1 hr at room temperature, counterstained with DAPI (5μg/mL) for 5 min, washed and sealed with VECTASHIELD® Antifade Mounting media.

### Immunofluorescence and image processing

Cells were grown on coverslips for 48 hours. After treatments, cells were washed, permeabilized using 0.02% Triton X-100 and fixed for 20 min with 4% PFA. Cells were blocked with 5% BSA and incubated with primary antibodies overnight at 4°C. Probing with AlexaFluor-488, AlexaFluor-555, or AlexaFluor-647 fluorescent secondary antibodies (Invitrogen) was carried out for 1 hr at room temperature. Slides were sealed (VECTASHIELD® Antifade; H-1000) and images were taken using an inverted fluorescence Nikon microscope, or LSM880 Zeiss confocal microscope. Images were processed using Photoshop (version 21.1.2) and analyzed by Fiji/ImageJ (version 2.1.0) software.

### Immunoblotting analysis

Cell lysates were collected, cleared, and processed as previously described ^123^. Samples were loaded on Mini-PROTEAN®Precast 4–15% gradient gels (BioRad; 456-1083). Antibodies against ARG1 (#79404), Notch Activated Targets Antibody Sampler Kit (#68309), phospho-SMAD2/3 (#8685), CSF-1R (#3152), phospho-AKT Ser473 (#4060), AKT (#9272) and phospho-ERK (#9101) were purchased from Cell Signaling.

### Cytokine arrays

Mouse XL Cytokine Arrays were purchased from R&D Systems and used according to the manufacturer’s instructions. Briefly, similar volumes of conditioned media were collected, debris was removed by centrifugation (2,000g, 10 min 4°C), and cleared supernatants were loaded on spotted membranes for 16hrs, 4°C with tilting. Secreted cytokines, chemokines, and growth factors (111 in total) were probed in duplicate along with positive and negative controls.

### Spatial protein and transcriptome sequencing (SPOTS)

SPOTS protocol was performed as previously reported ^124^. In advance of SPOTS, sections were obtained for RNA quality measure with the RNeasy Plus kit. Tissues with RNA integrity number (RIN) of 7 or greater were used for future experiments. The 10x Genomics Visium Spatial Gene Expression kit and the associated Tissue Optimization kit were implemented for this study. 10µm thick, OCT-embedded sections were deposited on the Visium slides; slides were stored at -80°C for up to 1 week in advance of the experiment.

The following solutions were made immediately in advance of the experiment and were kept on ice. Their compositions are as follows: 2x blocking buffer (400µl needed per sample): 6X SSC, 0.2% Tween 20, 4% BSA, 0.2µg/µl sheared salmon sperm DNA, 2 U/µl RNAse inhibitor; saponin mixture (75µl per sample): 5x saponin, 0.4U/µl RNAse inhibitor, remaining volume is compensated with PBS; antibody staining mix (50µl per sample): 1x blocking buffer (dilution of 2x blocking buffer above), 20µM blocking oligos (dT25), fluorescent and TotalSeqA antibodies as needed; 1x blocking buffer (75µl per sample): 1x blocking buffer, 5mM ribonucleoside vanadyl complex (heated to 65°C), Fc block (5µl per sample); wash buffer (600µl per sample): 1x blocking buffer. Any remaining volume was compensated with water unless noted otherwise.

Slides were moved on dry ice to the thermocycler and then incubated for 1 minute at 37°C on the Thermocycler Adaptor. The slides were then immersed in 1% PFA in PBS for 10 minutes at room temperature to fix the tissue. The slides were washed by dunking five times into a falcon tube with 3X SSC and then secured in the cassette. Each sample was blocked in 75µl of 1x blocking buffer for 15 minutes at 4°C. Post-blocking, 75µl of the saponin mixture was added to each sample at 4°C for 10 minutes for initial permeabilization; this step was optimized for breast tissue using the Tissue Optimization kit. The saponin was removed and 50µl of antibody staining mix was added to each sample. The slide was then incubated, protected from light, for 90 minutes at 4°C with the antibodies. The samples were then washed with 100µl of wash buffer four times for one minute each. The entire slide was then washed by dunking the slide twenty times in a falcon tube with 3X SSC and then imaged. After imaging, the tissue was permeabilized a second time for 27 minutes using the Tissue Optimization Tissue Removal Enzyme (per sample: 3.5µL Tissue Removal Enzyme (3000387), 59.5µl 1X SSC, 7.0µl 10% SDS); this second permeabilization time was also optimized using the Tissue Optimization kit. After permeabilizing, the slides were washed three times with 150µl of 0.1X SSC. We followed the 10x Genomics Gene Expression protocol, starting with step 2.2 Reverse Transcription, for the remainder of the experiment. The only deviations from the original protocol are the addition of additive primer to amplify the ADT during Second Strand Synthesis and cDNA Amplification (8.8µl of 100uM and 4µl of 0.2µM additive primer were added to each reaction, respectively) and retention of the supernatant in step 4.3 cDNA Cleanup - SPRIselect. Two sequential 1.9X SPRIselect cleanups (130µl and 95µl of SPRI for the first and second cleanups, respectively) were performed on said supernatant and the final product was eluted in 45µl of water.

The ADT product was then amplified and indexed using the SI-PCR and TruSeq Small RNA RPIx primers before Illumina sequencing. The master mix and temperatures for this reaction were as follows: 45µl ADT, 50µl of Amp Mix (2000047), 2.5µl of 20µM TruSeq Small RNA RPIx primer, 2.5µl of 20µM SI-PCR primer; 95°C for 3 minutes > (95°C for 20 seconds, 60°C for 30 seconds, 72°C for 20 seconds)10x > 75°C for 5 minutes. Approximately 50,000 reads were designated per spatial barcode for gene expression and 8,500 reads per spatial barcode were designated for ADT sequencing.

### Quantification and statistical analysis

Parameters such as sample size, precision (mean±SD or SEM), and statistical test significance are reported in the text, Figures, and Figure Legends. Statistical significance was calculated by one-way ANOVA when comparing two groups or two-way ANOVA when comparing three or more groups. A *p*<0.05 was considered statistically significant. *Prism v8* software was used for statistical data analysis.

### Bulk RNA-Seq Analysis

BMDMs were co-cultured with oTME cells (educated) or left alone with CSF-1 for 10 days (n=3 replicates, each). Then, BMDMs were FACS-sorted and RNA was extracted using Trizol-LS. RNA-sequencing libraries were generated with the SMART-Seq preparation kit (CloneTech) with the Nextera XT kit (Illumina), and pair-end 150-bp sequencing was performed by GeneWiz (South Plainfield, New Jersey, USA) on an Illumina HiSeq 2500. FASTQ files were mapped to the mouse genome (mm10) using *STAR (version 2.5.3a)* with default parameters ^125^. Transcript count was quantified using STAR *-quantMode* option with Gencode mouse release M21 annotation GTF files (https://www.gencodegenes.org/mouse/release_M21.html). The resulting count matrix was analyzed and normalized using *DESeq2 v1.18.1* ^126^. Differential gene expression was assessed with default parameters. We defined differentially expressed genes as any gene with an absolute log-fold change (logFC) larger than 1 at a false discovery rate (FDR) of 0.01. The DESeq2 output results (**Figure 1e**) are listed in **Supplementary Table 1**. To generate the volcano plot, we removed the genes that: (i) have NA logFC values, and (ii) genes below average count of 3 in CSF- 1 treated. For the genes that have *P*-values of zero, the -log10 *P*-values were set to 0.1 times the smallest non-zero *P*-value. The Gene Set Enrichment Analysis (GSEA) of Gene Ontology (GO) was performed using the *gseGO* function in the *clusterProfiler v3.16.0* R package (^127^), with 10,000 permutations and other default parameters.

### Single Cell RNA-seq Analysis

The scRNA-seq data were processed using Cell Ranger (v1.3.1) with default parameters (https://support.10xgenomics.com/single-cell-gene-expression/software/overview/welcome). Samples were sequenced at an average of 50,000 reads per cell. Raw sequencing data were demultiplexed and post-processed following the Cell Ranger pipelines provided by 10x Genomics. Briefly, raw base calls were demultiplexed into fastq files using the cellranger *mkfastq* command, followed by alignment to the Cell Ranger mouse (mm10) genome reference 3.0.0. Barcode and UMI counting were performed using the cellranger *count* command with default parameters. The *in vivo* single-cell RNA-sequencing data were generated in the same way except they were processed using Cell Ranger (v3.0.1) and aligned to the Cell Ranger human (hg19) and mouse (mm10) genome reference.

R markdown output with codes for the scRNA-seq analyses are provided as Supplementary Notes (named as oTME Macrophage scRNA-seq Analysis and MMTV-PyMT Macrophage scRNA- seq Analysis respectively). The oTME and the MMTV-PyMT scRNA-seq samples were analyzed as described in the following sections. We excluded the oTME cells that were cultured alone (2 and 10-days) only for visualization purposes.

### Preprocessing

The count matrices of all six oTME (*ex vivo*) samples were pooled and normalized using log-transformed transcript per million (logTPM). Specifically, we denoted the UMI count of *j_th_* gene in *i_th_* cell as *Count_i_*_,*j*_, and the logTPM was calculated as*logTPM_i_*_,*j*_ = 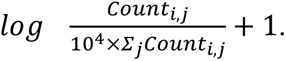 We used the *Seurat V2* R package ^128^ for downstream analysis. Low-quality cells with fewer than 500 genes detected or mitochondrial gene percentage >10% were filtered out. Genes that were expressed in fewer than 50 cells were also removed. After filtering, we retained 19,280 cells in total with an average of 13,013 ± 56.49 UMIs per cell (mean ± s.e.m.) and an average of 2,947 ± 7.49 (mean ± s.e.m.) genes detected across all cells. A set of 1,000 highly variable genes were identified using the *FindVariableGenes* function with default parameters, which finds variable genes while controlling for the strong relationship between variability and average expression.

### Dimensionality reduction

We scaled and centered the normalized count data using *ScaleData* function with default parameters to calculate z-score for each gene. We then performed Principal Component Analysis (PCA) using the *RunPCA* function with the 1,000 highly variable genes identified from the preprocessing step. We selected the first 15 PCs for downstream clustering and dimensionality reduction analysis based on the observation of the PC “elbow” using the *PCElbowPlot* function. We used tSNE ^129^ to visualize the scRNA-seq data. We selected the top 15 PCs and used the *RunTSNE* function to perform tSNE dimensionality reduction to embed the data into two dimensions.

### Clustering

We used the modularity based shared nearest neighbor (SNN) clustering algorithm implemented in Seurat’s *FindClusters* function with *resolution=1* and other default parameters using the top 15 PCs. Initially we retrieved 19 clusters. We further validated the clustering result by constructing a phylogenetic tree of the 19 clusters using their average expressions. We used the *BuildClusterTree* function with the top 15 PCs as input to calculate the distance between clusters. We assessed the clustering quality using the *AssessNodes* function to calculate the out of bag error for a random forest classifier trained on the bottom 25% of the nodes. Nodes with an out of bag error bigger than 0.03 were merged together. We repeated the above steps until all the nodes had an out of bag error below 0.03. We noted that one of the clusters expressed heterogeneous lineage markers of epithelial (Epcam), CAF (Fn1, Acta2), and basal-like (Cd24a) cells. It also exhibited higher UMI counts per cell compared with the non-cycling cells. These features are common to doublet/multiplet cells that are considered as artifacts of droplet-based scRNA-seq technologies ^120,130^. Therefore, we annotated this cluster as doublets and removed them from downstream analysis. Macrophage clusters were taken from the data for further analysis. We noted that cells from one of the clusters expressing Mki67 were the cycling macrophages. They are heterogeneous cells that contain both cycling CSF-1-treated and cycling educated macrophages, but this biological difference was masked by the strong cell cycle effect. Therefore, we dissected this cycling population by following the same procedures as described in the *Preprocessing, Dimensionality reduction*, and *Clustering* sections with the top 10 PCs and an out of bag error of 0.07. In addition, we further identified and removed a group of doublet cells of macrophages and CAFs. The macrophages were clustered mainly by education status (CSF-1-treated vs. educated), time points (early vs. late), and cell cycle phases (non-cycling vs. cycling). Thus, we annotated them accordingly as CSF-1-treated CD11c+, CSF-1-treated, CSF-1-treated-cycling, early, early-cycling, late, late-cycling macrophages. In total, we identified 2 clusters of Epcam+ tumor cells, 3 clusters of Cd24a+ Epcam-Basal-like cells, 3 clusters of Acta2+ CD24^neg^(CAFs), and 7 clusters of Cd68+ CSF-1-treated and educated macrophages. The cell type annotation and metadata can be found in **Supplementary Table 7.**

### Differential expression

We first performed differential expression analysis using the Wilcoxon Rank Sum test implemented in Seurat’s *FindAllMarkers* function. We defined two sets of differentially expressed genes. First, we defined 4 cell lineages based on known markers and grouped the clusters into Epcam+ tumor epithelial cells, Acta2+ CAF cells, Cd24+ Epcam-basal-like cells, and *Cd68*+ macrophages lineages. We conducted differential express analysis among these 4 lineages with the parameter *min.pct=0.5* which only tests genes that were detected in a minimum percentage of 50% of the cells in each lineage in order to get a consensus list of differentially expressed markers. Genes with a log fold change greater than 1 and a *P*-value of less than 0.01 were considered lineage markers. Second, we defined a set of cell-type markers by comparing the clusters within each lineage (i.e cycling tumor epithelial cells vs. all tumor epithelial cells). In addition, non-cycling and cycling macrophage clusters were separately analyzed for differential expression. We used the parameter *min.pct=0.4* in order to observe more subtle differences between similar cell types. Genes with a log fold change greater than 0.75 and a *P*- value of less than 0.01 were considered cell-type markers. All the *P*-values were adjusted in Seurat using Bonferroni correction. The lineage and cell type differentially expressed markers are reported in **Supplementary Table 3**. To generate the volcano plot in **Figure 2d**, we compared all the genes between macrophage populations in early and late time points using Wilcoxon Rank Sum test followed by Benjamini-Hochberg procedure, the results are reported in **Supplementary Table 4.**

### Education trajectory

We used the diffusion map ^131^, a non-linear dimensionality reduction technique, to capture the continuous transitions (aka pseudotime) during macrophage education in the scRNA-seq data. Specifically, we first identified the differentially expressed markers of CSF- 1-treated (*Itgax*; CD11c+) and educated (late) non-cycling macrophage clusters as described in the *Differential expression* section but using the *MAST* method ^132^. We then used the scaled and centered macrophage scRNA-seq data of these 74 marker genes (Supplementary Table 4) as input for the *DiffusionMap* function in the *destiny v2.6.2* R package ^133^ with default parameters. The first two eigenvectors (DM1 and DM2) captured the continuous transition between CSF-1- treated and educated macrophages. Therefore we used them to represent the macrophage education trajectory. Then we fitted a principal curve ^134^ using the *princurve v2.1.4* R package to the education trajectory (DM1 and DM2). We defined the education pseudotime as the projection of each cell onto this principal curve (the arc-length from the beginning of the curve) and normalized the pseudotime into 0-1 range as described previously ^135^. To identify the gene expression dynamics, we first smoothed the gene expression profiles of the 1,000 highly variable genes as described above on the education pseudotime by fitting a local polynomial regression, using *loess* R function with the smoothing parameter *span=0.7.* We then calculated the mutual information between the smoothed gene expression profiles and the pseudotime using the *discretize* and *mutinformation* functions in the *infotheo v1.2.0* R package ^136^ with default parameters. We filtered the result by removing the genes that have lower mutual information than the 25% quantile of the total mutual information calculated and retained 750 genes (reported in Supplementary Table 4). We applied K-means clustering using the *kmeans* R function to cluster these gene expression dynamics into 3 clusters. The 3 clusters of genes corresponded to CSF-1-treated, transient (early day 2) and educated (late day 10) macrophage signatures according to their expression dynamics on the education pseudotime. To visualize the gene expression dynamics during macrophage education as shown in Figure 2, we sorted the single cell transcriptomes and split them into a number of equal-width bins (n = 20 in Figure 2c; n = 50 in Figure 2e) on the education pseudotime, and average expression profiles were calculated within each pseudotime bin.

### Gene signature scoring

For a given gene set, we calculated gene signature score in single cells as described previously ^137^. We used the *AddModuleScore* function in Seurat to calculate the gene signature score. Briefly, signature genes were split into 10 bins based on their average expression levels. For each gene, 100 control genes were selected at random within the same expression bin to serve as control sets. The gene signature score was calculated as the differences between the aggregated expression of signature genes and the controls.

### Human breast cancer scRNA-seq

Human breast cancer scRNA-seq data were obtained from GSE114725 ^16^. We used the Final Annotation based on bulk combined with differential expressed genes as described by the authors ^16^ to select all the monocytic populations including macrophages, monocytes, monocyte precursors, plasmacytoid dendritic cells (pDCs), and myeloid DCs. We normalized the count data to logTPM as described in the *Preprocessing* section. We defined the *ex vivo* education signature as differentially expressed genes between CSF-1-treated (CD11c+) and educated (late) non-cycling macrophages using Wilcoxon Rank Sum test implemented in Seurat’s *FindAllMarkers* function with parameter *min.pct=0.4*. Genes with a log fold change greater than 0.75 and a Bonferroni corrected *P*-value of less than 0.01 were selected. Similarly, we defined human M2-signature by performing differential expression analysis between the three TAM clusters (23, 25, 28) with other monocytic populations using the same procedure. To qualitatively visualize the data, we performed the Principal Component Analysis using only the *ex vivo* murine signature as shown in **Extended Data Figure 13a** (middle panel). We found the TAMs and other monocytic populations could be largely separated by the first PC using our *ex vivo* signature. We then calculated the gene signature score using the *ex vivo* signatures in this human breast cancer scRNA-seq data as described in the previous section. Differential expression analysis between cluster 28 and the other myeloid cells was conducted across the entire transcriptome using the Wilcoxon Rank Sum test implemented in Seurat’s *FindAllMarkers* function with FDR corrections, and the markers are reported in **Supplementary Table 11**.

### MMTV-PyMT scRNA-seq

We processed the *in vivo* scRNA-seq data using Seurat V3 in a similar way as described above. The original data contain a fraction of spiked-in cells from human breast cancer cell lines that were sequenced together and removed analytically upon clustering (Supplementary Note MMTV-PyMT Macrophage scRNA-seq). In total, we sequenced 1,424 cells from the MMTV-PyMT mouse model. Briefly, we used the *FindVariableFeatures* function with first the ‘vs.t’ method to identify 2,000 highly variable genes, and then performed tSNE dimension reduction and clustering (*resolution=0.2*) using the top 20 PCs of cells. We identified 7 distinct cell types including 2 tumor epithelial subtypes, 2 macrophage subpopulations (Mac-1 and Mac-2), MDSCs, T/NK cells and red blood cells. Only immune cells were used for downstream analysis. The cell type annotation and metadata can be found in Supplementary Table 7. To identify the differentially expressed genes between Mac-1/SAM and Mac-2/TNM, we used the Wilcoxon Rank Sum test implemented in Seurat’s *FindAllMarkers* function with FDR and Bonferroni corrections (Supplementary Table 7). The SAM survival signature genes (n=38) used for patient survival analysis were selected as genes with log fold change > 0.75 and Bonferroni adjusted *P*-value < 0.05, and the TNM survival signature genes (n=71) were selected as genes with log fold change < -0.75 and Bonferroni adjusted *P*-value < 0.05. To calculate survival differences, human homologs of the corresponding mouse genes were used, and are reported in Supplementary Table 7.

## SPOTS Analysis

### Preprocessing

tumor tissues were first aligned onto the Visium slides with Loupe Browser 5.0 (10x Genomics) using immunofluorescent images as input. The resulting alignment JSON files were used as input for Space Ranger software v.1.3.0 (10x Genomics), and the transcriptomic reads were mapped to a custom genome (mm10 plus PyMT transgene). The ADT reads were mapped to the antibody barcode sequences (**Supplementary Table 8**) with the 4,992 whitelist tissue spatial barcodes provided by 10x Genomics as input (https://kb.10xgenomics.com/hc/en-us/articles/360041426992-Where-can-I-find-the-Space-Ranger-barcode-whitelist-and-their-coordinates-on-the-slide-) using the Python package CITE-seq-Count v1.4.3 ^138^. The count matrices of the transcriptome and ADT data were then normalized using log-transformed transcript per million (logTPM) as described in the **Single Cell RNA-seq Analysis** section.

### Visualization

To better visualize the sparse ST data, we utilized local Gi* statistics (^139^), which is similar to local Moran’s I and local Geary’s C statistics (^140^), to create smoother expression profiles for visualization and diagnostics. The local Gi* statistics for each feature/gene was defined as 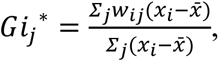 where *x_j_* is the expression value of a feature in cell *j* and *w_ij_* is the spatial weight between cell *i* and *j*, the 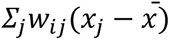 term is interpreted as weighted neighborhood or spatial lag. By definition local Gi* statistics is a Z-score and its *P*-value can be calculated using mathematical derivations provided in ^141^ or permutation-based methods ^140^. Because Visium tissue barcodes are arranged on hexagonal lattices, we defined the weight matrix using hexagonal nearest neighbor distance (HNN) ^142^ which reflects its true spatial orientation. We assigned the weights using HNN with a Gaussian kernel of width 1 unit distance, and only neighbors within 3 unit distance were considered.

### Deconvolution

We performed deconvolution of the spatial transcriptomics data using SPOTlight R package ^143^ with default parameters, except we set *min_cont=0* which didn’t remove any cells, and a recent unbiased scRNA-seq data of wild type MMTV-PyMT mouse model ^144^ as reference. Briefly, the scRNA-seq data were downloaded from GSE158677 and analyzed using Seurat 4.0 (^95^). Upon clustering, we identified six major cell types including tumor, CAF, Endothelial, Myeloid, B and T/NK cells. Differentially expressed genes were calculated by Seurat’s *FindAllMarkers* function with parameters *min.pct=0.5* and *logfc.threshold=1*, and used for deconvolution with SPOTlight’s *spotlight_deconvolution* function. For the ADT deconvolution, we first scaled the cell type ADTs (tumor: EpCAM and KIT; CAF: PDPN; Macrophage: CD11c, CD86 and F4/80; B cell: CD19 and B220; T/NK: CD4, CD8 and NK-1.1) using cosine normalization (^145^) and calculated the mean expression of their ADTs for each cell type. The deconvoluted percentages were calculated as the fraction of the cell typic ADT expressions over the total sum of all ADTs. The scaled deconvolution values were calculated using local Gi* statistics as described above. The IF staining intensity was quantified for each spatial barcode and normalized to the 0-1 range, outlier values below 1st or above 99th percentiles were removed. The deconvolution values are listed in **Supplementary Table 9**.

### Integration and Clustering

We followed Seurat’s tutorial for data integration and clustering. For both transcriptome and ADT data of the three tumor replicates, we first conducted Principal Component Analysis with the *RunPCA* function and used the *FindIntegrationAnchors* function to find a set of integration anchors. Then we used the *IntegrateData* function to integrate the three tumor replicates based on the anchors found. The UMAP dimensional reduction (^146^) was performed with the first 20 PCs. The clustering was done using *FindNeighbors* and *FindClusters* functions with the first 20 PCs and parameter *resolution = 0.5*. Overall, 15 integrated transcriptome-base clusters and 11 ADT-based clusters were found. Subsequently, we used the first 20 PCs to calculate the *K* nearest neighbor for each cell, where *K* is set to 5% of the total cell number. The batch effect was defined as the difference between replicate frequencies in the nearest neighbors and its theoretical values when the data are well mixed.

### Modality Weights

In order to integrate the transcriptome and ADT data and assess the contributions of each modality, we adopted a bootstrap-based approach that is similar to Jackknife ^147^ or Jackstraw ^148^. First, the data matrix was normalized and Principal Component Analysis was conducted either with top highly variable features or all the features. Second, a continuous *K* nearest neighbor graph was calculated using a Gaussian kernel with adaptive kernel width ^149^. Then we iterate for *N* times the previous steps, and each time randomly subsample a fraction (*Q*) of the features. The norms between these bootstrapped *K* nearest neighbor graphs and the original one served as weighted local robustness (WLR) measurements of each modality in determining the data manifold for every cell. Then, we simply performed the same procedures for each modality and the ratio between them represented the contribution for each modality. Data integration can also be done on the fused nearest neighbor graph using WLR as weights. For transcriptome data, we used the top 5,000 highly variable features and the first 20 PCs as input, while for the ADT data (32 features) we omitted the PCA step. The number of neighbors was set to 10% of the total cells and the subsample fraction was set to 10% of the total features used. In total 100 iterations were run and the L1 norm was used to calculate the WLR weights. The WNN weights were calculated using Seurat’s *FindMultiModalNeighbors* function with first 30 PCs of transcriptome and first 10 PCs of ADTs.

### Spatial Correlation

The spatial correlation of each gene can be defined by either univariate or bivariate Moran’s I. In the univariate case, Moran’s I of any feature *X* was calculated as *I* = 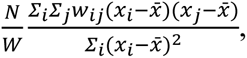 where *w_ij_* is a spatial weight matrix with *w_ii_* = 0 and *W* = *Lw_ij_*, and *N* is the number of total spatial barcodes. The *P*-value of univariate Moran’s I can be obtained by using mathematical derivations provided by ^150^. In the bivariate case, Moran’s I between feature *X* and *Y* was calculated as 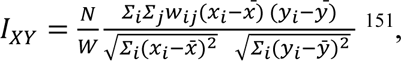 ^151^, and the *P*-value can be obtained by using mathematical derivations provided by ^152^.

### Spatial Component Clustering

To identify both transcriptionally and spatially distinct cell populations, we modified the current clustering scheme, which is solely based on expression data, by incorporating the spatial components of each spatial barcode. We adopted a method that has been developed in the field of spatial statistics by ^151^, where multivariate spatial correlation (MSC) was defined as 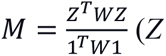 (*Z* is the z-score of the data matrix). This *M* matrix captures the spatial relationship between each gene across tissue spatial barcode and was used for subsequent principle component analysis. This approach dictated that the spatial correlation matrix (*M*) must be positive semidefinite, like the covariance matrix used in Principal Component Analysis (PCA), where the eigenvalues must be nonnegative. However, the *M* matrix is not positive semidefinite and can have negative eigenvalues as noted in the original paper. To solve this, another spatial correlation matrix *L* has been proposed by ^153^ with 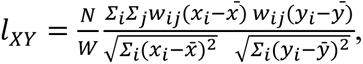, such that 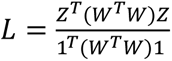 is positive semidefinite. By using this L matrix, we performed the singular value decomposition (SVD) in the same manner as PCA. We termed this approach the Spatial Component Analysis (SCA) to imply its connection to canonical PCA. For clustering of the ADT data, we used the HNN distance weights as described in the **Visualization** section, and first 10 Spatial Components (SCs) as input to build the nearest neighbor graph for clustering as described in the **Integration and Clustering** section. We merged the spatial clusters across different samples using hierarchical clustering (*hclust* and *cutree* functions in R with parameter *h=0.55*) based on their intercluster correlations and obtained 11 spatial clusters across three tumor tissues. The clustering result is reported in **Supplementary Table 10**.

### Differential Expression Analysis

Differentially expressed (DE) genes for each spatial cluster were calculated using Seurat’s *FindAllMarkers* function with default parameters or *logfc.threshold = 0.1* for ADT data. For DE genes between SAM (cluster 2) and TNM (cluster 5) regions, the top 5,000 variable genes with highest dispersion values were selected using *FindVariableFeatures* function with ‘*dispersion’* method. Gene Set Enrichment Analysis (GSEA) was performed using *fgsea v1.16* R package ^154^ with 10,000 permutations. Notch signaling pathway genes (n=179) were selected from corresponding GO terms (GO:0007219) with annotation term “Notch signaling pathway”. The result of this analysis is reported in **Supplementary Table 10**.

### Patient survival analysis

The METABRIC ^100^ microarray mRNA expression data (n=1,904) were obtained from cBioPortal (https://www.cbioportal.org/study/summary?id=brca_metabric) together with the corresponding clinical information including ER/PR/HER2 status, PAM50 subtypes, neoplasm histologic grade, inferred menopausal state, and survival information, including overall survival (OS) and relapse free survival (RFS) time (month), survival (living/deceased) and relapse free (recurred/not recurred) status. Of note, the disease specific survival status (DSS) is not provided by cBioPortal at the time when the analysis was performed, instead we obtained it from Synapse (https://www.synapse.org/#!Synapse:syn1757055; n=1,895). We took the average expression of SAM/Mac-1 and TNM/Mac-2 signature genes as defined above, to calculate the signature expression levels. To remove any bias due to macrophage infiltration differences across tumor samples, we further divided the expression of the signature genes by the pan-macrophage marker CSF1R. We then grouped the patients based on their signature expressions, and used the lower (<25^th^ percentile) and upper (>75^th^ percentile) quartiles to stratify them into high and low expression groups. Cox proportional hazard regression model, implemented by *survival* R package, was used for survival analysis to correct for ER (+/-), PR (+/-), HER2 (+/-), tumor grade (I, II or III) and menopausal (pre/post) status. Kaplan-Meier survival curves were generated by using *survminer* R package. The univariate and multivariate Cox proportional hazard regression models of OS, DSS and RFS are reported in **Supplementary Table 11**.

## Supplementary Movies and Tables

**Supplementary Movie 1:** Live imaging of mT-labeled BMDMs (Rosa26^mTmG^ mouse) plated with oTME culture. Related to **Extended Data Figure 1f**.

**Supplementary Movie 2:** Live imaging of mT-labeled monocyte-derived macrophages (*Rosa26^mTmG^* mouse) differentiated with tumor epithelial cells (TNMs). Note the distinct dendritic morphology of TNMs. Related to Figure 6a.

**Supplementary Movie 3:** Live imaging of mT-labeled monocyte-derived macrophages (*Rosa26^mTmG^*mouse) differentiated with PDGFRA+CD24^neg^ stromal cells (SAMs), highlighting their increased size and granularity. Related to Figure 6a.

**Supplementary Table 1.** Bulk RNA-seq analysis of CSF-1-treated (MO) and oTME-educated (CO) macrophages. Related to Figure 1. Table with differentially expressed marker genes of CSF- 1-treated and educated macrophages using DESeq2 analytical pipeline.

**Supplementary Table 2.** Differentially expressed marker genes between non-cycling CSF-1- treated (CD11c+) (Tab1) and 10-day educated (Late) macrophage clusters (Tab2). Related to methods. Table with marker genes for CSF-1-treated and educated macrophages calculated using MAST method within the Seurat analytical pipeline.

**Supplementary Table 3.** Cell type markers of oTME culture. Table of differentially expressed genes for, between, and within oTME cell lineages as calculated using MAST method within the Seurat analytical pipeline.

**Supplementary Table 4.** Tab 1: Clustering of dynamically expressed genes in the education trajectory of macrophages. Table with 750 genes assigned to three macrophage clusters (CSF-1- treated, educated early, and educated late) using K-means clustering. Tab 2: Differentially expressed genes between non-cycling early (day-2) vs. late (day-10) educated macrophages. Table with marker genes calculated using Wilcoxon Rank Sum test followed by Benjamini-Hochberg procedure within Seurat analytical pipeline.

**Supplementary Table 5.** tumor characteristics of the human breast cancer cohort.

**Supplementary Table 6.** Differentially expressed marker genes between SAM/Mac-1 and TNM/Mac2 *in vivo*.

**Supplementary Table 7**. Cell type annotations and metadata for oTME (Tab 1) and TME MMTV- PyMT scRNA-seq. (Related to **Extended Data Figures 2b and 8d**). Tables with description of cell types, sample metadata and summary statistics for single cell transcriptome used in this study.

**Supplementary Table 8.** 32 ADTs (Biolegend Total-seq-A) used in SPOTS experiments. Related to Figure 6.

**Supplementary Table 9**. Raw and scaled spatial deconvolution values of the transcriptomic and ADT data. Related to Figure 6 **and Extended Data Figure 10**.

**Supplementary Table 10.** Tab 1: Spatial clustering results from SPOTS analysis. Related to Figure 6. Tab 2: Differentially expressed ADTs for each spatial cluster. Related to Figure 6. Tab 3: Differentially expressed mRNAs for each spatial cluster. Related to **Extended Data Figure 10**. Tab 4: Differentially expressed marker genes between SAMs (Cluster 2) and TNMs (Cluster 5) enriched regions. Related to Figure 6.

**Supplementary Table 11**. Tab 1: Differentially expressed marker genes between human SAM (Cluster 28) and other myeloid cells. Related to **Extended Data Figure 10a**. Tab 2: SAM/Mac-1 and TNM/Mac-2 gene expression signatures used for survival analysis. Related to **Extended Data Figure 10b**. Tab 3: The multivariate Cox proportional hazard regression models of OS, DSS and RFS in METABRIC cohort. Related to Figure 7.

**Supplementary-Note-oTME-Macrophage-scRNA-seq-Analysis:** R markdown output with code to reproduce the oTME scRNA-seq results. Related to **Figures 2, Extended Data Figure 2-3** and **methods (‘Single Cell RNA-seq Analysis’ section**): https://drive.google.com/file/d/1A7rDgX3jWdepfS8BcgYRhd8LoiRWEC_O/view?usp=sharing

**Supplementary-Note-MMTV-PyMT-Macrophage-scRNA-seq-Analysis:** R markdown output with code to reproduce the MMTV-PyMT scRNA-seq results. Related to Figures 5**, Extended Data Figure 7** and methods (**‘MMTV-PyMT scRNA-seq’ section**): https://drive.google.com/file/d/1KzFg_lm-oOEOi7DiXnxWlgCYFXGeI36i/view?usp=sharing

---

**Extended Data Figure 1.**
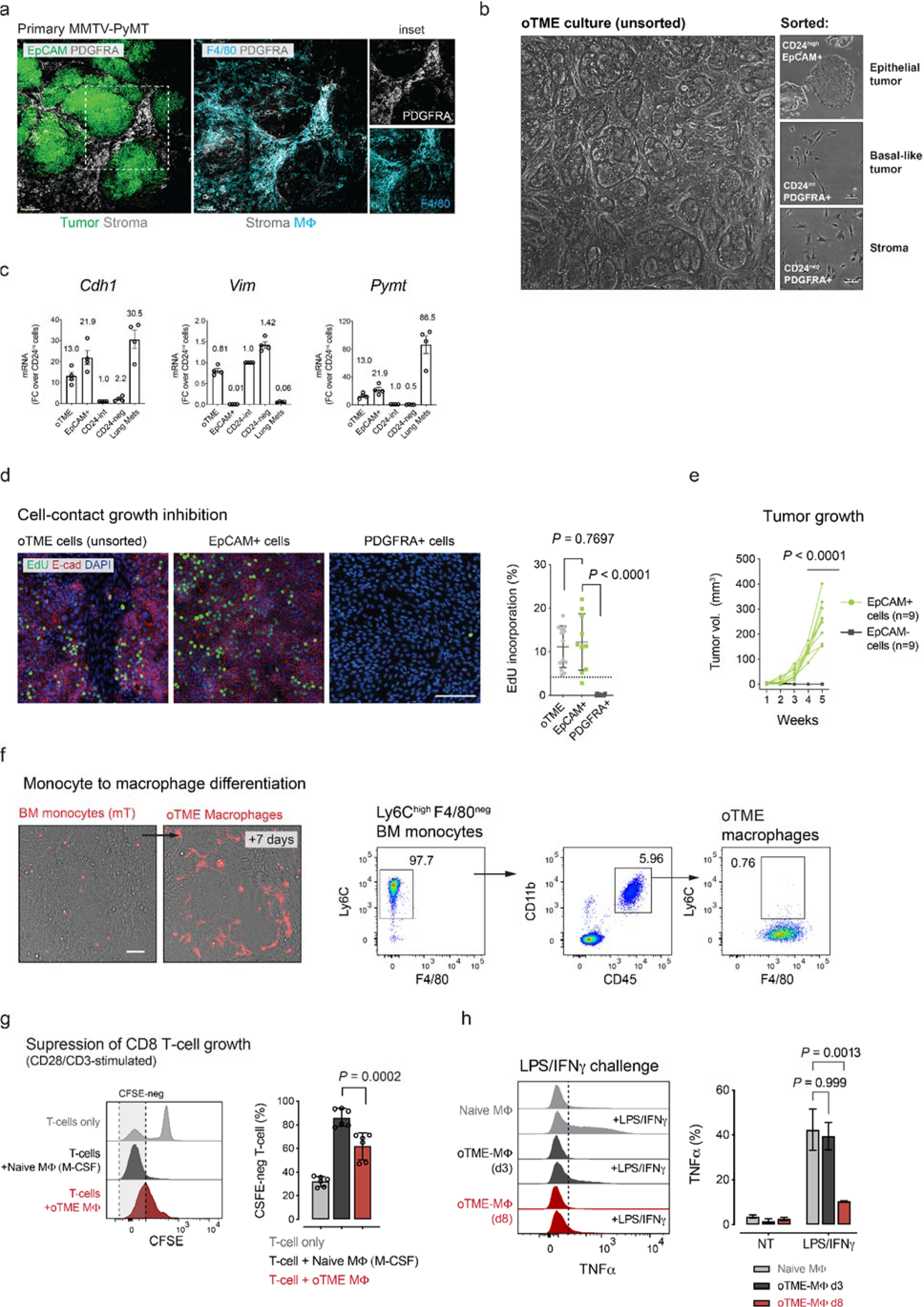
Phenotypic characterization of organotypic TME (oTME) cellular components. (**a**) Immunofluorescence of EpCAM (tumor), PDGFRA (stroma), and F4/80 (macrophages) in intact tumor fragments from the MMTV-PyMT model (*in vivo)* demonstrating the typical cellular organization. Scale bars, 50μm. Representative images from 3 independent experiments. (**b**) Phase images of typical oTME culture along with its sorted components based on CD24, PDGFRA, and EpCAM expressions. (**c**) Representative qRT-PCR of E-Cadherin (*Cdh1*), Vimentin (*Vim*) and *Pymt* transgene expression, normalized to PDGFRA+CD24^int^ cells (n=4 replicates). Data are shown as mean±SE. (**d**) Contact-growth inhibition measured by EdU incorporation (1hr exposure) in confluent cultures of oTME, EpCAM+ and PDGFRA+ cells. Data (fields of view; oTME, n=13; EpCAM+ n=10; PDGFRA+ n=16) are shown as mean±SD, 2-way ANOVA test, Bonferroni-corrected. Scale bar, 100μm. (**e**) Tumor growth kinetics of orthotopic transplants in mammary glands generated either by EpCAM+ or EpCAM- cells (n=9 replicates). (**f**) *Ex vivo* differentiation of bone marrow (BM) Ly6C+ monocytes in the oTME model. Purified BM monocytes from *Rosa26^mTmG^* mice were cultured for 7 days in the presence of oTME cells and analyzed by flow cytometry for mT, CD45, Ly6C, CD11b, F4/80 expression. Representative data from at least 5 independent experiments. Images were taken at plating and 7 days later. Scale bar, 100μm. (**g**) CFSE proliferation assay in CD8 T-cells in the presence of oTME and CSF-1-treated BMDMs. Splenic T-cells were purified and stimulated with CD3/CD28 antibodies for 1hr, co-plated as indicated, and analyzed by flow cytometry 5 days later (n=6 replicates). Bars show mean±SD, 2- way ANOVA test, Bonferroni-corrected. (**h**) BMDMs were either treated with CSF-1 as control or added to the oTME cultures for 3, and 8 days and treated with LPS (20ng/mL) and IFNγ (10ng/mL) for 1hr. TNFα was measured by flow cytometry. Data are shown as mean±SD, 2-tailed unpaired Student’s t-test.

**Extended Data Figure 2:**
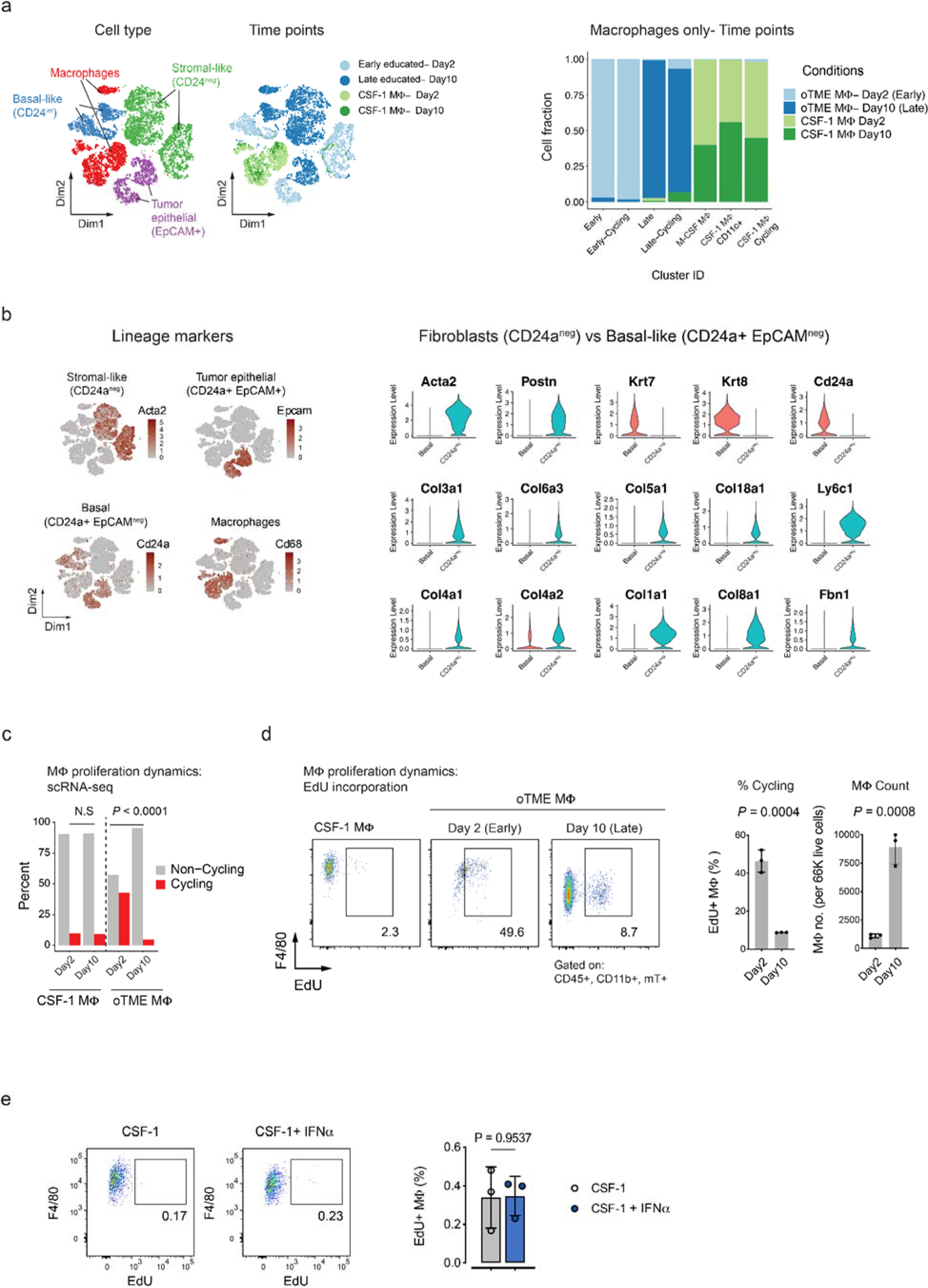
Analysis of proliferation dynamics in oTME macrophages. (**a) Left:** Two-dimensional t-SNE plots (Dim1 and Dim2) of annotated single-cell transcriptomes colored by cell type (left) and time point (right). **Right:** Stacked bar graph showing the composition (fraction) of each macrophage cluster according to the corresponding conditions. (**b) Left:** t-SNE plots of annotated single-cell transcriptomes and individual genes colored by cell lineage markers; *Acta2*+ (CD24^neg^PDGFRA+ stromal cells), *Epcam+* (tumor epithelial cells), *Cd24a*+ and *Epcam*- (basal-like tumor cells), and *Cd68* (macrophages). **Right:** violin plots showing expression of extracellular matrix (ECM) and lineage marker genes in the basal-like and CD24^neg^PDGFRA+ stromal cells (**Supplementary Table 2**). (**c**) Changes in relative proportions of cycling cells during macrophage education according to scRNA-seq clustering (n=1,866 CSF-1-treated Day 2; n=1,705 CSF-1-treated Day 10; n=145 Educated Day 2; n=552 Educated Day 10). Two-proportions Z-test, N.S. (not significant). (**d**) Cell cycle dynamics of oTME macrophages as depicted in (**c**) were validated using EdU incorporation (n=3 replicates); data are shown as mean±SD, 2-tailed unpaired Student’s t-test. (**e**) Tumor-naive BMDMs were stimulated with CSF-1 (10ng/mL) or CSF-1 and IFNɑ (10ng/mL each) and labeled with EdU for 48hrs. EdU incorporation was analyzed by flow cytometry (n=3 replicates). Data are shown as mean±SD, 2-tailed unpaired Student’s t-test.

**Extended Data Figure 3:**
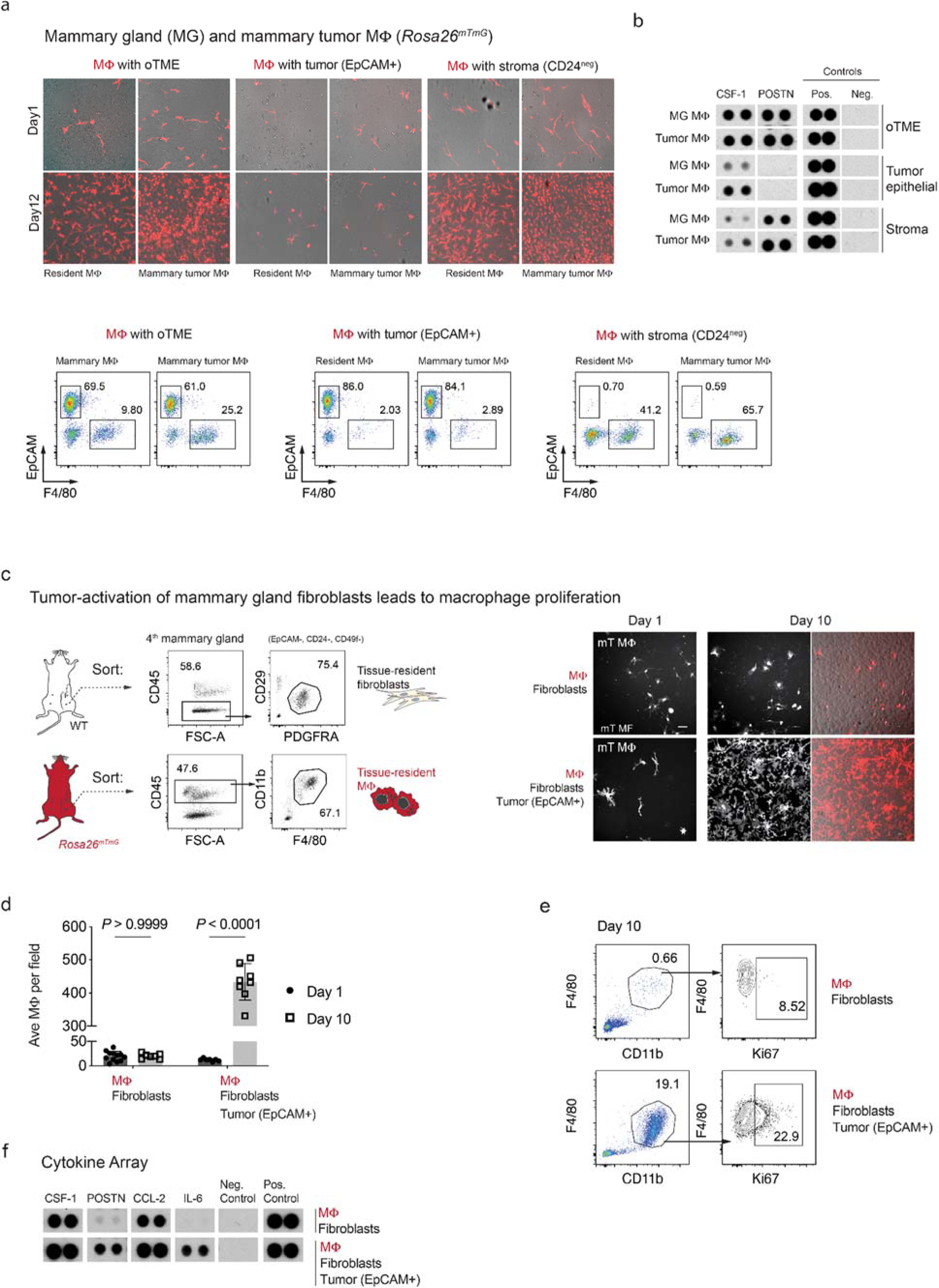
Cancer-associated fibroblasts but not resting fibroblasts promote macrophage proliferation. (**a**) Macrophages from healthy mammary gland or mammary tumors of *Rosa26^mTmG^* mice (n=5) were harvested and plated in the presence of the indicated cells. Fluorescent images of mT macrophages were taken on day-1 and day-12 and analyzed by flow cytometry using CD45, EpCAM, CD11b and F4/80. (**b**) Cytokine expression analysis of supernatants collected from (**a**). Growth media were conditioned for 4 days and probed for 111 cytokines using cytokine array. CSF-1, and POSTN are presented (**c**) Left, experimental design. Macrophages from *Rosa26^mTmG^* mice (n=5) and fibroblasts from WT mice (n=5) were FACS-sorted from the 4th mammary gland (MG4) and plated with or without tumor epithelial cells for additional 10 days. Right, fluorescent images of mT macrophages were taken on day 1 and day 10. Scale bar, 50μm. (**d**) Cell number quantification of mT macrophages from cell cultures in (**c**). Dots show mean±SD, 2-way ANOVA test, Bonferroni-corrected. (**e**) Flow cytometry of mT macrophage from cell cultures in (**c**), using CD45, F4/80, CD11b, and Ki67 antibodies. (**f**) Cytokine arrays analysis of supernatants from the culture conditions in (**c**). Enhanced proliferation of macrophages was detected in the presence of tumor-activated fibroblasts marked by expression of POSTN and IL-6.

**Extended Data Figure 4:**
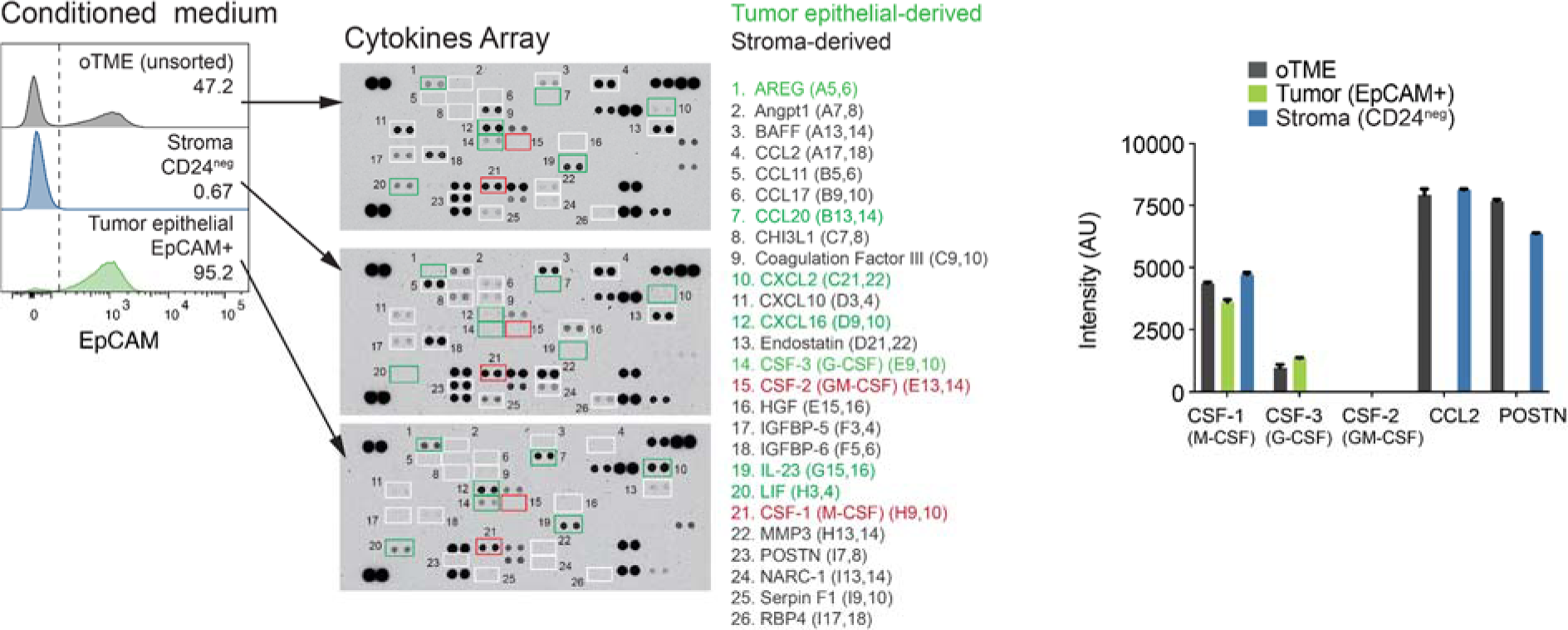
Cell-type specific cytokine expression profiles. Cytokine expression analysis of supernatants collected from sorted EpCAM+ tumor epithelial, CD24^neg^PDGFRA+, or unsorted oTME cells. Growth media were conditioned for 5 days from confluent cultures and probed for 111 cytokines. CSF-1 (M-CSF), CSF-3 (G-CSF), POSTN and CCL2 are presented in Fig. 3c and quantified along with the negative and positive control spots. Data are shown as mean±SD of signal intensity (arbitrary units) of technical replicates of the array.

**Extended Data Figure 5:**
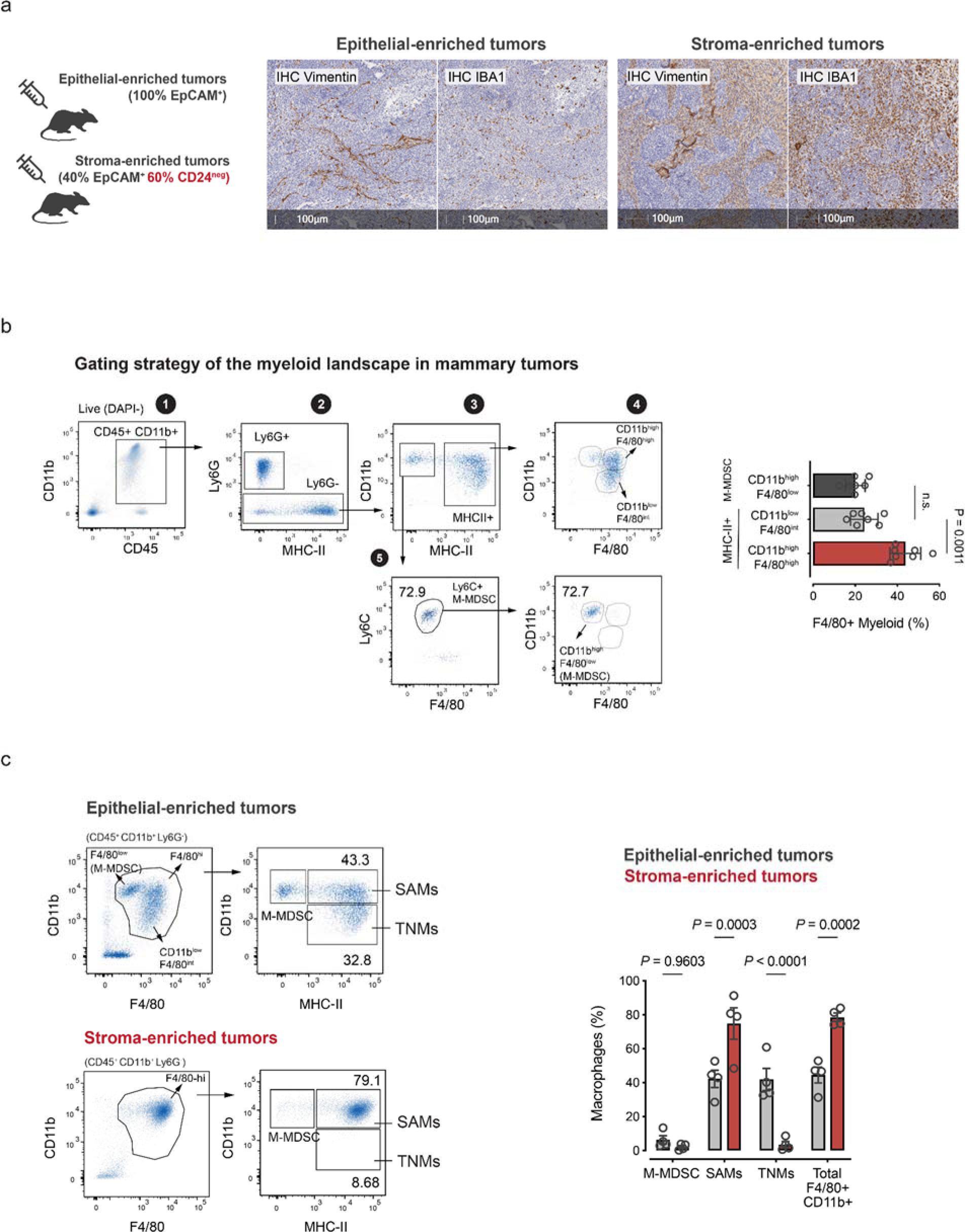
Immunophenotyping of macrophage populations (TNMs and SAMs) in murine mammary tumors. (**a**) Mammary tumor transplants originating from tumor epithelial cells (EpCAM+), or following enrichment with PDGFRA+CD24^neg^ stromal cells (n=4). IHC for vimentin (fibroblasts) and IBA1 (macrophages) in tumor transplants at study endpoint. Scale bars, 100μm. (**b**) Gating strategy for macrophages (SAMs/TNMs) in PyMT tumors. Myeloid cells were selected based on (**1**) CD45+ and CD11b+, followed by (**2**) exclusion of Ly6G+ granulocytes (**3**) and then classified into two subpopulations; (**4**) MHC-II^neg^Ly6C+F4/80^low^ (M-MDSC), and MHC-II+ macrophages that were further divided based on CD11b expression (**5**) Cd11b^low^F4/80^int^ (TNMs) and Cd11b^high^F4/80^high^ (SAMs). Relative cell proportions following this gating were quantified (n=7 replicates). Data are shown as mean ±SD, 2-way ANOVA test, Bonferroni-corrected. (**c**) Immunoprofiling of the changes in the myeloid landscape in mammary tumors from (**a**) and applying the gating strategy in (**b**). Three populations were presented: (i) Ly6C+MHC- II^neg^F4/80^low^ (**M-MDSC),** (ii) MHC-II^+^F4/80^high^CD11b^high^ (**SAMs**) and (iii) MHC- II^+^F4/80^int^CD11b^low^ (**TNMs).** Frequencies were calculated as the percentage of DAPI-CD45+ cells. Data are shown as mean±SD. (n=4 replicates), 2-way ANOVA test, Bonferroni-corrected.

**Extended Data Figure 6:**
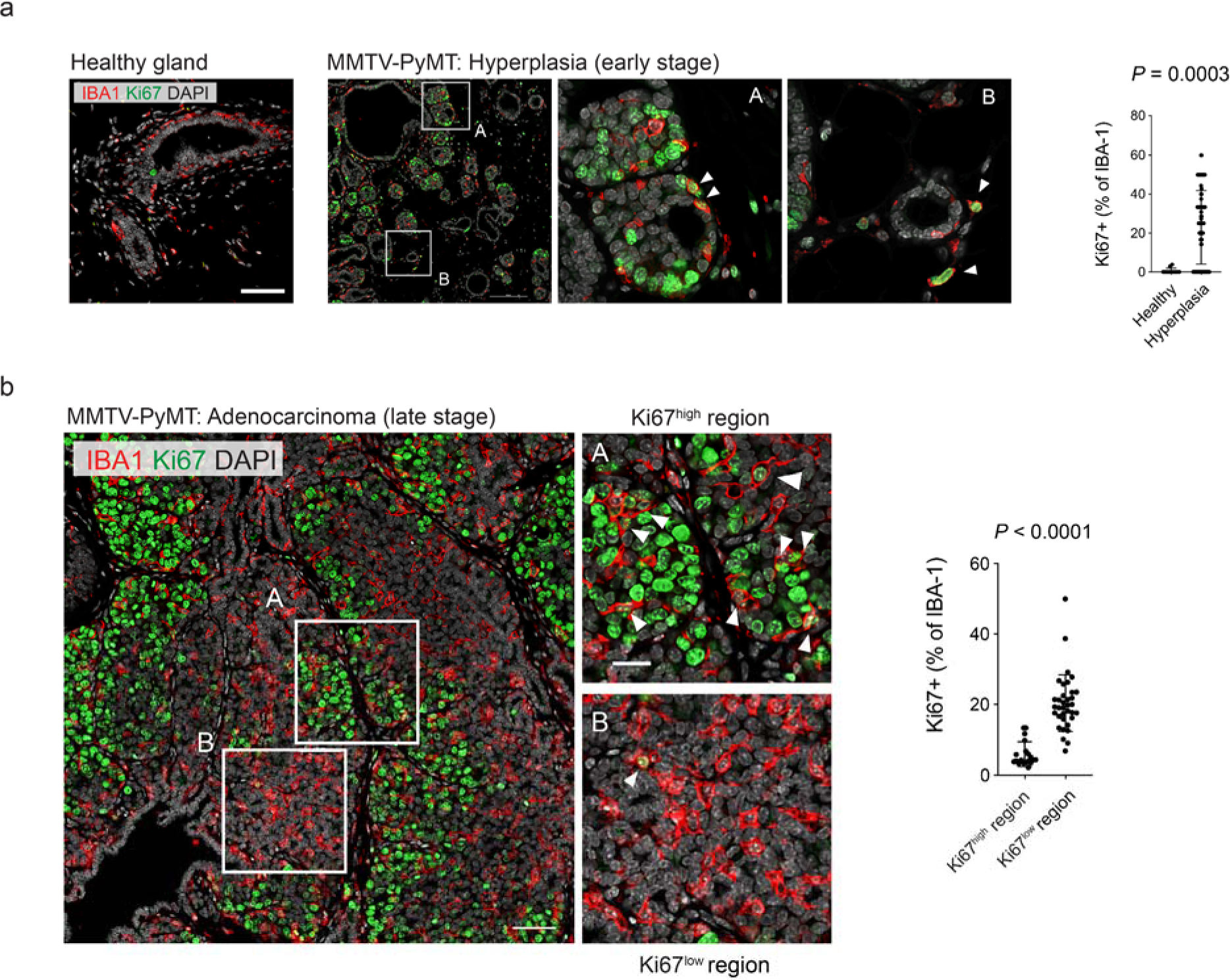
Macrophage proliferation in different stages of mammary gland tumorigenesis. (**a**) IF staining of IBA1 and Ki67 in mammary tissues from healthy (left) or hyperplasia lesions (right) of the MMTV-PyMT tumor model. Insets show intraepithelial (A) and stromal (B) macrophages. Dots represent fields of view (n=11 for healthy and n=92 for malignant) from 4 mammary tumors. Scale bars, 100μm. Data are shown as mean±SD, Mann–Whitney U test. (**b**) Immunofluorescence staining of IBA1 and Ki67 in late adenocarcinoma lesions of the spontaneous MMTV-PyMT model. Inset A; cycling regions (Ki67^high^), inset B; non-cycling regions (Ki67^low^). Dots represent fields of view from 6 tumors. Scale bars, 100μm. Data (fields of view; Non-cycling n=19, Cycling n=35) are shown as mean±SD, Mann-Whitney U test.

**Extended Data Figure 7:**
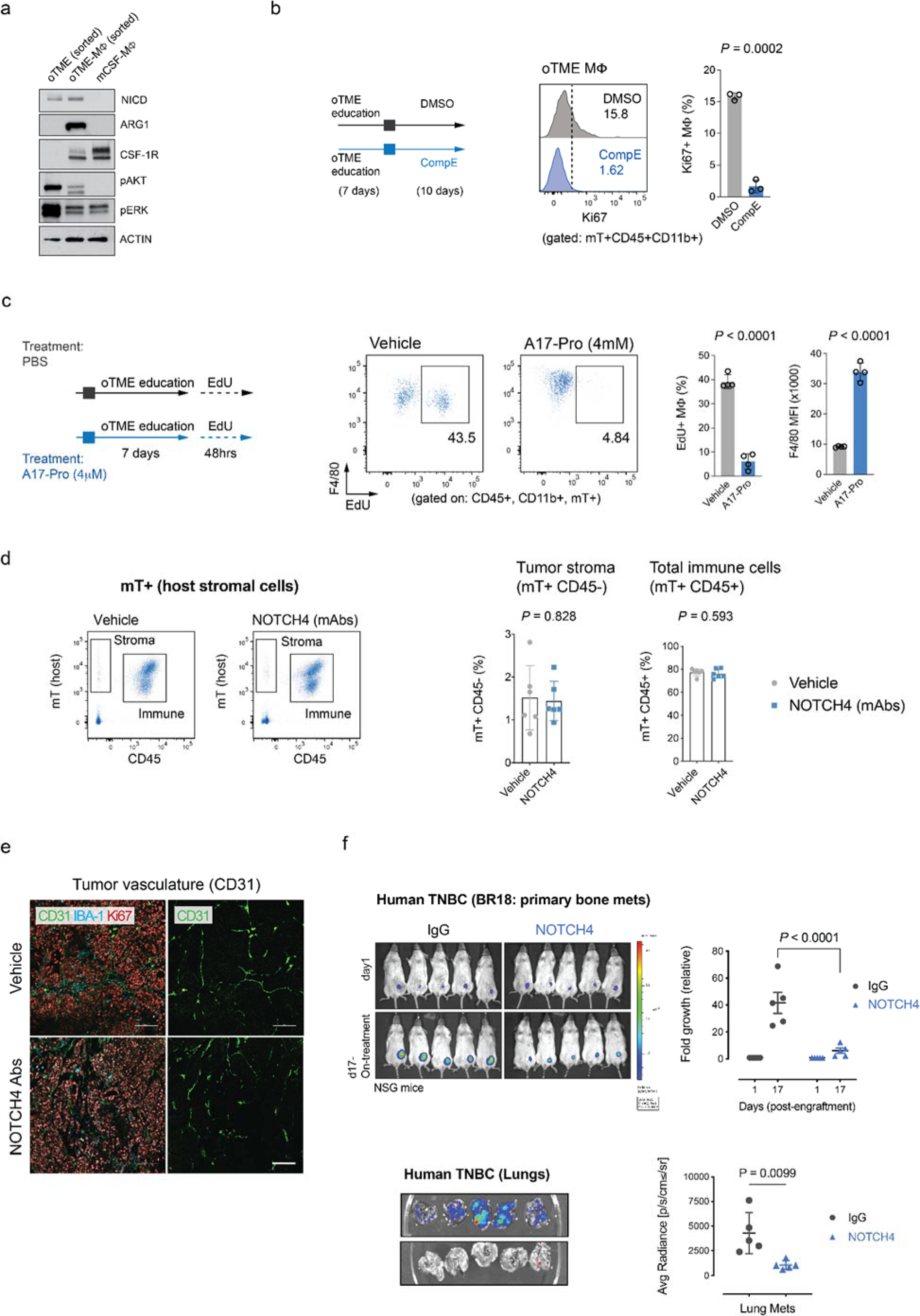
Impact of Notch inhibitions on macrophage proliferation and tumor growth of HGSOC cells. (**a**) Immunoblotting for the cleaved Notch intracellular domain (NICD), ARG1, phospho-PI3K (pAKT), and phospho-MAPK (pERK), confirming active Notch and PI3K pathways in oTME macrophages but not in CSF-1 macrophages. (**b**) Impact of γ-secretase inhibition on proliferation of oTME macrophages. BM monocytes were isolated and plated with oTME cells for 7 days and then were treated with either compound-E (CompE; 10μM) or DMSO as a control for additional 10 days. Macrophages and oTME cells were analyzed by flow cytometry for Ki67 expression. Data (n=3) are shown as mean±SD, Welch’s t-test. (**c**) EdU incorporation in BMDMs following treatment with Adam17 protease inhibitor (A17Pro) or PBS as vehicle control. Cells were plated with oTME cells in the presence of A17Pro or PBS and cultured for seven days. Cell cultures were then labeled with EdU for 48hrs and were analyzed by flow cytometry for EdU incorporation (n=4 replicates). Data are shown as mean±SD, Welch’s t-test. MFI; mean fluorescence intensity. (**d**) Flow cytometry of relative abundances of Ki67+ tumor cells (mT^neg^ CD45^neg^), total immune cells (mT+ CD45^+^), and stromal cells (mT+ CD45^neg^) in control vs. NOTCH4-treated tumors (n=6 replicates). Data are shown as mean ±SD, 2-way ANOVA test, Bonferroni-corrected. (**e**) Immunofluorescence staining of CD31, IBA1, and Ki67 in tumor transplants from (a). Scale bars, 100μm. (**f**) Primary tumor growth (left) and lung metastasis (right) of metastatic TNBC PDX cells (BR18) in NSG mice following NOTCH4 and IgG treatment. Mice were engrafted with 50K cells and treated as in (**c).** Growth kinetics and lung dissemination were evaluated by bioluminescence.

**Extended Data Figure 8:**
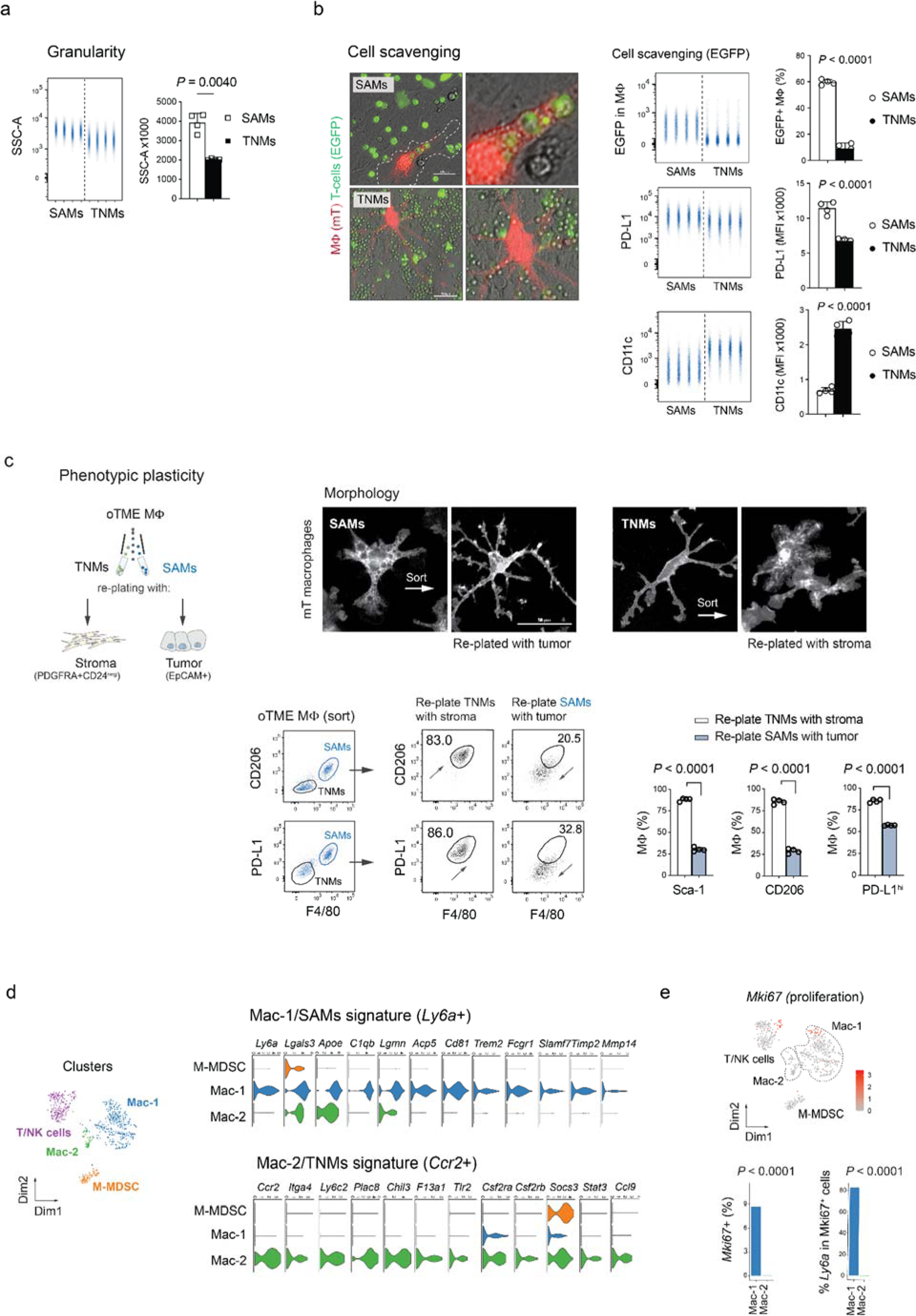
Functional characterization of macrophage plasticity in breast cancer. (**a**) Top, experiment design. Bottom, flow cytometry analysis of granularity (SSC) in monocyte-derived macrophages from *Rosa26^mTmG^* mice (n=4 replicates) that were differentiated in the presence of EpCAM+ tumor epithelial or PDGFRA+CD24^neg^ stromal cells. Data are shown as mean±SD. Student t-test. (**b**) Scavenging activity of SAMs vs. TNMs. Monocyte-derived macrophages were generated as in **a** and splenic EGFP+ T-cells from *Flt3^Cre^Rosa26^mTmG^* were purified and added for additional 5 days (**Methods**). Fluorescent images were taken (left), and flow analysis (right) was performed in mT+F4/80+CD11b+ macrophages for EGFP signal, PD-L1 and CD11c. Data (SAMs n=4; TNMs n=4) are shown as mean±SD, student t-test. Scale bars, 100μm. (**c**) Functional plasticity of oTME macrophages. Monocyte-derived oTME macrophages were sorted based on SAM (F4/80^hi^CD206+) or TNM (F4/80^low^CD206-) markers. SAMs were then replated with tumor epithelial cells and TNMs with PDGFRA+CD24^neg^ stromal cells. Seven days later, immunofluorescent images were taken, and cells were analyzed by flow cytometry with the indicated surface markers. (n=4; biological replicates). Data are shown as mean±SD, 2-way ANOVA test, Bonferroni-corrected. Scale bar, 50μm. (**d**) Violin plots from the scRNAseq analysis in Fig. 5g of key genes in Mac-1/SAMs (*Ly6a*^+^) and Mac-2/TNMs (*Ccr2*^+^) macrophage subpopulation. (**e**) Two-dimensional t-SNE plot (top) and barplots (bottom) of Mki67+ immune cells from scRNAseq analysis of MMTV-PyMT tumor. Right: Fractions of Mki67+ and Ly6a+ cells were quantified per cell type. The *P*-values were calculated by One-proportion Z-test with expected proportion of 10^-4^.

**Extended Data Figure 9:**
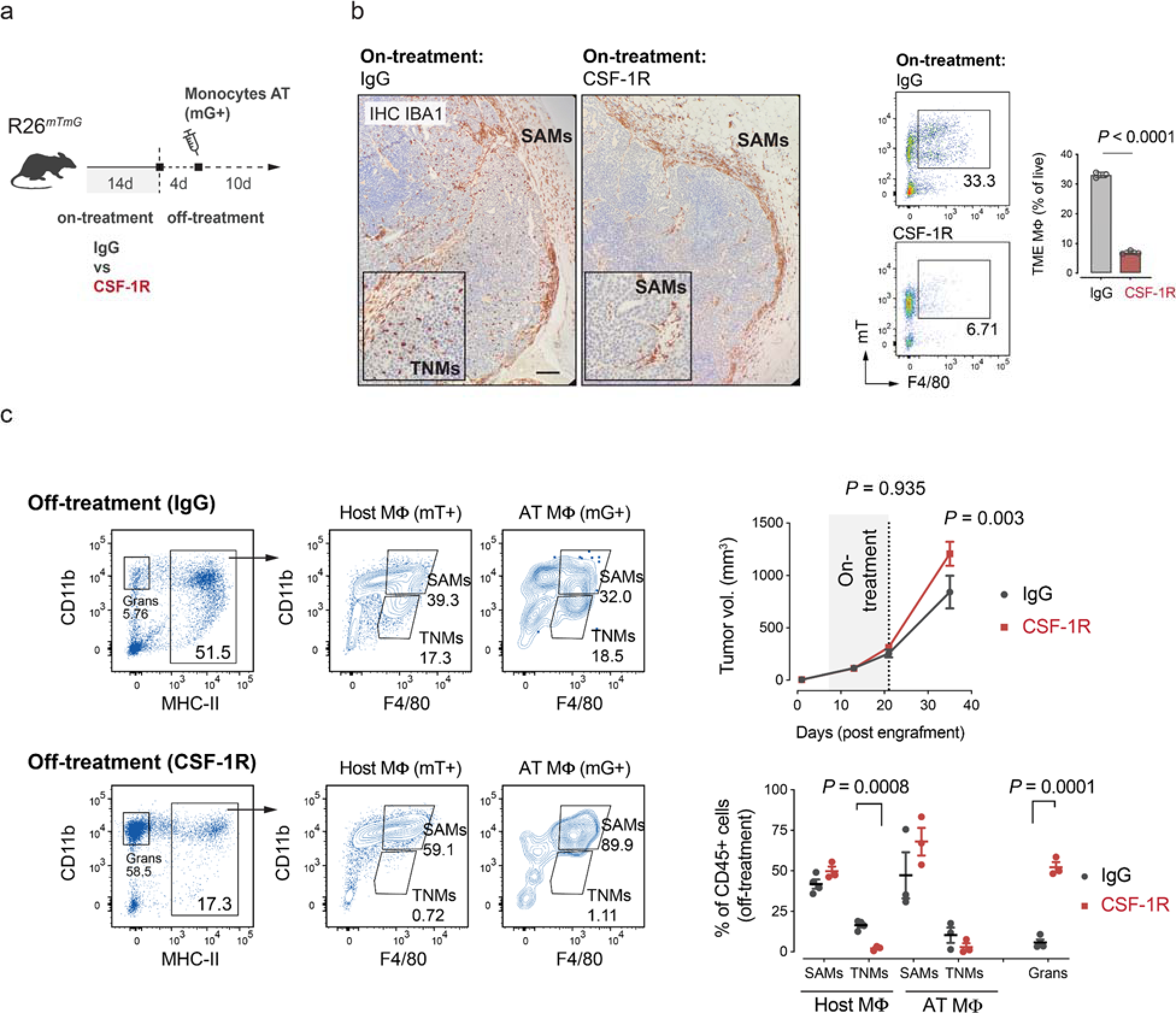
Spatial interactions in the TME override macrophage ontogeny and dictate macrophage immunophenotypes. (**a**) Experiment design. (**b**) IHC (IBA1) and flow cytometry analyses of mammary tumors following IgG or CSF-1R treatments. (**c**) Adoptive transfer (AT) of GFP+ (mG) BM monocytes (harvested from *Rosa26^mTmG^Flt3^Cre^* mice) into tumors in *Rosa26^mTmG^* mice following two weeks of CSF-1R or IgG treatments.

**Extended Data Figure 10:**
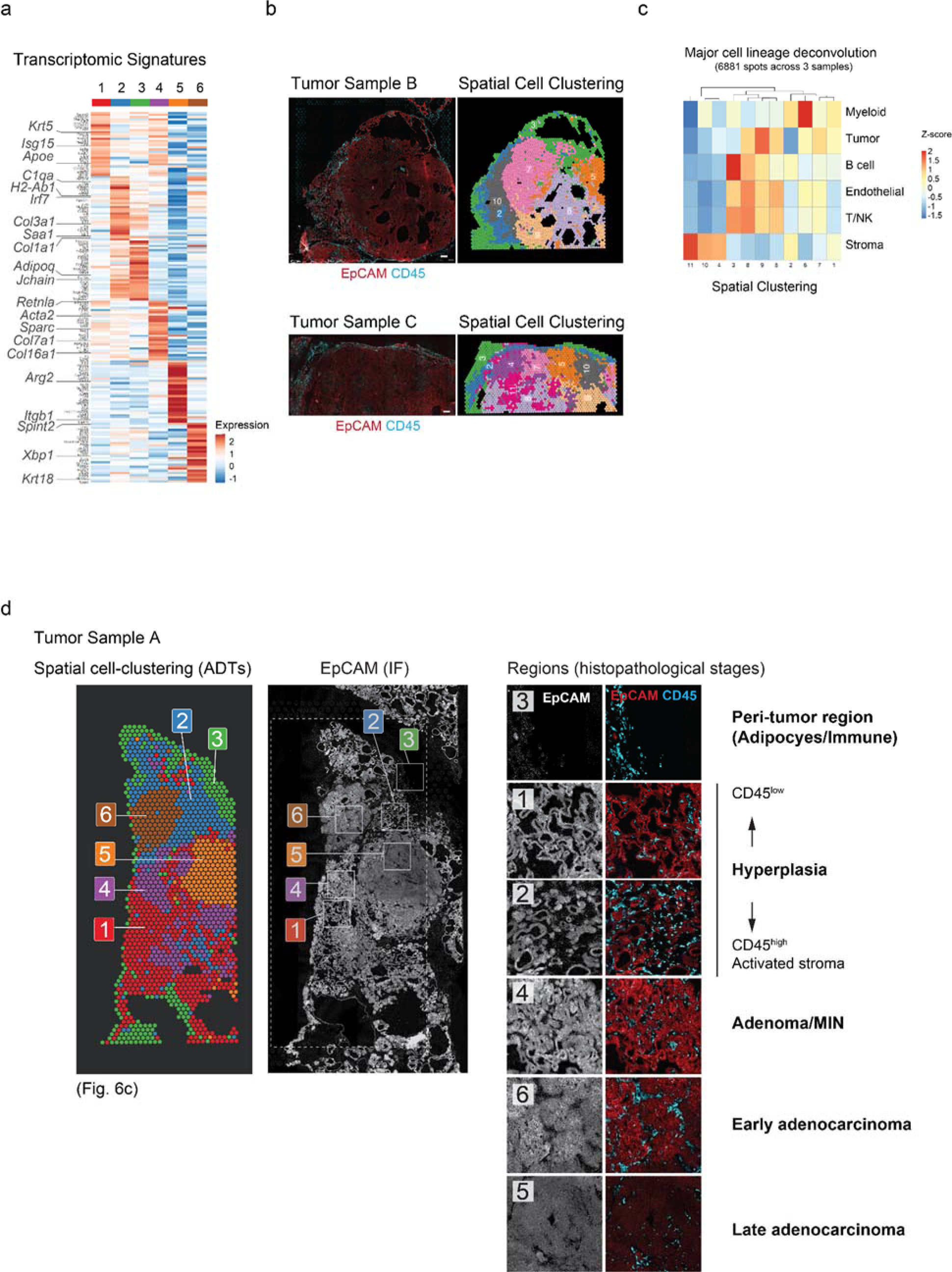
SPOTS clustering identified distinct microenvironments typical to the histopathological stages of mammary gland transformation. (**a**) Heatmap showing the differentially expressed genes (mRNA) for each of the spatial clusters from Fig. 6c. Key marker genes were highlighted. (**b**) IF staining of EpCAM and CD45 (left) and spatially-informed clustering (right) of tumor samples B and C. Scale bar, 200μm. (**c**) Relative ADT deconvolution values (Z-score) across all clusters. (**d**) Spatial clustering of tumor samples A (left; from Fig. 6c) along with the corresponding IF staining for EpCAM and CD45 (right). Insets display IF images in the indicated cluster regions along with the established histopathological staging of mammary gland transformation. Scale bar, 200μm. MIN, mammary intraepithelial neoplasia.

**Extended Data Figure 11:**
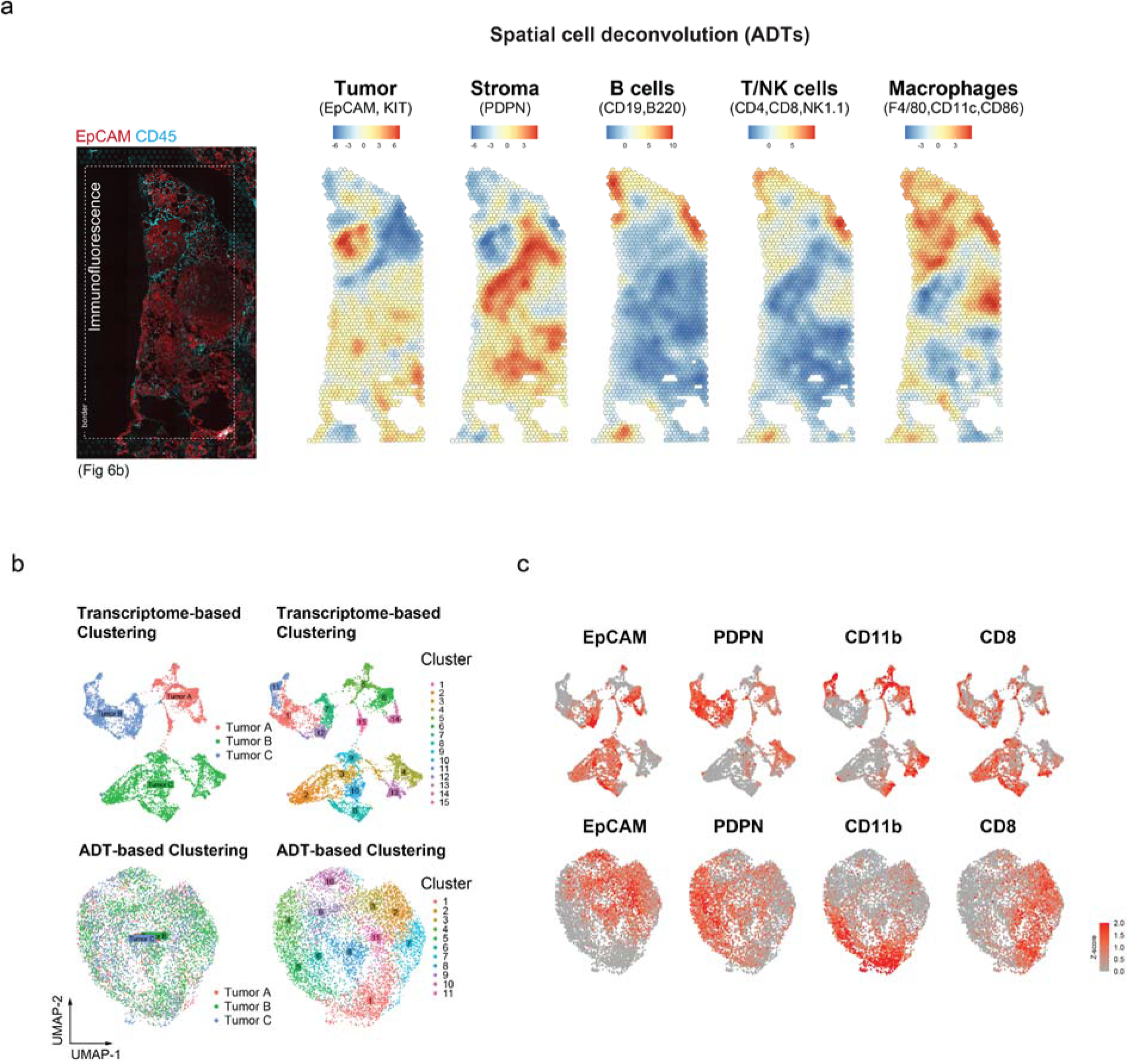
ADT vs. mRNA cell clustering results of three biological samples. (**a**) Spatial cell deconvolution of antibody-derived tag (ADT) data. Relative ADT deconvolution values of each spatial barcode (**Methods**) overlaid onto the breast cancer tissue to the left. Full deconvolution values are provided in **Supplementary Table 9**. (**b**) Two-dimensional UMAP plots showing the clustering results based on transcriptomic (top) or ADT (lower) data. Cells were colored by replicates (left) or clusters (right). (**c**) ADT expression levels (Z-score) of key cell type markers.

**Extended Data Figure 12:**
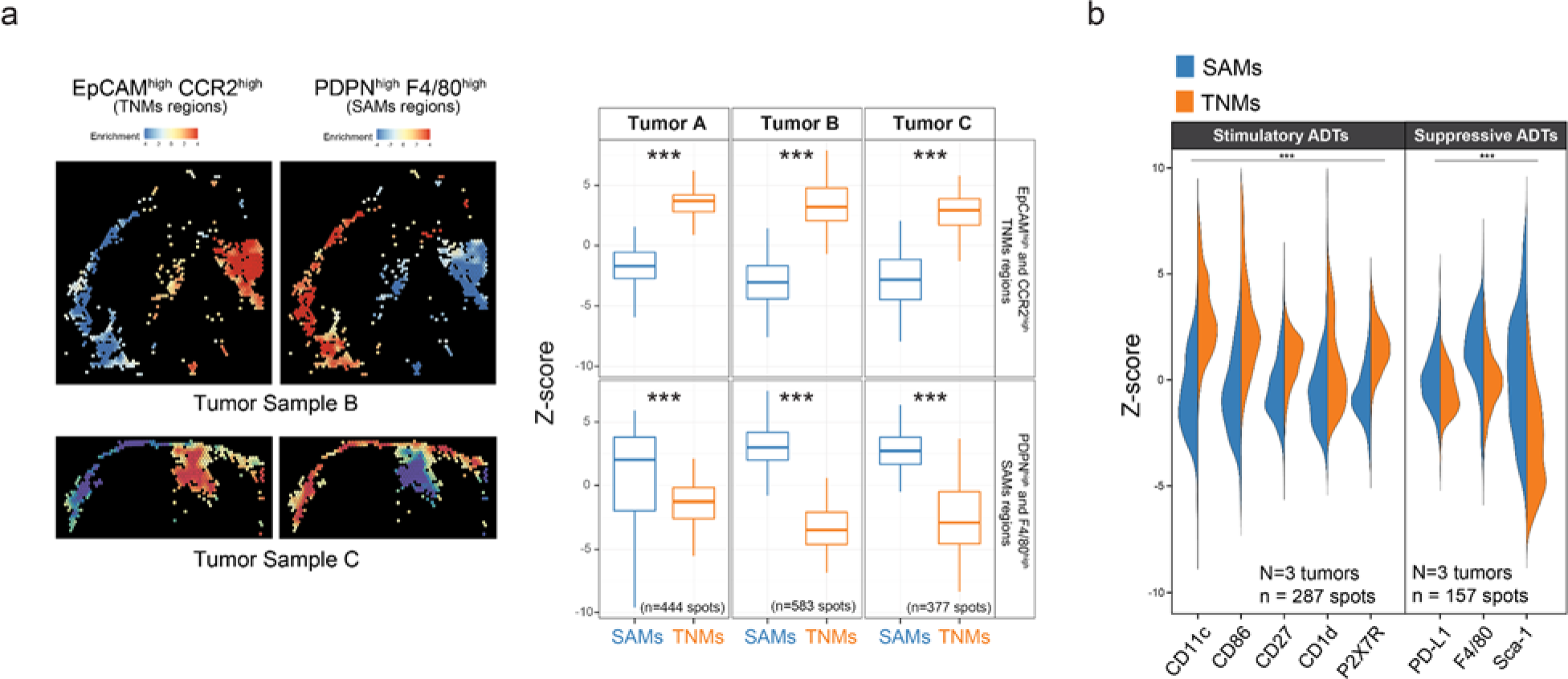
Co-expression of EpCAM and CCR2 ADTs in tumor-enriched regions and PDPN and F4/80 ADTs in fibroblast-enriched regions. (**a**) EpCAM and CCR2 vs. PDPN and F4/80 ADTs co-expression levels overlaid onto tumor sample B and C (sample A is in Fig. 6). Middle panel: boxplots of EpCAM and CCR2 vs. PDPN and F4/80 co-expression levels. Right panel: violin plots of immune-stimulatory and suppressive ADT expression levels. Kolmogorov–Smirnov test, \**P* < 0.05, \*\**P* < 0.01, \*\*\**P* < 0.001, otherwise not significant (n.s). Each boxplot ranges from the first and third quartiles with median shown as middle line, and the whiskers represent 1.5 times the interquartile range. (**b**) Violin plots of immune-stimulatory and immune-suppressive ADTs expression levels (Z-score) in fibroblast-enriched regions and tumor-enriched regions across all three tumor samples. Kolmogorov–Smirnov test, \**P* < 0.05, \*\**P* < 0.01, \*\*\**P* < 0.001, otherwise not significant (n.s).

**Extended Data Figure 13:**
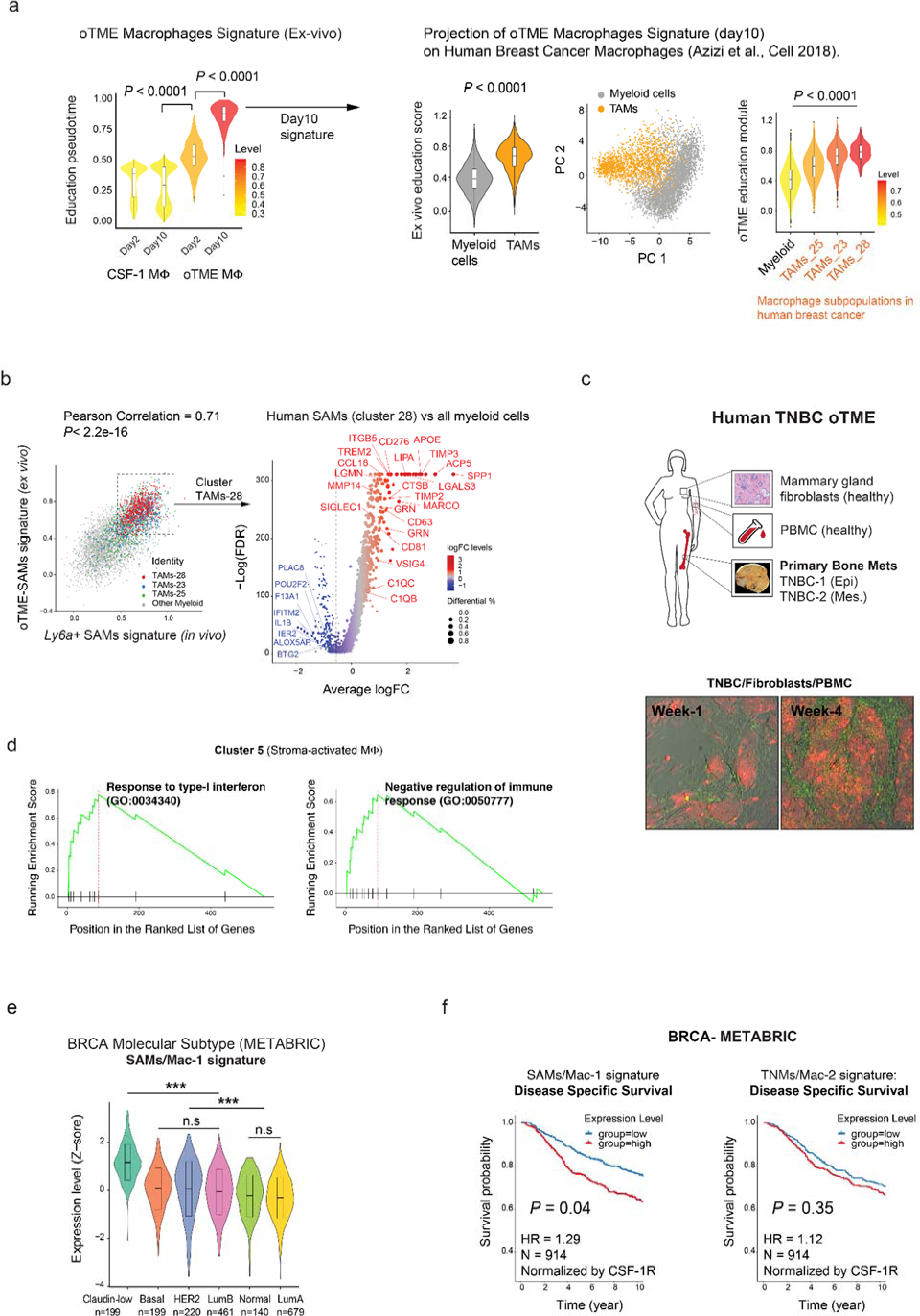
Establishment and association of SAMs and TNMs gene signatures with pathological features in human breast cancer. (**a) Left:** Macrophage oTME education pseudo-time from CSF-1-treated, 2-and-10 days educated macrophages. Wilcoxon Rank Sum test. **Right:** Projection of oTME education-signature (at day 10) on the human “M2-signature” as described by Azizi et al. ^16^ using scRNA-seq data from human breast cancer immune cells. The murine *ex vivo* education-signature score in human myeloid and human tumor-associated macrophages (TAMs; subpopulation (left panel)). Principal component (PC) analysis of myeloid and TAM cells (middle panel). *Ex vivo* M2-signature score in myeloid and three TAMs subpopulations of human breast cancer: TAM_25, TAM_23, TAM_28. The left-to-right ordering represents the least-to-most M2-activation status according to ^16^ (right panel). Wilcoxon Rank Sum test (**b**) Left, gene expression signatures correlation between *Ly6a*+SAMs (*in vivo*), oTME macrophages (*ex vivo*), and breast cancer myeloid subpopulations in human breast cancer. Right, volcano plot showing log fold change (logFC) of differentially expressed genes between SAMs (cluster TAMs-28) and other tumor-infiltrating macrophages and myeloid cells in human breast cancer (**Methods**). Genes are dotted and colored by logFC levels (color scale), the size of each dot represents the difference in the fraction of detection between the two groups. The *P*-values were determined by a Wilcoxon Rank Sum test. Vertical dotted lines represent ±0.5 logFC. Horizontal dotted lines represent a *P*-value of 0.01 (-log10 scale). Right side: key SAM-associated genes. Left side: key TNM-associated genes. (**c**) Generation and *ex vivo* imaging of oTME of human TNBC. TdTomato+ tumor cells are labeled in red and fibroblasts are in green (autofluorescence). (**d**) Gene Set Enrichment Analysis of cluster 5 macrophages shows significant enrichments for type-I Interferon response (GO:0034340) and negative regulation of immune response (GO:0050777). FDR < 0.05. (**e**) Expression levels (Z-score) of SAM/Mac-1 gene signature in different molecular subtypes of breast cancer (METABRIC). Wilcoxon Rank Sum test, \**P* < 0.05, \*\**P* < 0.01, \*\*\**P* < 0.001, otherwise not significant (n.s). (**f**) Kaplan-Meier plots showing disease-specific survival (DSS) in breast cancer patients with higher and lower expression of human CSF1R-normalized SAM and TNM gene signatures in the METABRIC dataset. *P*-value was calculated by multivariate Cox regression adjusting for ER, PR, HER2 status, tumor histologic grade and menopause state (**Methods**). The 10-year survival was shown but the analyses were conducted using all of the data. HR, hazard ratio.

## References

1. Binnewies, M. et al. Understanding the tumor immune microenvironment (TIME) for effective therapy. Nat. Med. 24, 541–550 (2018).

2. de Visser, K. E. & Joyce, J. A. The evolving tumor microenvironment: From cancer initiation to metastatic outgrowth. Cancer Cell 41, 374–403 (2023).

3. Gentles, A. J. et al. The prognostic landscape of genes and infiltrating immune cells across human cancers. Nat. Med. 21, 938–945 (2015).

4. Lin, E. Y. et al. Macrophages regulate the angiogenic switch in a mouse model of breast cancer. Cancer Res. 66, 11238–11246 (2006).

5. Mantovani, A., Allavena, P., Marchesi, F. & Garlanda, C. Macrophages as tools and targets in cancer therapy. Nat. Rev. Drug Discov. (2022) doi:10.1038/s41573-022-00520-5.

6. Qian, B.-Z. et al. CCL2 recruits inflammatory monocytes to facilitate breast-tumour metastasis. Nature 475, 222–225 (2011).

7. Keklikoglou, I. et al. Periostin Limits Tumor Response to VEGFA Inhibition. Cell Rep. 22, 2530–2540 (2018).

8. Hughes, R. et al. Perivascular M2 Macrophages Stimulate Tumor Relapse after Chemotherapy. Cancer Res. 75, 3479–3491 (2015).

9. Krishna, C. et al. Single-cell sequencing links multiregional immune landscapes and tissue-resident T cells in ccRCC to tumor topology and therapy efficacy. Cancer Cell 39, 662–677.e6 (2021).

10. Casanova-Acebes, M. et al. Tissue-resident macrophages provide a pro-tumorigenic niche to early NSCLC cells. Nature 1–7 (2021).

11. DeNardo, D. G. et al. Leukocyte complexity predicts breast cancer survival and functionally regulates response to chemotherapy. Cancer Discov. 1, 54–67 (2011).

12. Campbell, M. J. et al. Proliferating macrophages associated with high grade, hormone receptor negative breast cancer and poor clinical outcome. Breast Cancer Res. Treat. 128, 703–711 (2011).

13. Cassetta, L. & Pollard, J. W. Targeting macrophages: therapeutic approaches in cancer. Nat. Rev. Drug Discov. (2018) doi:10.1038/nrd.2018.169.

14. Cheng, S. et al. A pan-cancer single-cell transcriptional atlas of tumor infiltrating myeloid cells. Cell 184, 792–809.e23 (2021).

15. Sun, L. et al. Activating a collaborative innate-adaptive immune response to control metastasis. Cancer Cell 39, 1361–1374.e9 (2021).

16. Azizi, E. et al. Single-Cell Map of Diverse Immune Phenotypes in the Breast Tumor Microenvironment. Cell 174, 1293–1308.e36 (2018).

17. Chung, W. et al. Single-cell RNA-seq enables comprehensive tumour and immune cell profiling in primary breast cancer. Nat. Commun. 8, 15081 (2017).

18. Obradovic, A. et al. Single-cell protein activity analysis identifies recurrence-associated renal tumor macrophages. Cell 184, 2988–3005.e16 (2021).

19. Wynn, T. A., Chawla, A. & Pollard, J. W. Macrophage biology in development, homeostasis and disease. Nature 496, 445–455 (2013).

20. Kohyama, M. et al. Role for Spi-C in the development of red pulp macrophages and splenic iron homeostasis. Nature 457, 318–321 (2009).

21. Stamatiades, E. G. et al. Immune Monitoring of Trans-endothelial Transport by Kidney-Resident Macrophages. Cell 166, 991–1003 (2016).

22. Wolf, Y. et al. Brown-adipose-tissue macrophages control tissue innervation and homeostatic energy expenditure. Nat. Immunol. 18, 665–674 (2017).

23. Van Nguyen, A. & Pollard, J. W. Colony stimulating factor-1 is required to recruit macrophages into the mammary gland to facilitate mammary ductal outgrowth. Dev. Biol. 247, 11–25 (2002).

24. Chakrabarti, R. et al. Notch ligand Dll1 mediates cross-talk between mammary stem cells and the macrophageal niche. Science 360, (2018).

25. Lavin, Y. et al. Tissue-resident macrophage enhancer landscapes are shaped by the local microenvironment. Cell 159, 1312–1326 (2014).

26. Gosselin, D. et al. Environment drives selection and function of enhancers controlling tissue-specific macrophage identities. Cell 159, 1327–1340 (2014).

27. Gautier, E. L. et al. Gene-expression profiles and transcriptional regulatory pathways that underlie the identity and diversity of mouse tissue macrophages. Nat. Immunol. 13, 1118–1128 (2012).

28. Talmi-Frank, D. et al. Extracellular Matrix Proteolysis by MT1-MMP Contributes to Influenza-Related Tissue Damage and Mortality. Cell Host Microbe 20, 458–470 (2016).

29. Uderhardt, S., Martins, A. J., Tsang, J. S., Lämmermann, T. & Germain, R. N. Resident Macrophages Cloak Tissue Microlesions to Prevent Neutrophil-Driven Inflammatory Damage. Cell 177, 541–555.e17 (2019).

30. Zhou, X. et al. Microglia and macrophages promote corralling, wound compaction and recovery after spinal cord injury via Plexin-B2. Nat. Neurosci. 23, 337–350 (2020).

31. Dvorak, H. F. Tumors: wounds that do not heal. Similarities between tumor stroma generation and wound healing. N. Engl. J. Med. 315, 1650–1659 (1986).

32. Minutti, C. M. et al. Local amplifiers of IL-4Rα–mediated macrophage activation promote repair in lung and liver. Science 356, 1076–1080 (2017).

33. Knipper, J. A. et al. Interleukin-4 Receptor α Signaling in Myeloid Cells Controls Collagen Fibril Assembly in Skin Repair. Immunity 43, 803–816 (2015).

34. Wang, J. & Kubes, P. A Reservoir of Mature Cavity Macrophages that Can Rapidly Invade Visceral Organs to Affect Tissue Repair. Cell 165, 668–678 (2016).

35. Eming, S. A., Wynn, T. A. & Martin, P. Inflammation and metabolism in tissue repair and regeneration. Science 356, 1026–1030 (2017).

36. Ginhoux, F. et al. Fate mapping analysis reveals that adult microglia derive from primitive macrophages. Science 330, 841–845 (2010).

37. Mass, E. et al. Specification of tissue-resident macrophages during organogenesis. Science 353, (2016).

38. Hashimoto, D. et al. Tissue-resident macrophages self-maintain locally throughout adult life with minimal contribution from circulating monocytes. Immunity 38, 792–804 (2013).

39. Schulz, C. et al. A lineage of myeloid cells independent of Myb and hematopoietic stem cells. Science 336, 86–90 (2012).

40. Merad, M. et al. Langerhans cells renew in the skin throughout life under steady-state conditions. Nat. Immunol. 3, 1135–1141 (2002).

41. Yona, S. et al. Fate Mapping Reveals Origins and Dynamics of Monocytes and Tissue Macrophages under Homeostasis. Immunity 38, 79–91 (2013).

42. Wang, Z. et al. An immune cell atlas reveals the dynamics of human macrophage specification during prenatal development. Cell (2023) doi:10.1016/j.cell.2023.08.019.

43. Rantakari, P. et al. Fetal liver endothelium regulates the seeding of tissue-resident macrophages. Nature 538, 392–396 (2016).

44. Jäppinen, N. et al. Fetal-derived macrophages dominate in adult mammary glands. Nat. Commun. 10, 281 (2019).

45. Franklin, R. A. et al. The cellular and molecular origin of tumor-associated macrophages. Science 344, 921–925 (2014).

46. Movahedi, K. et al. Different Tumor Microenvironments Contain Functionally Distinct Subsets of Macrophages Derived from Ly6C(high) Monocytes. Cancer Research vol. 70 5728–5739 Preprint at 10.1158/0008-5472.can-09-4672 (2010).

47. Arwert, E. N. et al. A Unidirectional Transition from Migratory to Perivascular Macrophage Is Required for Tumor Cell Intravasation. Cell Rep. 23, 1239–1248 (2018).

48. Dai, X.-M. et al. Targeted disruption of the mouse colony-stimulating factor 1 receptor gene results in osteopetrosis, mononuclear phagocyte deficiency, increased primitive progenitor cell frequencies, and reproductive defects. Blood 99, 111–120 (2002).

49. Stanley, E. R. & Chitu, V. CSF-1 receptor signaling in myeloid cells. Cold Spring Harb. Perspect. Biol. 6, (2014).

50. Pollard, J. W. & Hennighausen, L. Colony stimulating factor 1 is required for mammary gland development during pregnancy. Proc. Natl. Acad. Sci. U. S. A. 91, 9312–9316 (1994).

51. Wang, Y. et al. IL-34 is a tissue-restricted ligand of CSF1R required for the development of Langerhans cells and microglia. Nat. Immunol. 13, 753–760 (2012).

52. Klappacher, G. W. et al. An induced Ets repressor complex regulates growth arrest during terminal macrophage differentiation. Cell 109, 169–180 (2002).

53. Pixley, F. J. & Stanley, E. R. CSF-1 regulation of the wandering macrophage: complexity in action. Trends Cell Biol. 14, 628–638 (2004).

54. Dawson, C. A. et al. Tissue-resident ductal macrophages survey the mammary epithelium and facilitate tissue remodelling. Nat. Cell Biol. 22, 546–558 (2020).

55. Lin, E. Y., Nguyen, A. V., Russell, R. G. & Pollard, J. W. Colony-stimulating factor 1 promotes progression of mammary tumors to malignancy. J. Exp. Med. 193, 727–740 (2001).

56. Lin, E. Y. et al. Progression to Malignancy in the Polyoma Middle T Oncoprotein Mouse Breast Cancer Model Provides a Reliable Model for Human Diseases. Am. J. Pathol. 163, 2113–2126 (2003).

57. Ben-Chetrit, N. et al. Integration of whole transcriptome spatial profiling with protein markers. Nat. Biotechnol. (2023) doi:10.1038/s41587-022-01536-3.

58. Wu, S. Z. et al. Stromal cell diversity associated with immune evasion in human triple-negative breast cancer. EMBO J. e104063 (2020).

59. Attalla, S., Taifour, T., Bui, T. & Muller, W. Insights from transgenic mouse models of PyMT- induced breast cancer: recapitulating human breast cancer progression in vivo. Oncogene 40, 475– 491 (2021).

60. Sleeman, K. E., Kendrick, H., Ashworth, A., Isacke, C. M. & Smalley, M. J. CD24 staining of mouse mammary gland cells defines luminal epithelial, myoepithelial/basal and non-epithelial cells. Breast Cancer Res. 8, R7 (2006).

61. Drobysheva, D. et al. Transformation of enriched mammary cell populations with polyomavirus middle T antigen influences tumor subtype and metastatic potential. Breast Cancer Res. 17, 132 (2015).

62. Lim, E. et al. Aberrant luminal progenitors as the candidate target population for basal tumor development in BRCA1 mutation carriers. Nat. Med. 15, 907–913 (2009).

63. Nolan, E. et al. RANK ligand as a potential target for breast cancer prevention in BRCA1-mutation carriers. Nat. Med. 22, 933–939 (2016).

64. Lim, E. et al. Transcriptome analyses of mouse and human mammary cell subpopulations reveal multiple conserved genes and pathways. Breast Cancer Res. 12, R21 (2010).

65. Eirew, P. et al. A method for quantifying normal human mammary epithelial stem cells with in vivo regenerative ability. Nat. Med. 14, 1384–1389 (2008).

66. Friedman, G. et al. Cancer-associated fibroblast compositions change with breast cancer progression linking the ratio of S100A4 and PDPN CAFs to clinical outcome. Nature Cancer vol. 1 692–708 Preprint at 10.1038/s43018-020-0082-y (2020).

67. Bartoschek, M. et al. Spatially and functionally distinct subclasses of breast cancer-associated fibroblasts revealed by single cell RNA sequencing. Nat. Commun. 9, 5150 (2018).

68. Muzumdar, M. D., Tasic, B., Miyamichi, K., Li, L. & Luo, L. A global double-fluorescent Cre reporter mouse. Genesis 45, 593–605 (2007).

69. Gabrilovich, D. I. Myeloid-Derived Suppressor Cells. Cancer Immunol Res 5, 3–8 (2017).

70. Keren, L. et al. A Structured Tumor-Immune Microenvironment in Triple Negative Breast Cancer Revealed by Multiplexed Ion Beam Imaging. Cell 174, 1373–1387.e19 (2018).

71. Kratochvill, F. et al. TNF Counterbalances the Emergence of M2 Tumor Macrophages. Cell Rep. 12, 1902–1914 (2015).

72. Biswas, S. K. & Mantovani, A. Macrophage plasticity and interaction with lymphocyte subsets: cancer as a paradigm. Nat. Immunol. 11, 889–896 (2010).

73. Colegio, O. R. et al. Functional polarization of tumour-associated macrophages by tumour-derived lactic acid. Nature 513, 559–563 (2014).

74. Zhou, X. et al. Circuit Design Features of a Stable Two-Cell System. Cell 172, 744–757.e17 (2018).

75. Aziz, A., Soucie, E., Sarrazin, S. & Sieweke, M. H. MafB/c-Maf deficiency enables self-renewal of differentiated functional macrophages. Science 326, 867–871 (2009).

76. Quail, D. F. et al. Obesity alters the lung myeloid cell landscape to enhance breast cancer metastasis through IL5 and GM-CSF. Nat. Cell Biol. 19, 974–987 (2017).

77. Honda, K. et al. IRF-7 is the master regulator of type-I interferon-dependent immune responses. Nature 434, 772–777 (2005).

78. Honda, K. & Taniguchi, T. IRFs: master regulators of signalling by Toll-like receptors and cytosolic pattern-recognition receptors. Nat. Rev. Immunol. 6, 644–658 (2006).

79. Motwani, M., Pesiridis, S. & Fitzgerald, K. A. DNA sensing by the cGAS-STING pathway in health and disease. Nat. Rev. Genet. 20, 657–674 (2019).

80. Zhou, Y. et al. Blockade of the Phagocytic Receptor MerTK on Tumor-Associated Macrophages Enhances P2X7R-Dependent STING Activation by Tumor-Derived cGAMP. Immunity 52, 357– 373.e9 (2020).

81. Ito, C. Y., Li, C. Y. J., Bernstein, A., Dick, J. E. & Stanford, W. L. Hematopoietic stem cell and progenitor defects in Sca-1/Ly-6A–null mice. Blood 101, 517–523 (2003).

82. Essers, M. A. G. et al. IFNalpha activates dormant haematopoietic stem cells in vivo. Nature 458, 904–908 (2009).

83. Walter, D. et al. Exit from dormancy provokes DNA-damage-induced attrition in haematopoietic stem cells. Nature 520, 549–552 (2015).

84. Burke, J. R. et al. Autoimmune pathways in mice and humans are blocked by pharmacological stabilization of the TYK2 pseudokinase domain. Sci. Transl. Med. 11, (2019).

85. Prchal-Murphy, M. et al. TYK2 kinase activity is required for functional type I interferon responses in vivo. PLoS One 7, e39141 (2012).

86. Bohrer, L. R. et al. ADAM17 in tumor associated leukocytes regulates inflammatory mediators and promotes mammary tumor formation. Genes Cancer 7, 240–253 (2016).

87. Wong, E. et al. Harnessing the natural inhibitory domain to control TNFα Converting Enzyme (TACE) activity in vivo. Sci. Rep. 6, 35598 (2016).

88. van Es, J. H. et al. Notch/gamma-secretase inhibition turns proliferative cells in intestinal crypts and adenomas into goblet cells. Nature 435, 959–963 (2005).

89. Imbimbo, B. P. Therapeutic potential of gamma-secretase inhibitors and modulators. Curr. Top. Med. Chem. 8, 54–61 (2008).

90. Paffenholz, S. V. et al. Senescence induction dictates response to chemo- and immunotherapy in preclinical models of ovarian cancer. Proc. Natl. Acad. Sci. U. S. A. 119, (2022).

91. Baghdadi, M. et al. TIM-4 glycoprotein-mediated degradation of dying tumor cells by autophagy leads to reduced antigen presentation and increased immune tolerance. Immunity 39, 1070–1081 (2013).

92. Alshetaiwi, H. et al. Defining the emergence of myeloid-derived suppressor cells in breast cancer using single-cell transcriptomics. Sci Immunol 5, (2020).

93. Veglia, F., Perego, M. & Gabrilovich, D. Myeloid-derived suppressor cells coming of age. Nat. Immunol. 19, 108–119 (2018).

94. Miller, B. F., Bambah-Mukku, D., Dulac, C., Zhuang, X. & Fan, J. Characterizing spatial gene expression heterogeneity in spatially resolved single-cell transcriptomics data with nonuniform cellular densities. Genome Res. (2021) doi:10.1101/gr.271288.120.

95. Hao, Y. et al. Integrated analysis of multimodal single-cell data. Cell 0, (2021).

96. Eschweiler, S. et al. JAML immunotherapy targets recently activated tumor-infiltrating CD8+ T cells. Cell Rep. 42, 112040 (2023).

97. Segovia, M. et al. Targeting TMEM176B Enhances Antitumor Immunity and Augments the Efficacy of Immune Checkpoint Blockers by Unleashing Inflammasome Activation. Cancer Cell 35, 767–781.e6 (2019).

98. 98. Yeboah, M., et al. LILRB3 (ILT5) is a myeloid cell checkpoint that elicits profound immunomodulation. JCI Insight 5, (2020).

99. Cassetta, L. et al. Human Tumor-Associated Macrophage and Monocyte Transcriptional Landscapes Reveal Cancer-Specific Reprogramming, Biomarkers, and Therapeutic Targets. Cancer Cell 35, 588–602.e10 (2019).

100. Curtis, C. et al. The genomic and transcriptomic architecture of 2,000 breast tumours reveals novel subgroups. Nature 486, 346–352 (2012).

101. DeNardo, D. G. & Ruffell, B. Macrophages as regulators of tumour immunity and immunotherapy. Nature Reviews Immunology vol. 19 369–382 Preprint at 10.1038/s41577-019-0127-6 (2019).

102. Kaneda, M. M. et al. PI3Kγ is a molecular switch that controls immune suppression. Nature 539, 437–442 (2016).

103. Tseng, D. et al. Anti-CD47 antibody-mediated phagocytosis of cancer by macrophages primes an effective antitumor T-cell response. Proc. Natl. Acad. Sci. U. S. A. 110, 11103–11108 (2013).

104. Pyonteck, S. M. et al. CSF-1R inhibition alters macrophage polarization and blocks glioma progression. Nat. Med. 19, 1264–1272 (2013).

105. Barkal, A. A. et al. CD24 signalling through macrophage Siglec-10 is a target for cancer immunotherapy. Nature 572, 392–396 (2019).

106. Ries, C. H. et al. Targeting tumor-associated macrophages with anti-CSF-1R antibody reveals a strategy for cancer therapy. Cancer Cell 25, 846–859 (2014).

107. Klichinsky, M. et al. Human chimeric antigen receptor macrophages for cancer immunotherapy. Nat. Biotechnol. 38, 947–953 (2020).

108. Rannikko, J. H., et al. Bexmarilimab-induced macrophage activation leads to treatment benefit in solid tumors: the phase I/II first-in-human MATINS trial. bioRxiv (2023) doi:10.1101/2023.04.17.23288693.

109. Angelo, M. et al. Multiplexed ion beam imaging of human breast tumors. Nat. Med. 20, 436–442 (2014).

110. Robbins, C. S. et al. Local proliferation dominates lesional macrophage accumulation in atherosclerosis. Nat. Med. 19, 1166–1172 (2013).

111. Bosurgi, L. et al. Macrophage function in tissue repair and remodeling requires IL-4 or IL-13 with apoptotic cells. Science 356, 1072–1076 (2017).

112. Amano, S. U. et al. Local proliferation of macrophages contributes to obesity-associated adipose tissue inflammation. Cell Metab. 19, 162–171 (2014).

113. Hamid, M. et al. A Phase 1 study Combining Pexidartinib, Radiation Therapy, and Androgen Deprivation Therapy in Men With Intermediate- and High-Risk Prostate Cancer. Adv Radiat Oncol 6, 100679 (2021).

114. Bendell, J. C. et al. A phase 1 study of ARRY-382, an oral inhibitor of colony-stimulating factor-1 receptor (CSF1R), in patients with advanced or metastatic cancers. in Molecular Cancer Therapeutics vol. 12 (AMER ASSOC CANCER RESEARCH 615 CHESTNUT ST, 17TH FLOOR, PHILADELPHIA, PA…, 2013).

115. Cannarile, M. A. et al. Colony-stimulating factor 1 receptor (CSF1R) inhibitors in cancer therapy. J Immunother Cancer 5, 53 (2017).

116. Meurette, O. & Mehlen, P. Notch Signaling in the Tumor Microenvironment. Cancer Cell 34, 536– 548 (2018).

117. Ohnuki, H. et al. Tumor-infiltrating myeloid cells activate Dll4/Notch/TGF-β signaling to drive malignant progression. Cancer Res. 74, 2038–2049 (2014).

118. Zhu, Y. et al. CSF1/CSF1R blockade reprograms tumor-infiltrating macrophages and improves response to T-cell checkpoint immunotherapy in pancreatic cancer models. Cancer Res. 74, 5057– 5069 (2014).

119. Kim, I. S. et al. Immuno-subtyping of breast cancer reveals distinct myeloid cell profiles and immunotherapy resistance mechanisms. Nat. Cell Biol. 21, 1113–1126 (2019).

120. McGinnis, C. S., Murrow, L. M. & Gartner, Z. J. DoubletFinder: Doublet detection in single-cell RNA sequencing data using artificial nearest neighbors. bioRxiv 352484 (2018) doi:10.1101/352484.

121. Hafemeister, C. & Satija, R. Normalization and variance stabilization of single-cell RNA-seq data using regularized negative binomial regression. Genome Biol. 20, 296 (2019).

122. Gu, Z., Eils, R. & Schlesner, M. Complex heatmaps reveal patterns and correlations in multidimensional genomic data. Bioinformatics 32, 2847–2849 (2016).

123. Ben-Chetrit, N. et al. Synaptojanin 2 is a druggable mediator of metastasis and the gene is overexpressed and amplified in breast cancer. Sci. Signal. 8, ra7 (2015).

124. Ben-Chetrit, N. et al. Integrated protein and transcriptome high-throughput spatial profiling. bioRxiv 2022.03.15.484516 (2022) doi:10.1101/2022.03.15.484516.

125. Dobin, A. et al. STAR: ultrafast universal RNA-seq aligner. Bioinformatics 29, 15–21 (2013).

126. Love, M. I., Huber, W. & Anders, S. Moderated estimation of fold change and dispersion for RNA- seq data with DESeq2. Genome Biol. 15, 550 (2014).

127. Yu, G., Wang, L.-G., Han, Y. & He, Q.-Y. clusterProfiler: an R package for comparing biological themes among gene clusters. OMICS 16, 284–287 (2012).

128. Butler, A., Hoffman, P., Smibert, P., Papalexi, E. & Satija, R. Integrating single-cell transcriptomic data across different conditions, technologies, and species. Nat. Biotechnol. 36, 411–420 (2018).

129. Maaten, L. van der & Hinton, G. Visualizing Data using t-SNE. J. Mach. Learn. Res. 9, 2579–2605 (2008).

130. DePasquale, E. A. K. et al. DoubletDecon: Deconvoluting Doublets from Single-Cell RNA-Sequencing Data. Cell Rep. 29, 1718–1727.e8 (2019).

131. Coifman, R. R. & Lafon, S. Diffusion maps. Appl. Comput. Harmon. Anal. 21, 5–30 (2006).

132. Finak, G. et al. MAST: a flexible statistical framework for assessing transcriptional changes and characterizing heterogeneity in single-cell RNA sequencing data. Genome Biol. 16, 278 (2015).

133. Angerer, P. et al. destiny: diffusion maps for large-scale single-cell data in R. Bioinformatics 32, 1241–1243 (2016).

134. Hastie, T. & Stuetzle, W. Principal Curves. J. Am. Stat. Assoc. 84, 502–516 (1989).

135. Wang, W. et al. A single-cell transcriptional roadmap for cardiopharyngeal fate diversification. Nat. Cell Biol. 21, 674–686 (2019).

136. Meyer, P. E. Information-theoretic variable selection and network inference from microarray data. Universite Libre de Bruxelles. [Google Scholar] (2008).

137. Tirosh, I. et al. Dissecting the multicellular ecosystem of metastatic melanoma by single-cell RNA- seq. Science 352, 189–196 (2016).

138. Roelli, P., bbimber, Flynn, B., santiagorevale & Gui, G. Hoohm/CITE-seq-Count: 1.4.2. (2019). doi:10.5281/zenodo.2590196.

139. Ord, J. K. & Getis, A. Local spatial autocorrelation statistics: Distributional issues and an application. Geogr. Anal. 27, 286–306 (2010).

140. Anselin, L. Local indicators of spatial association-LISA. Geogr. Anal. 27, 93–115 (2010).

141. Getis, A. & Ord, J. K. The analysis of spatial association by use of distance statistics. Geogr. Anal. 24, 189–206 (2010).

142. Middleton, L. & Sivaswamy, J. Edge detection in a hexagonal-image processing framework. Image Vis. Comput. 19, 1071–1081 (2001).

143. Elosua-Bayes, M., Nieto, P., Mereu, E., Gut, I. & Heyn, H. SPOTlight: seeded NMF regression to deconvolute spatial transcriptomics spots with single-cell transcriptomes. Nucleic Acids Res. (2021) doi:10.1093/nar/gkab043.

144. Valdés-Mora, F. et al. Single-cell transcriptomics reveals involution mimicry during the specification of the basal breast cancer subtype. Cell Rep. 35, 108945 (2021).

145. Haghverdi, L., Lun, A. T. L., Morgan, M. D. & Marioni, J. C. Batch effects in single-cell RNA- sequencing data are corrected by matching mutual nearest neighbors. Nat. Biotechnol. (2018) doi:10.1038/nbt.4091.

146. Becht, E. et al. Dimensionality reduction for visualizing single-cell data using UMAP. Nat. Biotechnol. (2018) doi:10.1038/nbt.4314.

147. Efron, B. Bootstrap Methods: Another Look at the Jackknife. aos 7, 1–26 (1979).

148. Chung, N. C. & Storey, J. D. Statistical significance of variables driving systematic variation in high-dimensional data. Bioinformatics 31, 545–554 (2015).

149. van Dijk, D. et al. Recovering Gene Interactions from Single-Cell Data Using Data Diffusion. Cell (2018) doi:10.1016/j.cell.2018.05.061.

150. Moran, P. A. P. Notes on continuous stochastic phenomena. Biometrika 37, 17–23 (1950).

151. Wartenberg, D. Multivariate spatial correlation: A method for exploratory geographical analysis. Geogr. Anal. 17, 263–283 (1985).

152. Czaplewski, R. L. Expected Value and Variance of Moran’s Bivariate Spatial Autocorrelation Statistic for a Permutation Test. (U.S. Department of Agriculture, Forest Service, Rocky Mountain Forest and Range Experiment Station, 1993).

153. Lee, S.-I. Developing a bivariate spatial association measure: An integration of Pearson’s r and Moran’s I. J. Geogr. Syst. 3, 369–385 (2001).

154. Korotkevich, G., Sukhov, V. & Sergushichev, A. Fast gene set enrichment analysis. bioRxiv 060012 (2019) doi:10.1101/060012.

