## Supplementary Note 1 for "Breast Cancer Macrophage Heterogeneity and Self-renewal are Determined by Spatial Localization"

### Supplementary Note MMTV-PyMT Macrophage scRNA-seq Analysis

Steve X. Niu*

Last Run: April 13, 2021

---

1. Load Data This markdown file contains the MMTV-PyMT scRNA-seq analysis for the paper titled **Organotypic Tumor Microenvironment Modeling of Macrophage Education and Proliferation in Breast Cancer**. The data used for this analysis can be found under the GEO SuperSeries GSE171463.

```
# install Seurat v3.0.0
if (!requireNamespace("Seurat", quietly = TRUE) | utils::packageVersion("Seurat") !=
    "3.0.0") remotes::install_version("Seurat", version = "3.0.0")

# if doesn't work install.packages('remotes')
# remotes::install_version('SDMTools', '1.1-221')

# load data
library(Seurat)
mouse <- Read10X("in_vivo_data/")
mouse <- CreateSeuratObject(mouse)

# Remove low quality cells that have more than 10%
# mitocondrial genes.
mouse[["hg.percent.mt"]] <- PercentageFeatureSet(mouse, pattern = "hg19-MT-")
mouse[["mm.percent.mt"]] <- PercentageFeatureSet(mouse, pattern = "mm10-mt-")
mouse <- SubsetData(mouse, subset.name = "hg.percent.mt", high.threshold = 10)
mouse <- SubsetData(mouse, subset.name = "mm.percent.mt", high.threshold = 10)

mouse <- NormalizeData(mouse, verbose = FALSE)
mouse <- FindVariableFeatures(mouse, selection.method = "vst",
    nfeatures = 2000, verbose = FALSE)
mouse <- ScaleData(mouse, verbose = FALSE)
mouse <- RunPCA(mouse, verbose = FALSE)
mouse <- RunTSNE(mouse, dims = 1:30)
mouse <- FindNeighbors(mouse, dims = 1:30, verbose = FALSE)
mouse <- FindClusters(mouse, resolution = 0.005, verbose = FALSE)

# Retain only the mouse cells
mouse <- subset(mouse, ident = 0)
```

*Tri-Institutional Training Program in Computational Biology and Medicine,

```r
mouse.data <- mouse@assays$RNA@counts[grep("mm10-", rownames(mouse)),
    ]
rownames(mouse.data) <- gsub("mm10-", "", rownames(mouse.data))
mouse <- CreateSeuratObject(mouse.data)
```

2. Dimensionality Reduction and Clustering

```r
mouse <- NormalizeData(mouse, verbose = FALSE)
mouse <- FindVariableFeatures(mouse, selection.method = "vst",
    nfeatures = 2000, verbose = FALSE)
mouse <- ScaleData(mouse, verbose = FALSE)
mouse <- RunPCA(mouse, verbose = FALSE)
mouse <- RunTSNE(mouse, dims = 1:20)
mouse <- FindNeighbors(mouse, dims = 1:20, verbose = FALSE)
mouse <- FindClusters(mouse, resolution = 0.2, verbose = FALSE)
mouse <- RenameIdents(mouse, `0` = "Epithelia1", `1` = "Mac1",
    `2` = "Epithelia2", `3` = "TNK", `4` = "MDSC", `5` = "Mac2",
    `6` = "RBC")
mouse@active.ident <- factor(mouse@active.ident, levels = sort(levels(mouse@active.ident)),
    ordered = T)

# Find cell type markers
markers.use <- FindAllMarkers(mouse, logfc.threshold = 1, min.pct = 0.5,
    only.pos = TRUE, verbose = FALSE)
markers.use <- unlist(lapply(split(markers.use, markers.use$cluster),
    function(x) head(x$gene, 5)))
mouse <- ScaleData(mouse, markers.use, verbose = FALSE)
DoHeatmap(mouse, features = markers.use, label = FALSE, raster = FALSE)
```

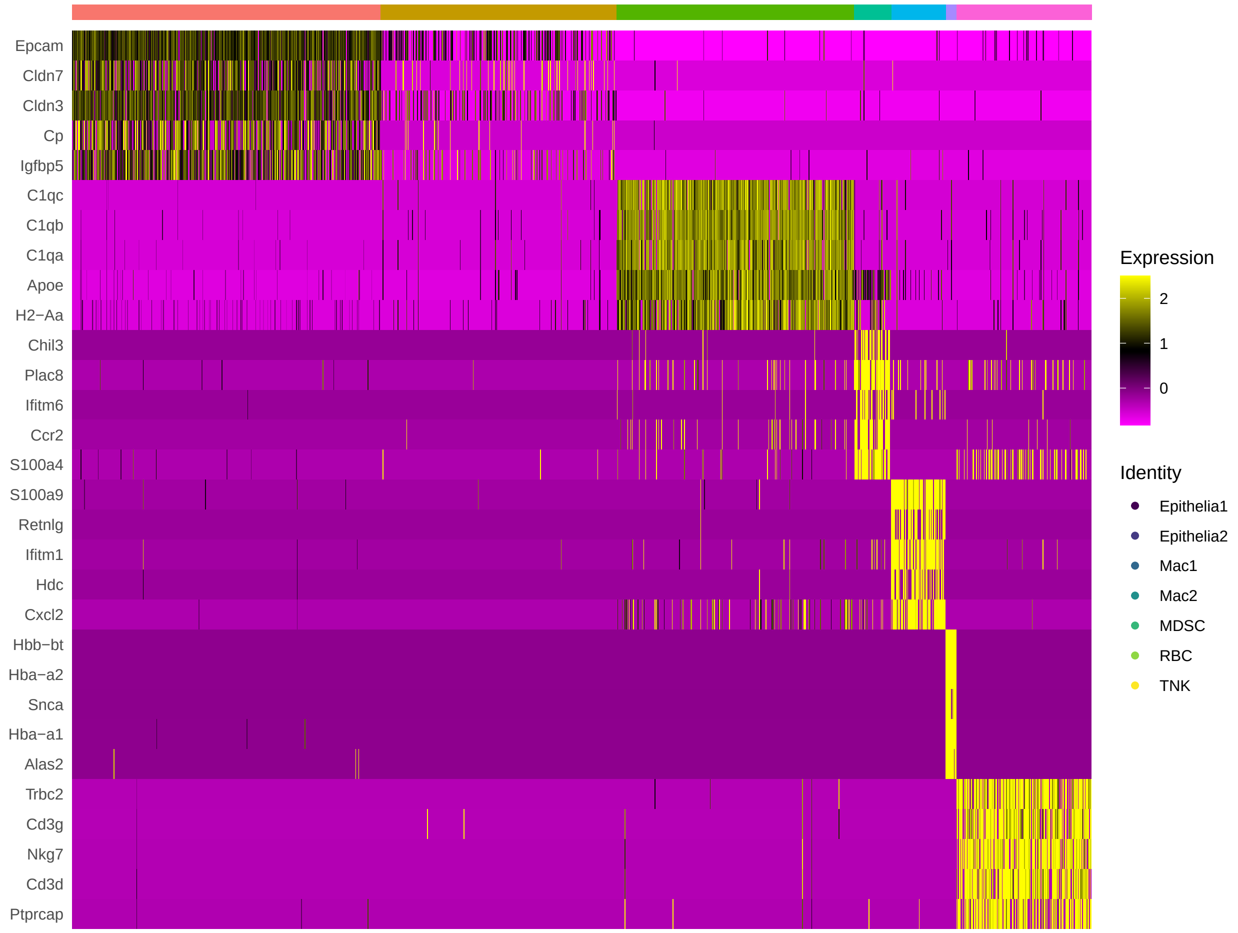


3. Immune cells and macrophage populations

```
mouse <- subset(mouse, ident = c("TNK", "Mac1", "Mac2", "MDSC"))
mouse <- FindVariableFeatures(mouse, selection.method = "vst",
    nfeatures = 2000, verbose = FALSE)
mouse <- ScaleData(mouse, verbose = FALSE)
mouse <- RunPCA(mouse, verbose = FALSE)
mouse <- RunTSNE(mouse, dims = 1:20)
DimPlot(mouse, reduction = "tsne")
```

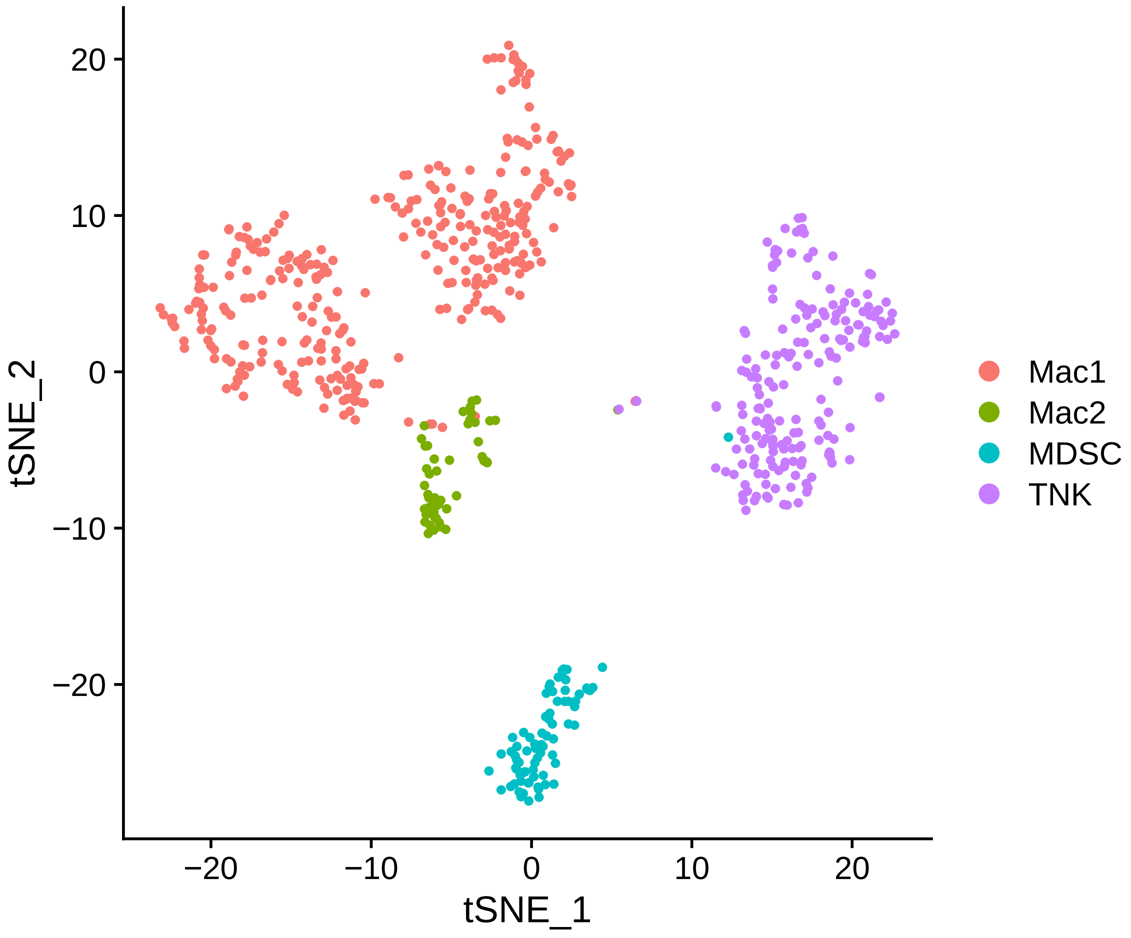


```
VlnPlot(mouse, c("Lyz2", "Cd14", "Cd68", "Aif1", "H2-Aa", "Ly6a"),
    idents = c("MDSC", "Mac1", "Mac2"))
```

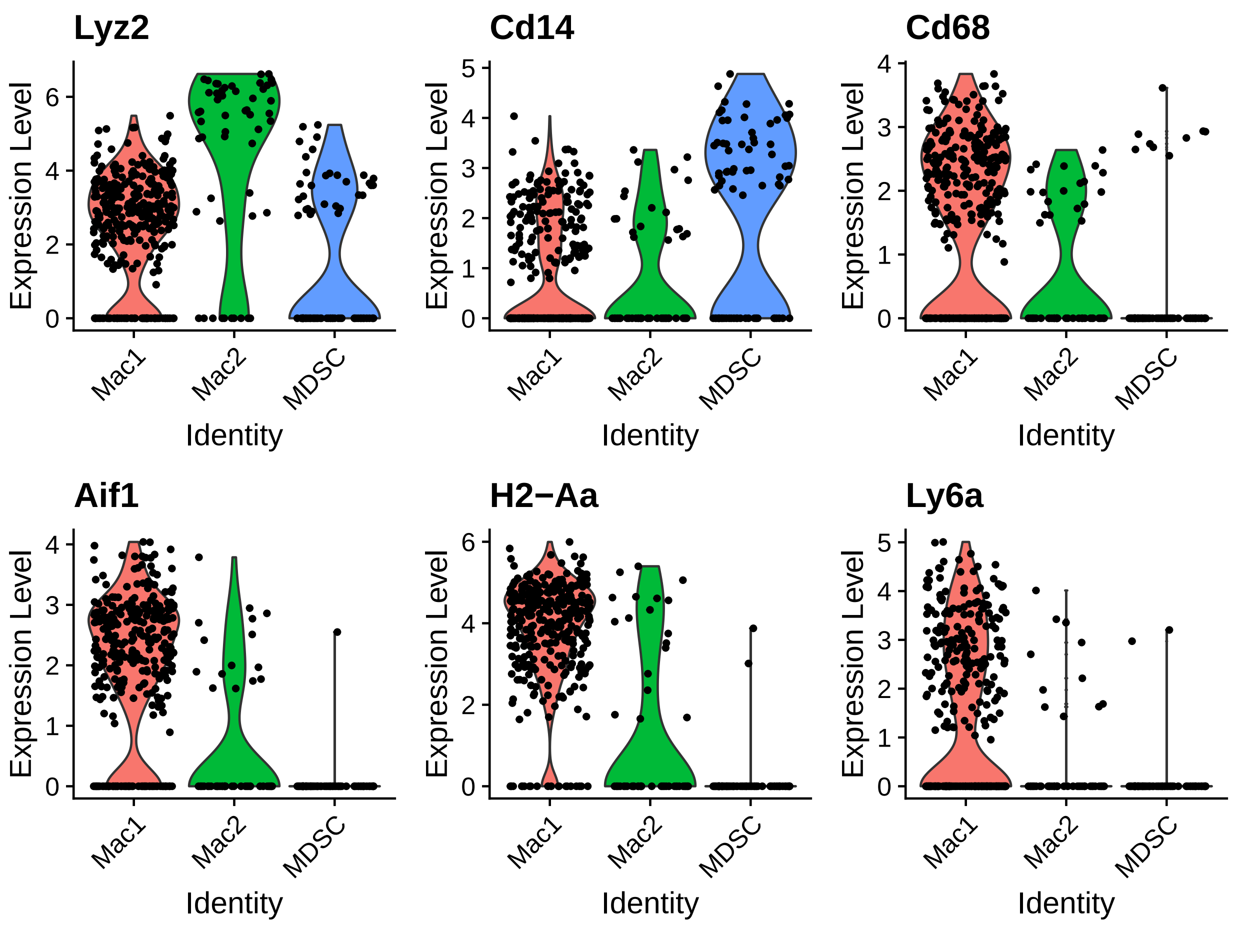


```
mac.markers <- FindMarkers(mouse, "Mac1", "Mac2", logfc.threshold = 0.4,
    min.pct = 0.75, verbose = FALSE)
mac.markers <- mac.markers[mac.markers$p_val_adj < 0.05, ]
mac.markers <- mac.markers[order(mac.markers$avg_logFC, decreasing = T),
    ]
macrophage <- subset(mouse, ident = c("Mac1", "Mac2"))
macrophage <- ScaleData(macrophage, rownames(mac.markers), verbose = FALSE)
DoHeatmap(macrophage, rownames(mac.markers), label = FALSE, disp.max = 1.5,
    disp.min = -1.5, raster = FALSE)
```

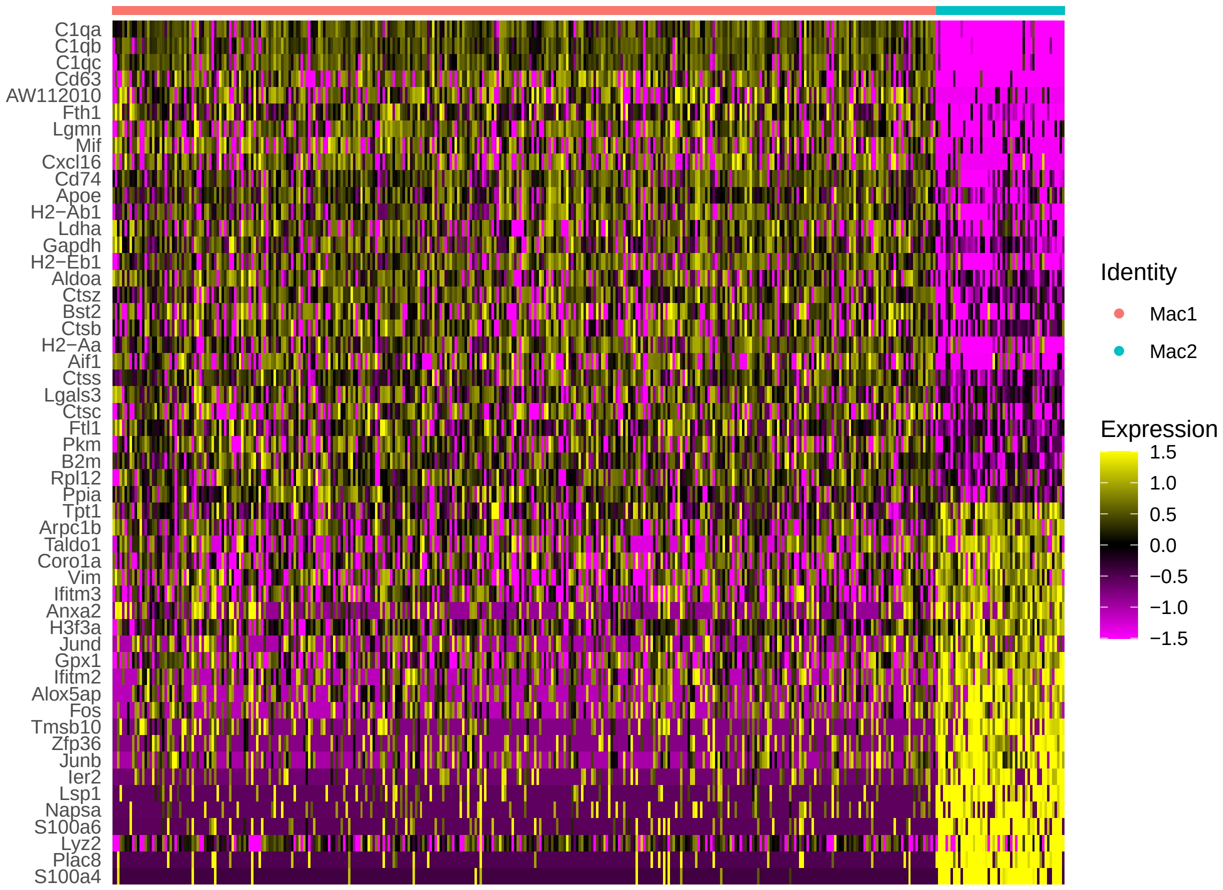


```r
# Session info
print(sessionInfo())
```

```
## R version 3.6.0 (2019-04-26)
## Platform: x86_64-pc-linux-gnu (64-bit)
## Running under: CentOS Linux 7 (Core)
##
## Matrix products: default
## BLAS:   /nfs/sw/R/R-3.6.0/lib64/R/lib/libRblas.so
## LAPACK: /nfs/sw/R/R-3.6.0/lib64/R/lib/libRlapack.so
##
## locale:
##  [1] LC_CTYPE=en_US.UTF-8       LC_NUMERIC=C
##  [3] LC_TIME=en_US.UTF-8        LC_COLLATE=en_US.UTF-8
##  [5] LC_MONETARY=en_US.UTF-8    LC_MESSAGES=en_US.UTF-8
##  [7] LC_PAPER=en_US.UTF-8       LC_NAME=C
##  [9] LC_ADDRESS=C               LC_TELEPHONE=C
## [11] LC_MEASUREMENT=en_US.UTF-8 LC_IDENTIFICATION=C
##
## attached base packages:
## [1] stats     graphics  grDevices utils     datasets  methods   base
##
## other attached packages:
```

```
## [1] Seurat_3.0.0
##
## loaded via a namespace (and not attached):
##   [1] tsne_0.1-3          nlme_3.1-145        RColorBrewer_1.1-2
##   [4] httr_1.4.2          numDeriv_2016.8-1.1 sctransform_0.3
##   [7] tools_3.6.0         R6_2.4.1            irlba_2.3.3
##  [10] KernSmooth_2.23-16  lazyeval_0.2.2      BiocGenerics_0.32.0
##  [13] colorspace_1.4-1    sn_1.5-5            withr_2.2.0
##  [16] gridExtra_2.3       tidyselect_1.1.0    mnormt_1.5-7
##  [19] compiler_3.6.0      Biobase_2.46.0      formatR_1.7
##  [22] TFisher_0.2.0       plotly_4.9.2.1      sandwich_2.5-1
##  [25] labeling_0.3        scales_1.1.1        lmtest_0.9-38
##  [28] mvtnorm_1.1-0       ggridges_0.5.2      pbapply_1.4-2
##  [31] stringr_1.4.0       digest_0.6.25       rmarkdown_2.7
##  [34] R.utils_2.10.1      pkgconfig_2.0.3     htmltools_0.5.0
##  [37] bibtex_0.4.2.2      plotrix_3.7-7       htmlwidgets_1.5.1
##  [40] rlang_0.4.6         farver_2.0.3        zoo_1.8-8
##  [43] jsonlite_1.6.1      ica_1.0-2           dplyr_0.8.5
##  [46] R.oo_1.24.0         magrittr_1.5        Matrix_1.2-18
##  [49] Rcpp_1.0.4.6        munsell_0.5.0       reticulate_1.15
##  [52] ape_5.3             lifecycle_0.2.0     R.methodsS3_1.8.1
##  [55] stringi_1.4.6       multcomp_1.4-12     yaml_2.2.1
##  [58] gbRd_0.4-11         MASS_7.3-51.5       Rtsne_0.15
##  [61] plyr_1.8.6          grid_3.6.0          parallel_3.6.0
##  [64] listenv_0.8.0       ggrepel_0.8.2       crayon_1.3.4
##  [67] lattice_0.20-40     cowplot_1.1.0       splines_3.6.0
##  [70] multtest_2.42.0     SDMTools_1.1-221    knitr_1.28
##  [73] pillar_1.4.4        igraph_1.2.5        reshape2_1.4.4
##  [76] future.apply_1.6.0  codetools_0.2-16    stats4_3.6.0
##  [79] mutoss_0.1-12       glue_1.4.1          evaluate_0.14
##  [82] metap_1.3           data.table_1.13.0   png_0.1-7
##  [85] vctrs_0.3.0         Rdpack_0.11-1       tidyr_1.1.0
##  [88] gtable_0.3.0        RANN_2.6.1          purrr_0.3.4
##  [91] future_1.19.1       assertthat_0.2.1    ggplot2_3.3.2
##  [94] xfun_0.22           rsvd_1.0.3          viridisLite_0.3.0
##  [97] survival_3.1-11     tibble_3.0.1        cluster_2.1.0
## [100] globals_0.13.0      fitdistrplus_1.1-1  TH.data_1.0-10
## [103] ellipsis_0.3.1      ROCR_1.0-11
```
