## Supplementary Note 2 for "Breast Cancer Macrophage Heterogeneity and Self-renewal are Determined by Spatial Localization"

### Supplementary Note oTME Macrophage scRNA-seq Analysis

Steve X. Niu*

Last Run: April 13, 2021

---

#### 1. Load Data

This markdown file contains the oTME scRNA-seq analysis for the paper titled **Organotypic Tumor Microenvironment Modeling of Macrophage Education and Proliferation in Breast Cancer**. The data used for this analysis can be found under the GEO SuperSeries GSE171463.

```
# install pcakages from CRAN if not installed
cran.packages <- setdiff(c("utils", "remote", "data.table", "princurve",
    "infotheo", "scatterpie", "ggrepel"), installed.packages())
if (length(cran.packages) > 0) install.packages(cran.packages)

# install pcakages from Bioconductor if not installed
bioc.packages <- setdiff(c("scater", "rhdf5", "destiny", "MAST"),
    installed.packages())
if (length(bioc.packages) > 0) {
    if (!requireNamespace("BiocManager", quietly = TRUE))
        install.packages("BiocManager")
    BiocManager::install(bioc.packages)
}

# install Seurat v2.3.4
if (!requireNamespace("Seurat", quietly = TRUE) | utils::packageVersion("Seurat") !=
    "2.3.4") remotes::install_version("Seurat", version = "2.3.4")

# load data
library(Seurat)
library(destiny)
library(princurve)
library(infotheo)
library(ggrepel)
source("functions.R")
data_all <- load.data("TME_Paper_Data/")

# Remove cells that have less than 500 genes and genes that
# are not expressed by at least 50 cells.
tme <- CreateSeuratObject(data_all, min.cells = 50, min.genes = 500,
    names.field = 1:4)
```

*Tri-Institutional Training Program in Computational Biology and Medicine,

```r
tme <- NormalizeData(tme, display.progress = FALSE)

# Get mitochondrial gene expression percentage.
mito.genes <- grep(pattern = "^mt", x = rownames(x = tme@data),
    value = TRUE)
percent.mito <- Matrix::colSums(tme@raw.data[mito.genes, ])/Matrix::colSums(tme@raw.data)
tme <- AddMetaData(object = tme, metadata = percent.mito, col.name = "percent.mito")
condition <- unlist(lapply(as.character(tme@ident), ExtractField,
    field = 1:2))
tme <- AddMetaData(object = tme, metadata = data.frame(condition,
    row.names = tme@cell.names), col.name = "condition")
cols.use <- gg.color(20)
VlnPlot(object = tme, features.plot = c("nGene", "percent.mito"),
    nCol = 2, group.by = "orig.ident", x.lab.rot = TRUE, point.size.use = 0.1,
    cols.use = cols.use)
```

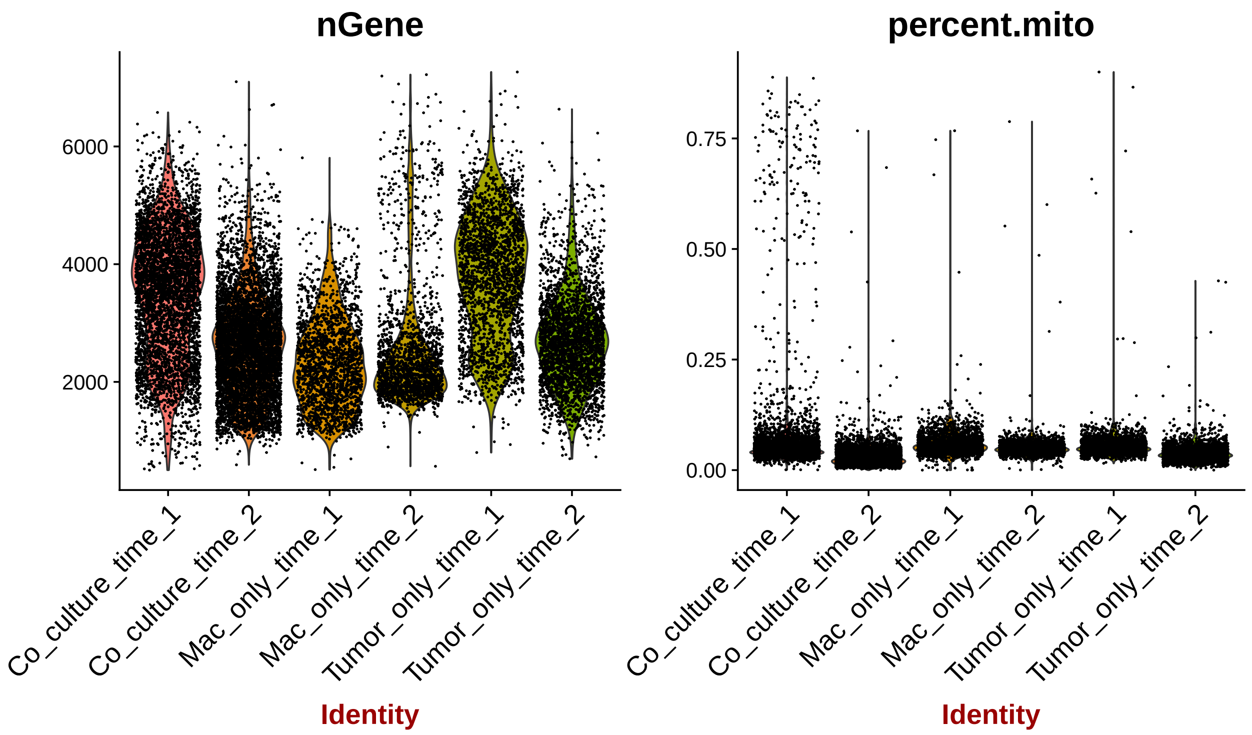


##### 2. Dimensionality Reduction and Clustering

```r
# Filter out cells that have more than 10% of mitochondrial
# gene counts.
tme <- SubsetData(tme, subset.name = "percent.mito", accept.high = 0.1)
tme <- ScaleData(tme, display.progress = FALSE)
tme <- FindVariableGenes(tme, do.plot = FALSE, display.progress = FALSE)
tme <- RunPCA(tme, do.print = FALSE)

# optional PC elbow plot PCElbowPlot(tme)
```

```r
tme <- RunTSNE(tme, dims.use = 1:15)
tme <- FindClusters(tme, save.SNN = T, dims.use = 1:15, reduction.type = "pca",
    resolution = 1, print.output = FALSE)
tme <- BuildClusterTree(tme, do.reorder = TRUE, reorder.numeric = TRUE,
    pcs.use = 1:15, show.progress = FALSE, do.plot = TRUE)
```

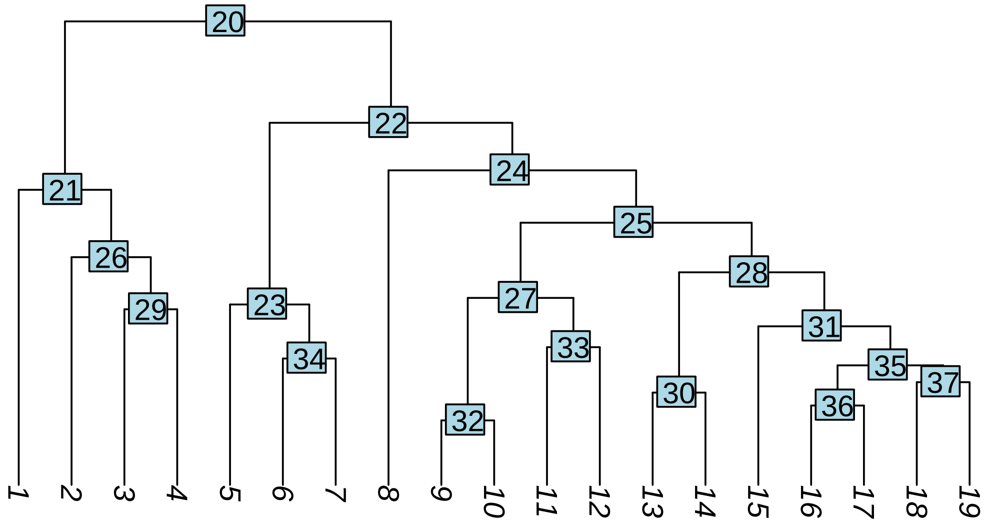


```r
# Assess 25% bottom nodes out of bag error,
oobe <- AssessNodes(tme, node.list = c(32, 30, 35, 36, 37))

## Growing trees.. Progress: 94%. Estimated remaining time: 1 seconds.
```

```r
# Merge nodes using 0.03 as criteria
for (i in oobe$node[which(oobe$oobe > 0.03)]) {
    tme <- MergeNode(tme, node.use = i, rebuild.tree = FALSE)
}
tme <- BuildClusterTree(tme, do.reorder = TRUE, reorder.numeric = TRUE,
    pcs.use = 1:15, do.plot = TRUE, show.progress = FALSE)
```

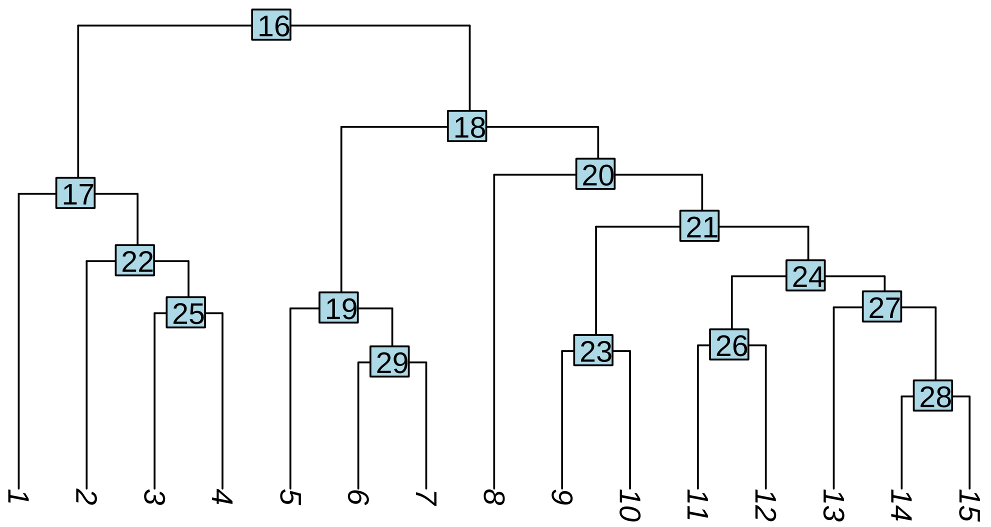


```r
# Repeate assess 25% bottom nodes out of bag error, using
# 0.03 as merge criteria
oobe.new <- AssessNodes(tme, node.list = c(23, 26, 28, 29))

# No out of bag error aboce 0.03
print(oobe.new)
```

```
##   node        oobe
## 1   23 0.004201681
## 2   26 0.001121705
## 3   28 0.015409570
## 4   29 0.007756233
```

##### 3. Doublets Removal

```r
# Cluster 7 have higher UMI counts and gene detection.
VlnPlot(tme, features.plot = c("nUMI", "nGene", "Mki67"), nCol = 3,
    do.return = TRUE, point.size.use = 0, do.sort = TRUE, cols.use = cols.use)
```

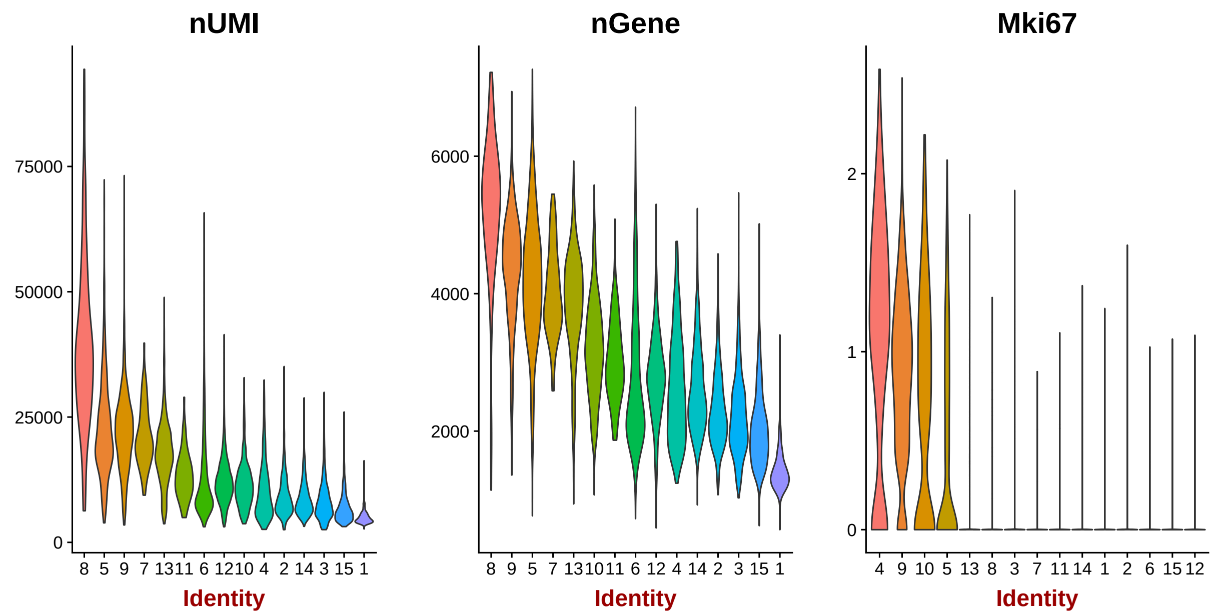


```r
# Cluster 7 expresses epithelial (Epcam), fibroblast (Acta2,
# Fn1) and basal-like (Cd24a) markers.
DotPlot(tme, genes.plot = c("Epcam", "Acta2", "Cd24a", "Fn1"),
    plot.legend = TRUE)
```

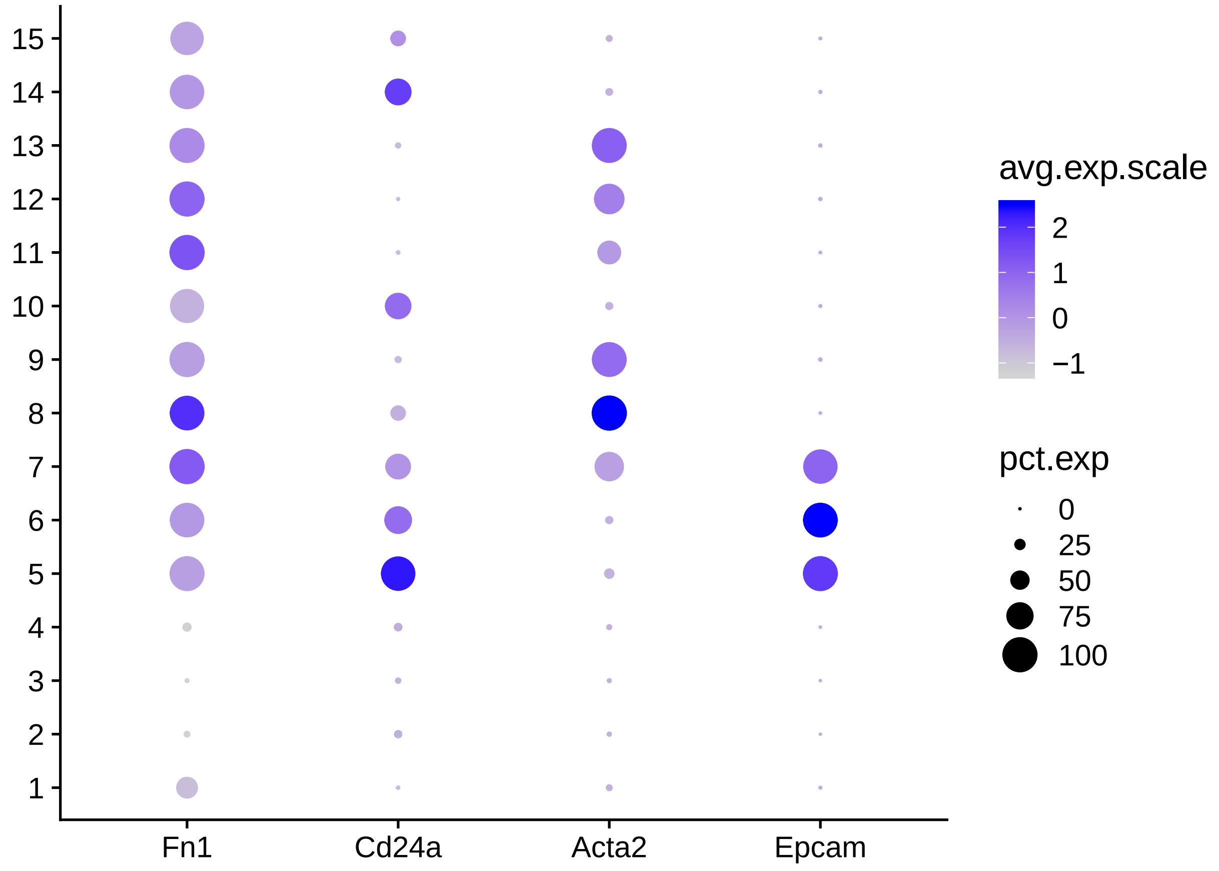

##### 4. Macrophage Subtype Clustering

```
# Subset macrophages.
macrophage <- SubsetData(tme, ident.use = c(1:4, 11), subset.raw = TRUE)
macrophage <- ScaleData(macrophage, display.progress = FALSE)
macrophage <- FindVariableGenes(macrophage, do.plot = FALSE,
    display.progress = FALSE)
macrophage <- RunPCA(macrophage, do.print = FALSE)

# optional PC elbow plot PCElbowPlot(tme)

macrophage <- RunTSNE(macrophage, dims.use = 1:10)
TSNEPlot(macrophage, do.label = TRUE, colors.use = cols.use)
```

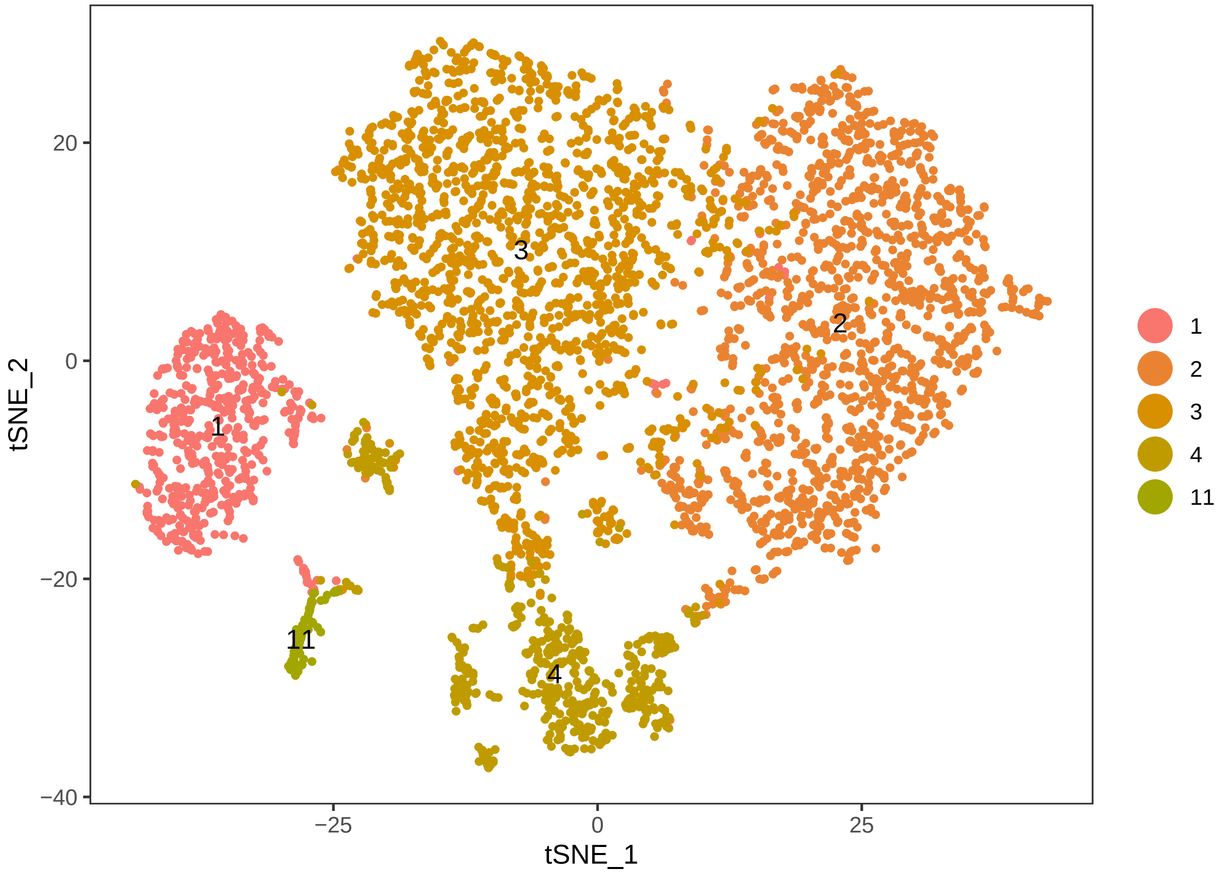


```
FeaturePlot(macrophage, features.plot = c("Cd68", "Ctss", "Csf1r",
    "Arg1", "Itgax", "Mki67"), nCol = 3)
```

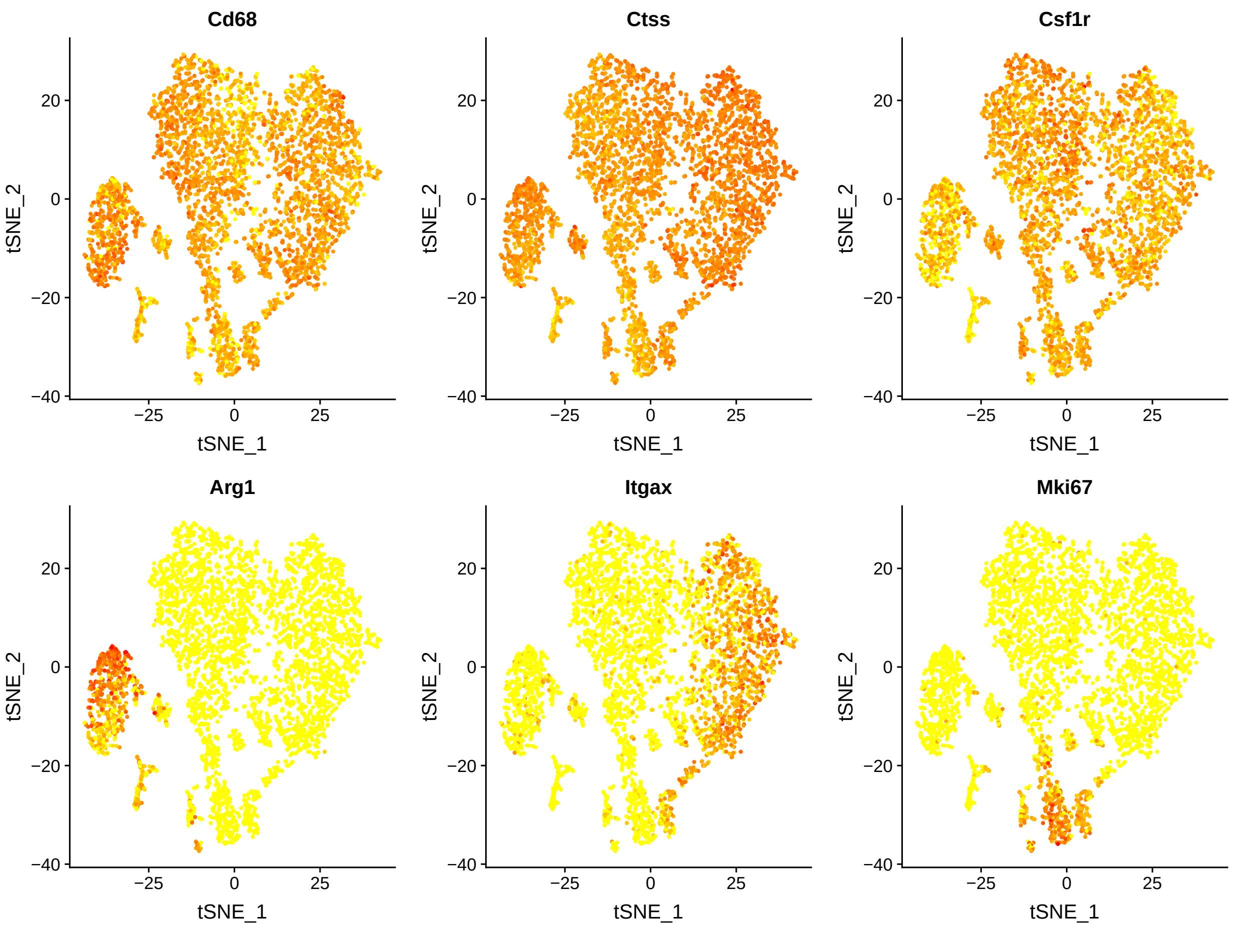


```
# Subset cycling macrophage for further clustering.
macrophage.cycling <- SubsetData(macrophage, ident.use = 4, subset.raw = TRUE)

# Repeat the clustering steps as above with 10PCs and an out
# of bag error of 0.07.
macrophage.cycling <- RunPCA(macrophage.cycling, do.print = FALSE)
macrophage.cycling <- FindClusters(macrophage.cycling, dims.use = 1:10,
    resolution = 1, save.SNN = TRUE, print.output = FALSE)
macrophage.cycling <- RunTSNE(macrophage.cycling, dims.use = 1:10)
macrophage.cycling <- BuildClusterTree(macrophage.cycling, do.reorder = TRUE,
    reorder.numeric = TRUE, pcs.use = 1:10)
```

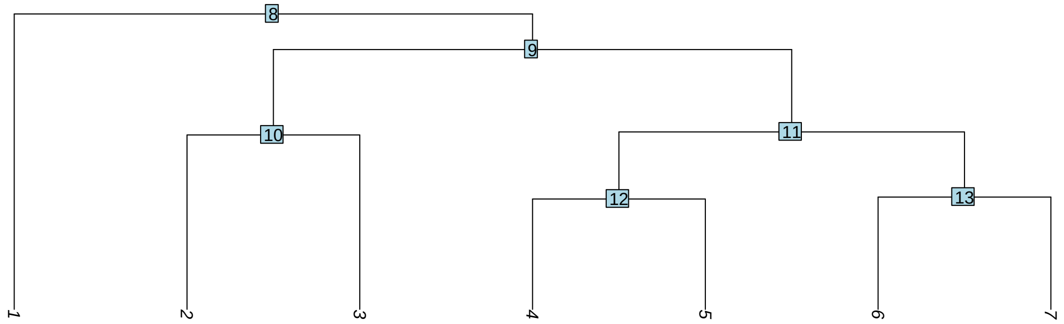


```
oobe.mac <- AssessNodes(macrophage.cycling, node.list = c(12,
    13))
print(oobe.mac)
```

```
##   node       oobe
## 1   12 0.09714286
## 2   13 0.13173653
```

```
macrophage.cycling <- MergeNode(macrophage.cycling, node.use = 12)
macrophage.cycling <- MergeNode(macrophage.cycling, node.use = 13)
macrophage.cycling <- BuildClusterTree(macrophage.cycling, do.reorder = TRUE,
    reorder.numeric = TRUE, pcs.use = 1:10)
```

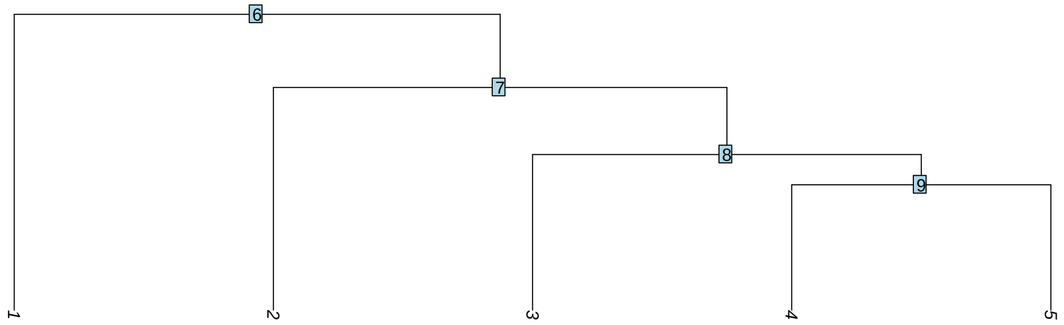


```
oobe.mac <- AssessNodes(macrophage.cycling, node.list = 9)
print(oobe.mac)
```

```
##   node       oobe
## 1    9 0.07017544
```

```
macrophage.cycling <- MergeNode(macrophage.cycling, node.use = 9)
macrophage.cycling <- BuildClusterTree(macrophage.cycling, do.reorder = TRUE,
    reorder.numeric = TRUE, pcs.use = 1:10)
```

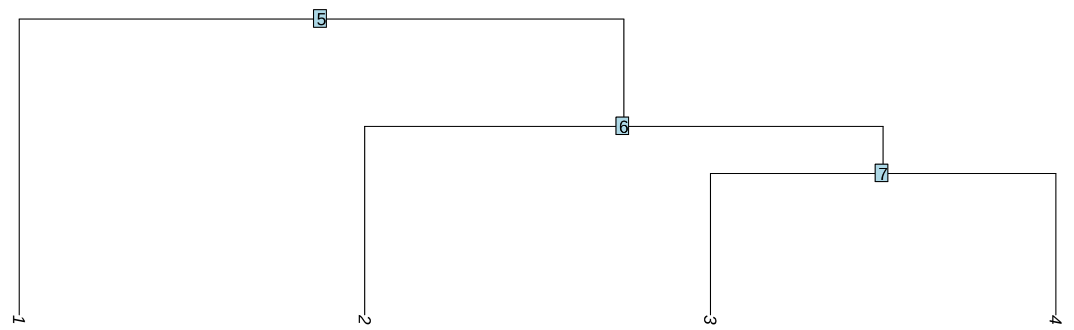


```
oobe.mac <- AssessNodes(macrophage.cycling)
print(oobe.mac)
```

```
##   node       oobe
## 1    5 0.03469388
## 2    6 0.01647059
## 3    7 0.06818182
```

```
p1 <- TSNEPlot(macrophage.cycling, do.label = TRUE, do.return = TRUE,
    colors.use = cols.use)
p2 <- TSNEPlot(macrophage.cycling, do.label = TRUE, group.by = "orig.ident",
    do.return = TRUE, colors.use = cols.use)
plot_grid(p1, p2, align = "hv", ncol = 2)
```

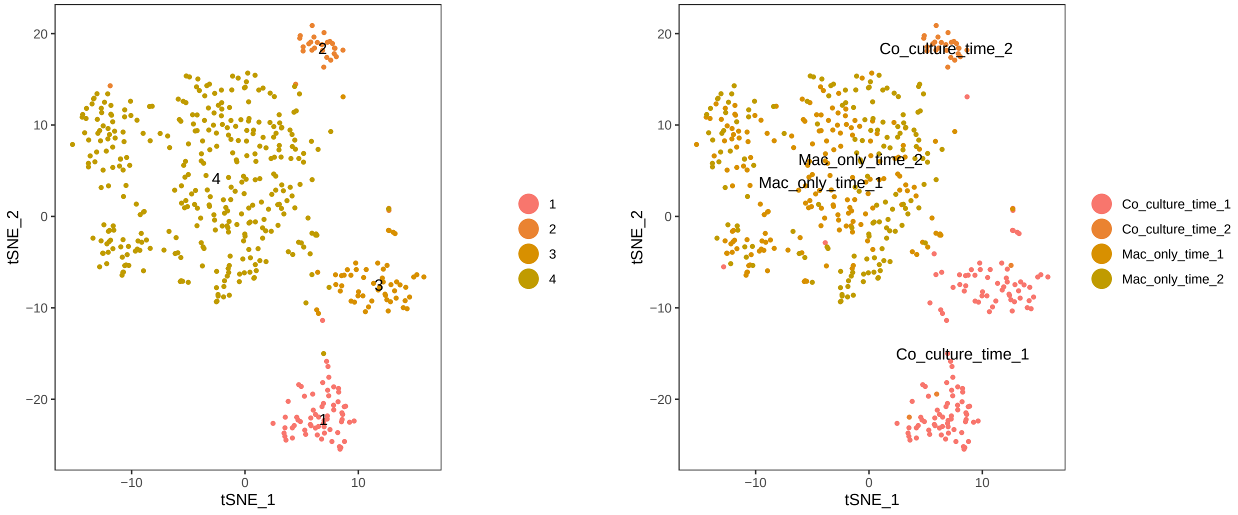

```r
p1 <- VlnPlot(macrophage.cycling, features.plot = c("Cd68", "Ctss",
    "Arg1", "Itgax", "Mki67"), point.size.use = 0.01, do.return = TRUE,
    cols.use = cols.use)
p2 <- ggplot(macrophage.cycling@meta.data, aes(x = macrophage.cycling@ident,
    fill = orig.ident)) + geom_bar(position = "fill", show.legend = T) +
    labs(x = "Cluster ID", y = "Percentage", fill = "orig.ident") +
    theme(axis.title = element_text(face = "bold"), axis.text.x = element_text(vjust = 0.9,
        hjust = 1, angle = 45, size = 15), legend.title = element_text(face = "bold"))
cowplot::plot_grid(p1, p2)
```

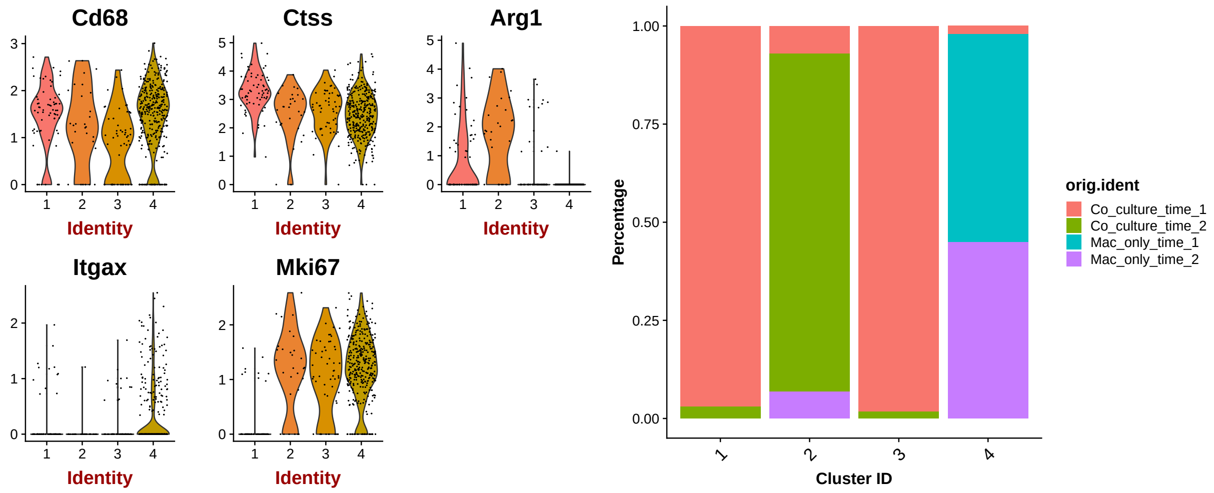


```r
# Rename according to cell cycle, education status and time
# point.
macrophage.cycling <- RenameIdent(macrophage.cycling, 1, "Early")
macrophage.cycling <- RenameIdent(macrophage.cycling, 2, "Late-Cycling")
macrophage.cycling <- RenameIdent(macrophage.cycling, 3, "Early-Cycling")
macrophage.cycling <- RenameIdent(macrophage.cycling, 4, "Naive-Cycling")

# Cluster 11 has higher UMI and gene detection.
VlnPlot(macrophage, c("nUMI", "nGene"), do.sort = TRUE, point.size.use = 0,
    cols.use = cols.use)
```

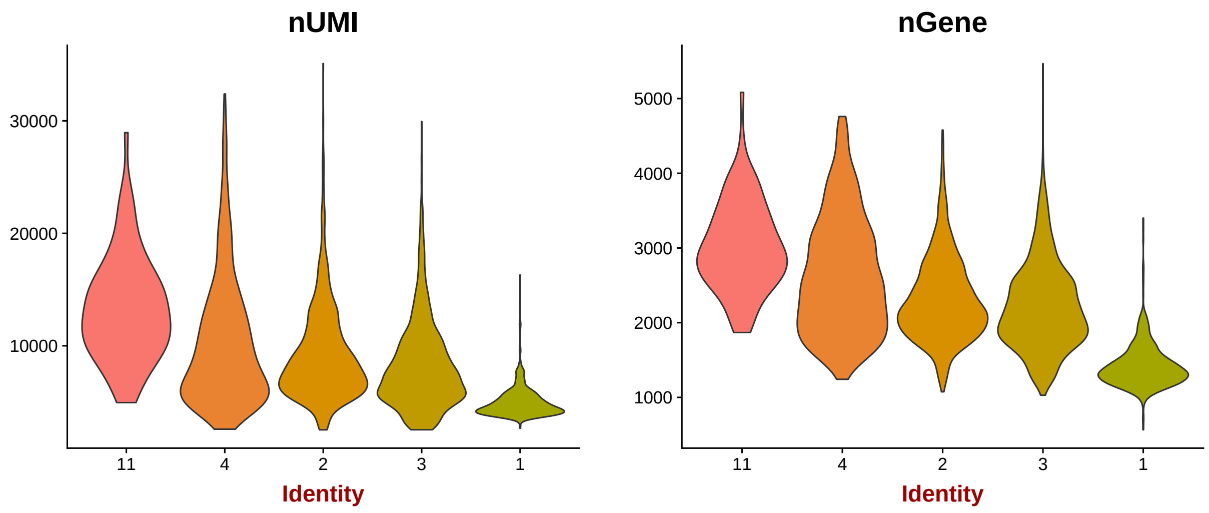

```r
# Cluster 11 expresses CAF (Acta2, Fn1) and Macrophage
# markers (Cd68, Ctss, Arg1)
DotPlot(macrophage, genes.plot = c("Cd68", "Ctss", "Arg1", "Itgax",
    "Mki67", "Acta2", "Fn1"), plot.legend = TRUE, group.by = "res.1")
```

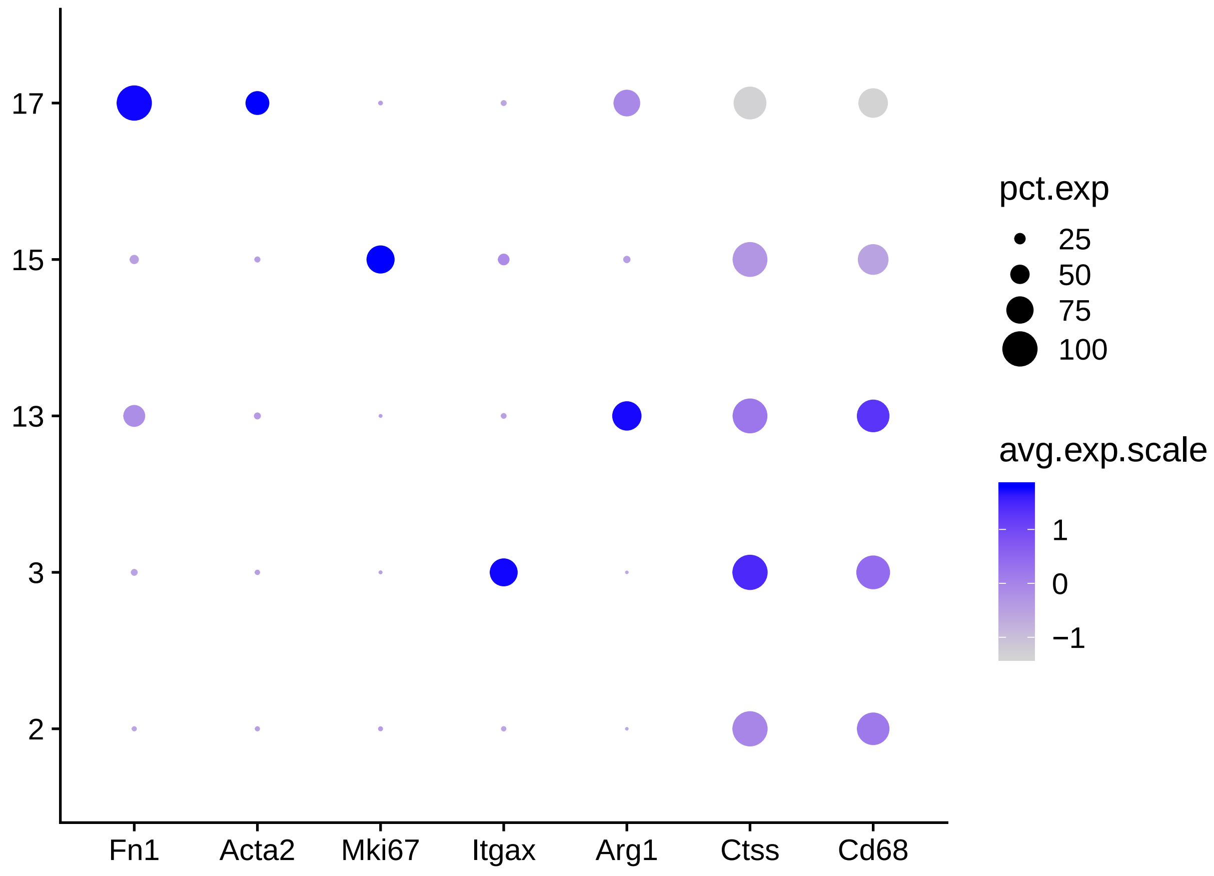


```r
# Remove doublet cells.
macrophage <- SubsetData(macrophage, ident.remove = 11)
tme <- SubsetData(tme, ident.remove = 11)

# Rename clusters.
macrophage <- RenameIdent(macrophage, 1, "Late")
macrophage <- RenameIdent(macrophage, 2, "Naive CD11c")
macrophage <- RenameIdent(macrophage, 3, "Naive")
macrophage <- SetIdent(macrophage, cells.use = macrophage.cycling@cell.names,
    ident.use = macrophage.cycling@ident)

# Visualization Subset macrophage only condition time point 2
# for re-embeding and visualization.
macrophage.vis <- SetIdent(macrophage, ident.use = macrophage@meta.data$orig.ident)
macrophage.vis <- SubsetData(macrophage.vis, ident.remove = "Mac_only_time_2",
    subset.raw = T)
macrophage.vis <- ScaleData(macrophage.vis, display.progress = FALSE)
macrophage.vis <- FindVariableGenes(macrophage.vis, do.plot = FALSE,
    display.progress = FALSE, top.genes = 2000)
```

```r
macrophage.vis <- RunPCA(macrophage.vis, do.print = FALSE, pcs.compute = 30)
macrophage.vis <- RunTSNE(macrophage.vis, dims.use = 1:30)
macrophage.vis <- SetIdent(macrophage.vis, ident.use = macrophage@ident[macrophage.vis@cell.names])

# For visualization purpose.
macrophage.vis@dr$tsne@cell.embeddings[, 2] <- -macrophage.vis@dr$tsne@cell.embeddings[,
    2]
macrophage.vis@ident <- factor(macrophage.vis@ident, levels = c("Naive CD11c",
    "Naive", "Naive-Cycling", "Early", "Early-Cycling", "Late",
    "Late-Cycling"), ordered = TRUE)
col4 <- c("#A6CEE3", "#1F78B4", "#B2DF8A", "#33A02C")
col7 <- c("#1B9E77", "#808080", "#F0027F", "#BF5B17", "#A6D854",
    "#FF7F00", "#377EB8")
p1 <- DimPlot(macrophage.vis, reduction.use = "tsne", dim.1 = 2,
    dim.2 = 1, cols.use = col7, pt.size = 1, do.return = TRUE,
    no.axes = TRUE)
p2 <- DimPlot(macrophage.vis, reduction.use = "tsne", dim.1 = 2,
    dim.2 = 1, cols.use = col4, group.by = "orig.ident", pt.size = 1,
    do.return = TRUE, no.axes = TRUE)
p3 <- ggplot(macrophage@meta.data, aes(x = macrophage@ident,
    fill = orig.ident)) + geom_bar(position = "fill", show.legend = TRUE) +
    scale_fill_manual(values = col4) + labs(x = "Cluster ID",
    y = "Percentage", fill = "orig.ident") + theme(axis.title = element_text(face = "bold"),
    axis.text.x = element_text(vjust = 0.9, hjust = 1, angle = 45,
        size = 15), legend.title = element_text(face = "bold"))
plot_grid(p1, p2, p3, align = "hv", ncol = 3)
```

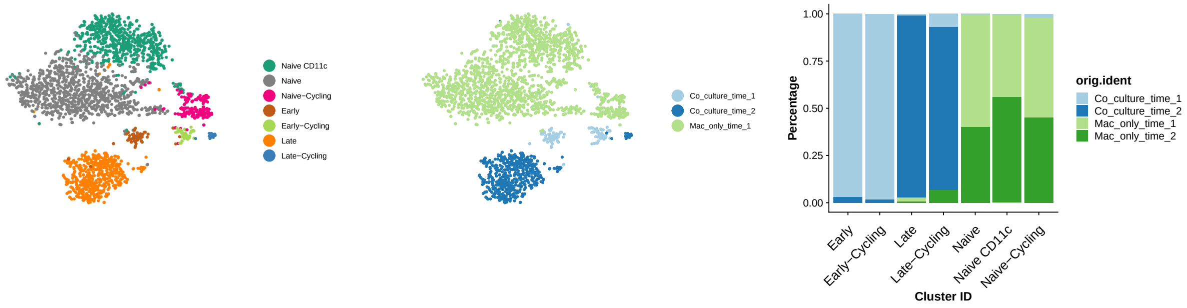


##### 5.Final Clustering Results

```r
# Rename clusters.
tme <- SetIdent(tme, WhichCells(tme, 5), "Tumor Cycling")
tme <- SetIdent(tme, WhichCells(tme, 6), "Tumor 1")
tme <- SetIdent(tme, WhichCells(tme, 9), "CAF Cycling")
tme <- SetIdent(tme, WhichCells(tme, 12), "CAF 1")
tme <- SetIdent(tme, WhichCells(tme, 13), "CAF 2")
tme <- SetIdent(tme, WhichCells(tme, 8), "CAF 3")
tme <- SetIdent(tme, WhichCells(tme, 10), "Basal Cycling")
tme <- SetIdent(tme, WhichCells(tme, 14), "Basal 1")
tme <- SetIdent(tme, WhichCells(tme, 15), "Basal 2")
tme <- SetIdent(tme, macrophage@cell.names, macrophage@ident)
```

```
# Rename lineage.
tme@meta.data[["lineage"]] <- "Macrophage"
tme@meta.data[WhichCells(tme, ident = c("CAF 1", "CAF 2", "CAF 3",
    "CAF Cycling")), "lineage"] <- "CAF"
tme@meta.data[WhichCells(tme, ident = c("Basal 1", "Basal 2",
    "Basal Cycling")), "lineage"] <- "Basal"
tme@meta.data[WhichCells(tme, ident = c("Tumor 1", "Tumor Cycling")),
    "lineage"] <- "Tumor"
tme@meta.data[["cell.type"]] <- tme@ident

# tSNE visualization
p1 <- TSNEPlot(tme, do.label = FALSE, do.return = TRUE, group.by = "orig.ident",
    colors.use = cols.use)
p2 <- TSNEPlot(tme, do.label = FALSE, do.return = TRUE, group.by = "condition",
    colors.use = cols.use)
p3 <- TSNEPlot(tme, do.label = FALSE, do.return = TRUE, group.by = "lineage",
    colors.use = cols.use)
p4 <- TSNEPlot(tme, do.label = FALSE, do.return = TRUE, group.by = "cell.type",
    colors.use = cols.use)
plot_grid(p1, p2, p3, p4, ncol = 2, align = "hv")
```

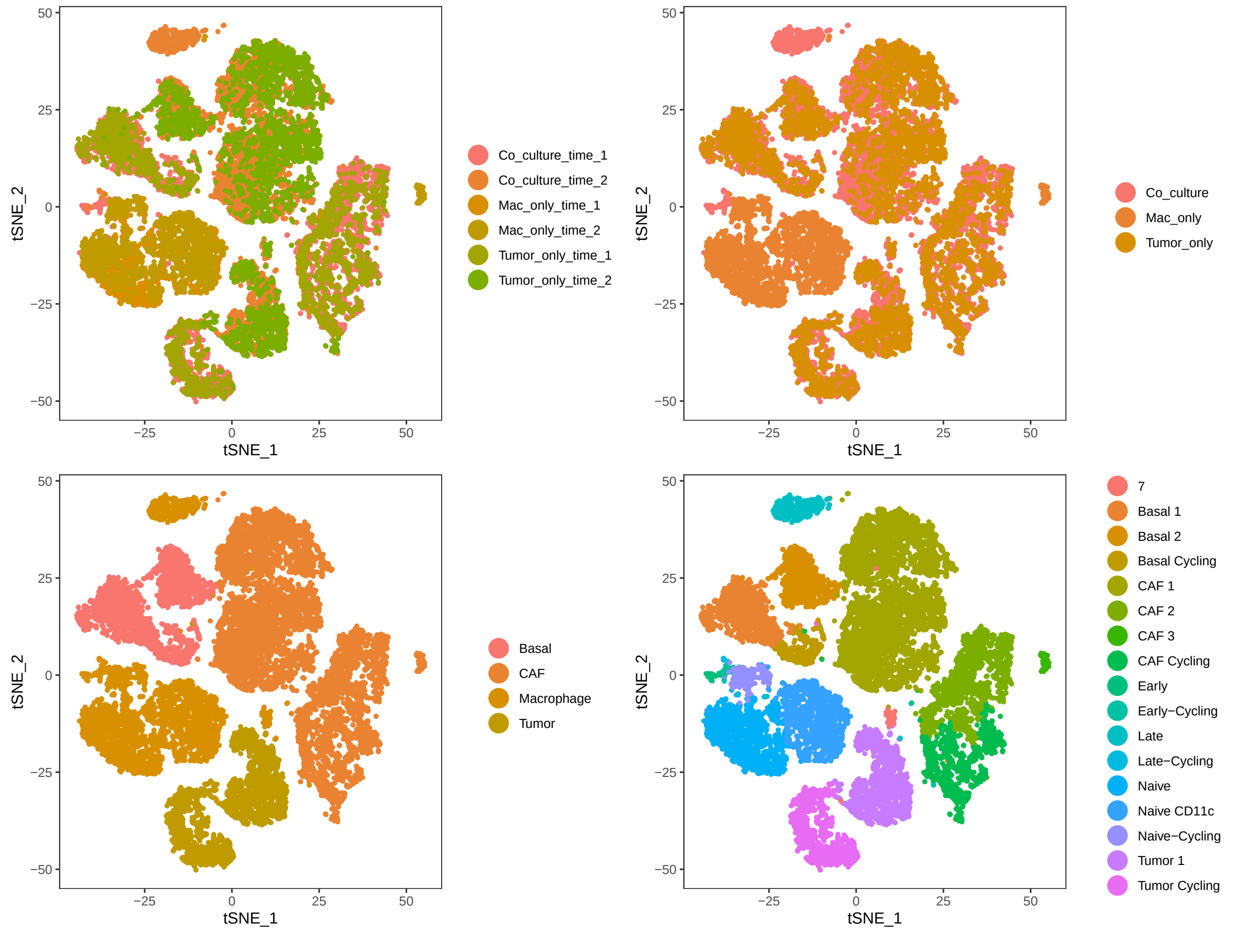

```r
FeaturePlot(tme, features.plot = c("Epcam", "Cd24a", "Acta2",
    "Cd68"), nCol = 2, reduction.use = "tsne", do.return = FALSE,
    no.legend = FALSE, pt.size = 1)
```

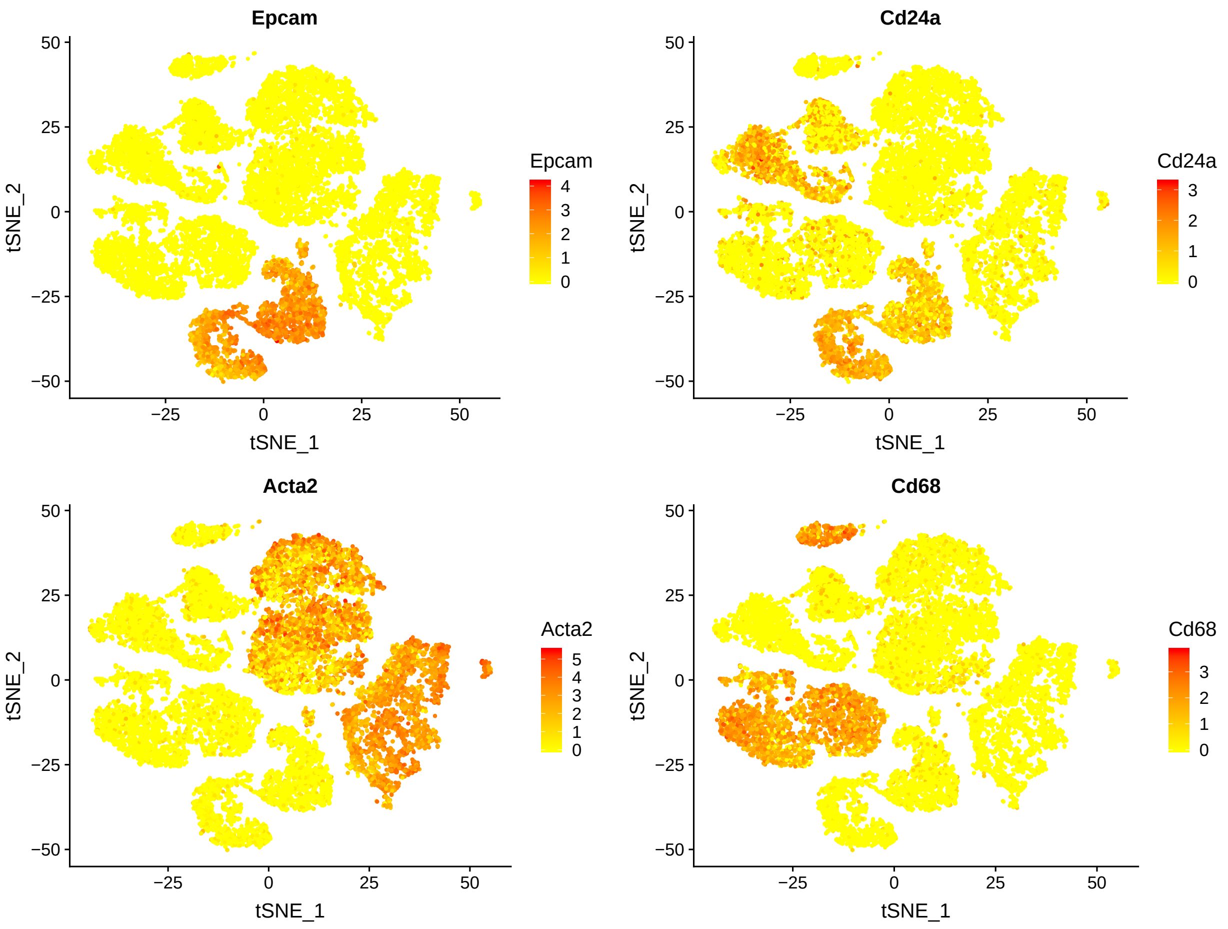


#### 6. Differential Expression

```r
# Lineage markers.
tme <- SetIdent(tme, ident.use = tme@meta.data$lineage)
lineage.markers <- FindAllMarkers(tme, logfc.threshold = 1, min.pct = 0.5,
    only.pos = TRUE, print.bar = FALSE)

# CAF subtype markers.
caf <- SubsetData(tme, ident.use = "CAF")
caf <- SetIdent(caf, ident.use = caf@meta.data$cell.type)
caf.markers <- FindAllMarkers(caf, logfc.threshold = 0.75, min.pct = 0.4,
    only.pos = TRUE, print.bar = FALSE)

# Tumor subtype markers.
tumor <- SubsetData(tme, ident.use = "Tumor")
tumor <- SetIdent(tumor, ident.use = tumor@meta.data$cell.type)
tumor.markers <- FindAllMarkers(tumor, logfc.threshold = 0.75,
```

```
    min.pct = 0.4, only.pos = TRUE, print.bar = FALSE)

# Basal subtype markers.
basal <- SubsetData(tme, ident.use = "Basal")
basal <- SetIdent(basal, ident.use = basal@meta.data$cell.type)
basal.markers <- FindAllMarkers(basal, logfc.threshold = 0.75,
    min.pct = 0.4, only.pos = TRUE, print.bar = FALSE)

# Cycling macrophage subtype markers.
cycling.macrophage <- SubsetData(macrophage, ident.use = c("Early-Cycling",
    "Late-Cycling", "Naive-Cycling"))
cycling.macrophage.markers <- FindAllMarkers(cycling.macrophage,
    logfc.threshold = 0.75, min.pct = 0.4, only.pos = TRUE, print.bar = FALSE)

# Non-cycling macrophage subtype markers.
noncycling.macrophage <- SubsetData(macrophage, ident.use = c("Early",
    "Late", "Naive", "Naive CD11c"))
noncycling.macrophage.markers <- FindAllMarkers(noncycling.macrophage,
    logfc.threshold = 0.75, min.pct = 0.4, only.pos = TRUE, print.bar = FALSE)
DoHeatmap(tme, genes.use = lineage.markers$gene, slim.col.label = TRUE,
    remove.key = TRUE, group.label.rot = TRUE, title = "Differential Expression Genes",
    cex.row = 4)
```

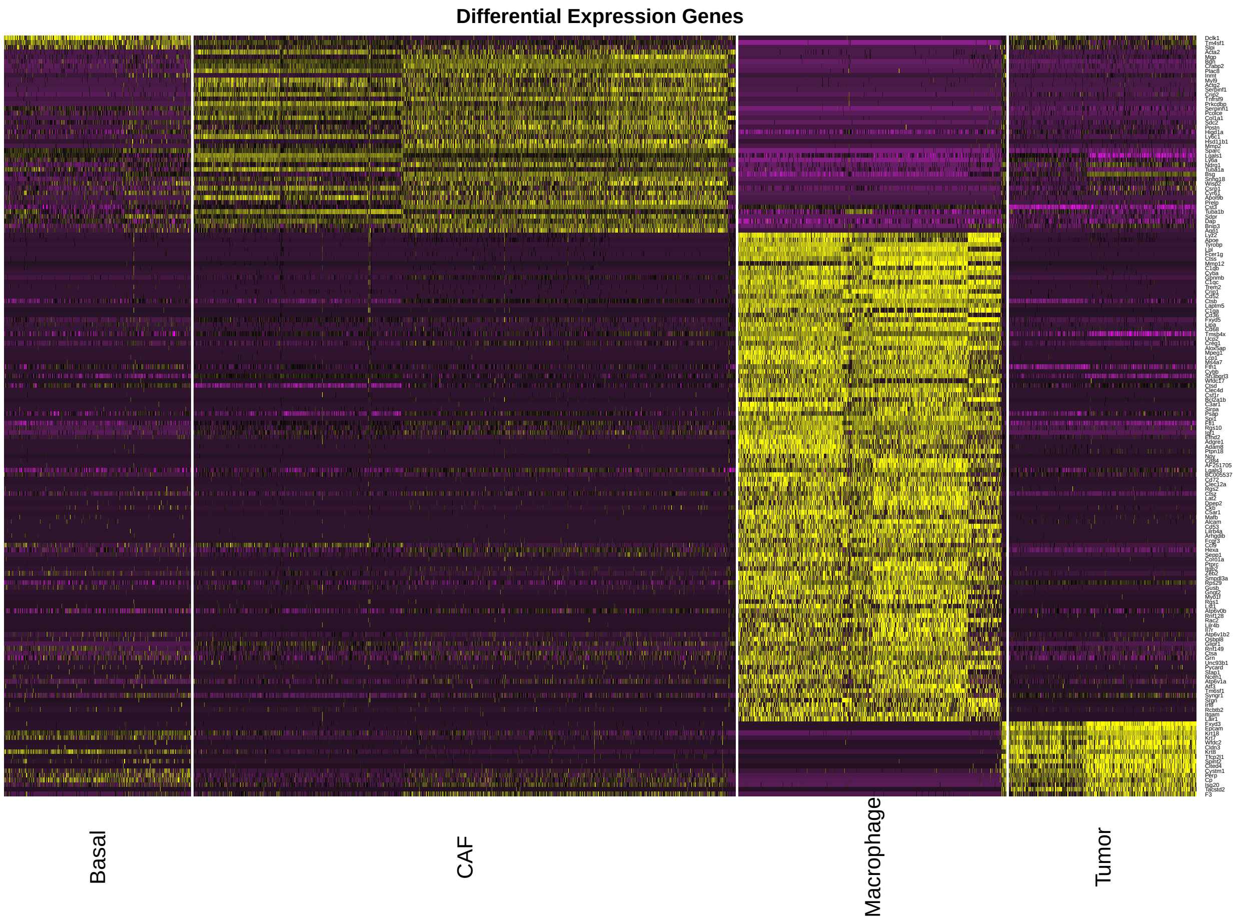

```r
macrophage@ident <- factor(macrophage@ident, levels = c("Naive CD11c",
    "Naive", "Naive-Cycling", "Early", "Early-Cycling", "Late",
    "Late-Cycling"), ordered = TRUE)
PlotVln(macrophage, features.plot = rev(c("Mki67", "Itgax", "Ccl2",
    "Ccl4", "Ly6a", "Csf2ra", "Fn1", "Arg1", "F13a1")), plot.group = T,
    cols.use = col7, alpha = 0.5, nCol = 1, do.jitter = F, align = "hv",
    blank.x = T)
```

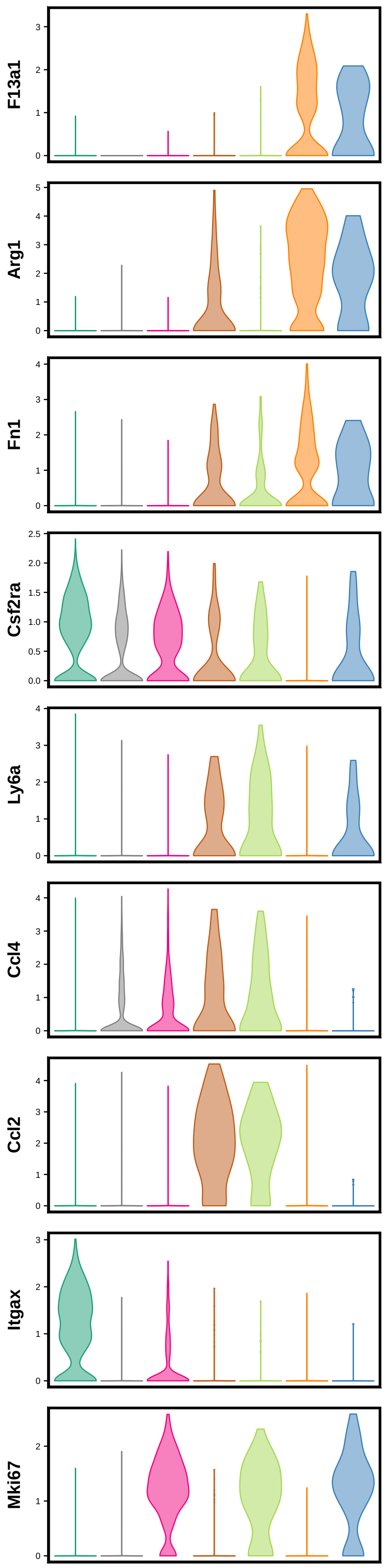
F13a1
Arg1
Fn1
Csf2ra
Ly6a
Ccl4
Ccl2
Itgax
Mki67

```r
# Test all the genes between early and late educated
# macrophages for differential expression.
all.education.de <- FindMarkers(macrophage, ident.1 = "Late",
    ident.2 = "Early", logfc.threshold = 0, min.pct = 0, print.bar = FALSE)
all.education.de$p_val_fdr <- p.adjust(all.education.de$p_val,
    method = "BH")
all.education.de$log_fdr <- -log(all.education.de$p_val_fdr,
    10)
all.education.de$gene.name <- rownames(all.education.de)
all.education.de$diff_pct <- abs(all.education.de$pct.1 - all.education.de$pct.2)

# Volcano plot
ggplot(all.education.de, aes(avg_logFC, log_fdr)) + geom_point(aes(col = (avg_logFC),
    size = abs(diff_pct)), alpha = 1) + scale_size_area(max_size = 10,
    name = "Differential %") + scale_color_gradientn(colours = c("blue4",
    "grey", "red3"), name = "logFC levels") + ylim(0, 70) + geom_vline(xintercept = c(0.75,
    -0.75), lty = 2, lwd = 1) + geom_hline(yintercept = -log(0.01,
    10), lty = 2, lwd = 1) + geom_label_repel(aes(label = ifelse(avg_logFC >
    0.75 & log_fdr > -log10(0.01) & diff_pct > 0.3, as.character(gene.name),
    "")), hjust = -1.5, vjust = 0.8, segment.color = "red", colour = "red",
    size = 4) + geom_label_repel(aes(label = ifelse(avg_logFC <
    (-1) & log_fdr > -log10(0.01) & diff_pct > 0.4, as.character(gene.name),
    "")), hjust = 3, vjust = -0.8, segment.color = "blue", colour = "blue",
    size = 4) + xlab("Average logFC") + ylab("-log(FDR)") + theme(plot.title = element_text(size = 20),
    axis.title = element_text(size = 20, face = 2), legend.title = element_text(size = 18),
    legend.text = element_text(size = 15))
```

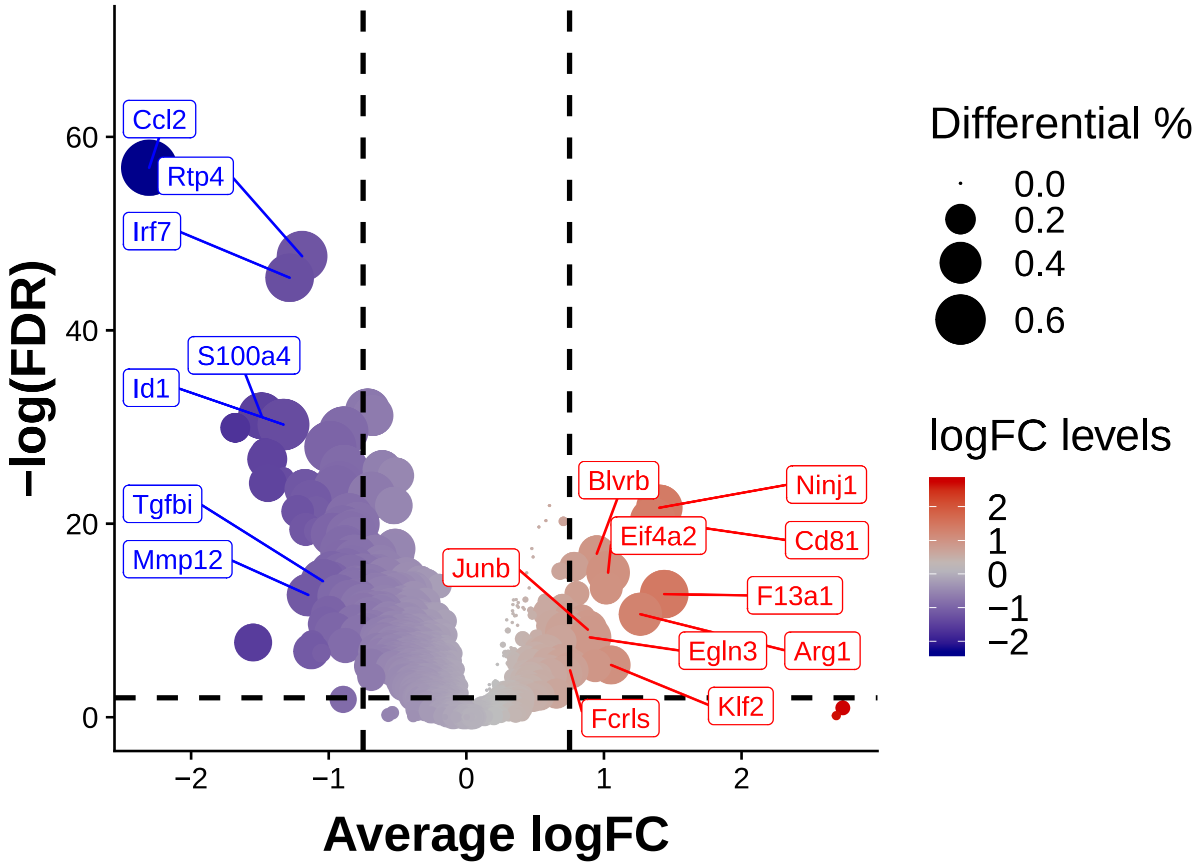


7. **Macrophage Education**

```
macrophage <- ScaleData(macrophage, display.progress = FALSE)
macrophage@meta.data$orig.ident <- factor(macrophage@meta.data$orig.ident,
    levels = c("Mac_only_time_1", "Mac_only_time_2", "Co_culture_time_1",
        "Co_culture_time_2"), ordered = TRUE)

# Get differential expressed genes of naive and educated
# macrophage
naive.genes <- FindMarkers(macrophage, "Naive CD11c", logfc.threshold = 0.75,
    min.pct = 0.4, test.use = "MAST", only.pos = TRUE, print.bar = FALSE)
education.genes <- FindMarkers(macrophage, "Late", logfc.threshold = 0.75,
    min.pct = 0.4, test.use = "MAST", only.pos = TRUE, print.bar = FALSE)
marker.genes <- c(rownames(naive.genes), rownames(education.genes))
education.dm <- DiffusionMap(t(x = macrophage@scale.data[marker.genes,
    ]), verbose = FALSE)

# Rotate the axis
if (education.dm@eigenvectors[1, 1] < 0) education.dm@eigenvectors[,
    1] <- -education.dm@eigenvectors[, 1]
if (education.dm@eigenvectors[1, 2] > 0) education.dm@eigenvectors[,
    2] <- -education.dm@eigenvectors[, 2]
rownames(education.dm@eigenvectors) <- macrophage@cell.names
```

```
macrophage <- SetDimReduction(macrophage, reduction.type = "dm",
    slot = "cell.embeddings", new.data = education.dm@eigenvectors)

# Pseudotime calculation
pseudotime <- principal_curve(education.dm@eigenvectors[, 1:2],
    start = education.dm@eigenvectors[, c(1, 1)], smoother = "smooth_spline")
# Normalize to 0-1 range
macrophage@meta.data$pseudotime <- (max(pseudotime$lambda) -
    pseudotime$lambda)/(max(pseudotime$lambda) - min(pseudotime$lambda))
macrophage@dr$dm@key = "DC"

p1 <- DimPlot(macrophage, reduction.use = "dm", cols.use = cols.use,
    do.return = TRUE) + geom_line(mapping = aes(DC1, DC2), data = as.data.frame(pseudotime$s))
p2 <- Pseudo.plot(macrophage, gene = "Arg1", reduction = "dm",
    col.low = "grey", col.high = "red", dot.min = 1, dot.max = 5)
p3 <- PlotVln(macrophage, features.plot = "pseudotime", group.by = "orig.ident",
    plot.quantile = TRUE, color.by.expr = TRUE, do.jitter = FALSE,
    x.lab.rot = TRUE) + ylab("")
cowplot::plot_grid(p1, p2, p3, nrow = 1, rel_widths = c(1.5,
    1.5, 1), rel_heights = c(1, 1, 2))
```

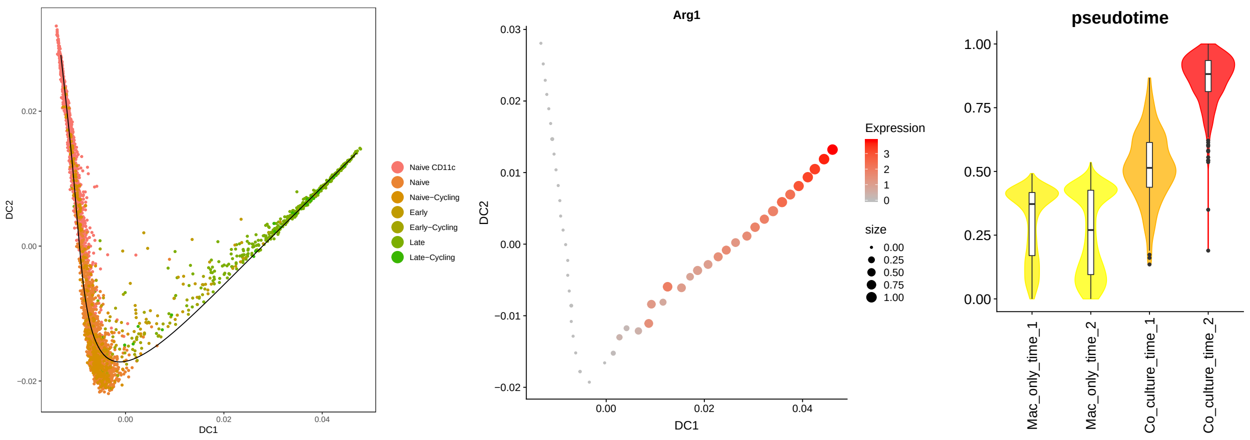


```
# Top 1000 highly varible gene expression profile on
# pseudotime axis.
trajectory.data <- macrophage@data[rownames([1:1000,
    ]), order(macrophage@meta.data$pseudotime, decreasing = FALSE)]
pseudotime <- sort(macrophage@meta.data$pseudotime, decreasing = FALSE)

# Smoothed expression profile.
trajectory.data <- apply(as.matrix(trajectory.data), 1, function(x) loess(x ~
    pseudotime, span = 0.7)$fitted)

# Mutual information
mi.data <- apply(trajectory.data, 2, function(x) mutinformation(discretize(pseudotime,
    disc = "equalfreq"), discretize(x, disc = "equalfreq")))
trajectory.data <- trajectory.data[, names(which(mi.data > quantile(mi.data,
    0.25)))]

# kmeans clustering
trajectory.data <- t(scale(trajectory.data))
```

```r
set.seed(1234)
kmeans.data <- kmeans(trajectory.data, centers = 3, nstart = 25)

# Bin data according to pseudotime
macrophage.bin <- SetIdent(macrophage, macrophage@cell.names,
    ggplot2::cut_interval(macrophage@meta.data$pseudotime, 20))
macrophage.bin@ident <- factor(macrophage.bin@ident, levels = levels(ggplot2::cut_interval(macrophage@m[illegible]
    20)), ordered = TRUE)
macrophage.bin <- AverageExpression(macrophage.bin, return.seurat = TRUE,
    show.progress = FALSE)

# Heatmap
DoHeatmap(macrophage.bin, genes.use = names(sort(kmeans.data$cluster)),
    cex.row = 0, slim.col.label = TRUE, group.by = NULL, remove.key = TRUE)
```

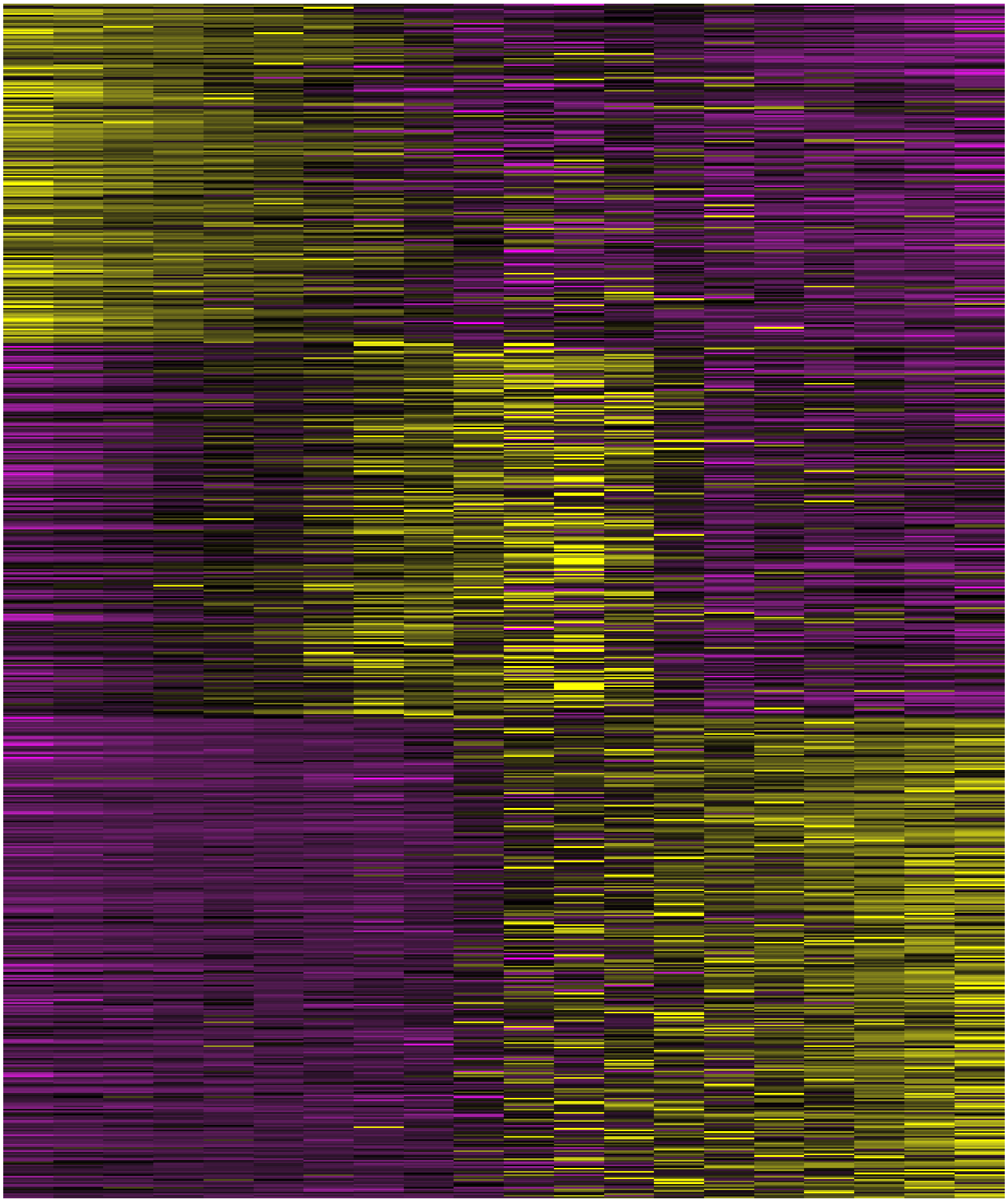

#### 8. Human Breast Cancer scRNA-seq

We used the human breast cancer scRNA-seq data from *Single-Cell Map of Diverse Immune Phenotypes in the Breast Tumor Microenvironment*. The expression data can be found on the GEO and the cell type annotation can be found in Table S2.

```
# Load data
brca.data <- data.table::fread("TME_Paper_Data/GSE114725_rna_raw.csv.gz",
    verbose = FALSE, showProgress = FALSE)
brca.label <- brca.data[, 1:5]
```

```r
brca.data <- brca.data[, -c(1:5)]
brca.data <- Matrix::Matrix(as.matrix(brca.data), sparse = TRUE)
rownames(brca.data) <- brca.label$cellid
brca <- CreateSeuratObject(Matrix::t(brca.data), min.cells = 0,
    min.genes = 0)
brca@meta.data <- cbind(brca@meta.data, brca.label)
brca <- SetIdent(brca, ident.use = brca@meta.data$cluster)

# Subset the monocytic population (monocytes, macrophages,
# dendritic cells, mDC, pDC)
annotation <- read.csv("TME_Paper_Data/Cluster-Annotations-Table.csv",
    header = TRUE, stringsAsFactors = FALSE)
monocytic <- SubsetData(brca, ident.use = annotation$ClusterID[grep("MONO|MAC|mDC|pDC",
    annotation$Final.Annotation.based.on.bulk.combined.with.differential.expressed.genes)])
monocytic <- NormalizeData(monocytic, display.progress = FALSE)
monocytic <- RenameIdent(monocytic, "23", "TAM_23")
monocytic <- RenameIdent(monocytic, "25", "TAM_25")
monocytic <- RenameIdent(monocytic, "28", "TAM_28")
monocytic <- SetIdent(monocytic, cells.use = monocytic@cell.names[which(!monocytic@ident %in%
    c("TAM_23", "TAM_25", "TAM_28"))], ident.use = "Monocytic")
monocytic@meta.data$identity <- monocytic@ident

# Get education signatures.
genes.to.project <- FindMarkers(macrophage, "Late", "Naive CD11c",
    logfc.threshold = 0.75, min.pct = 0.4, print.bar = FALSE)
gene.names <- intersect(toupper(rownames(genes.to.project)),
    rownames(monocytic@data))
monocytic <- ScaleData(monocytic, genes.use = gene.names, display.progress = FALSE)

# PCA using only the ex-vivo mouse signature.
monocytic <- RunPCA(monocytic, pc.genes = gene.names, seed.use = 123,
    do.print = FALSE)
monocytic@meta.data$meta.cluster <- as.character(monocytic@ident)
monocytic@meta.data$meta.cluster[which(monocytic@ident != "Monocytic")] <- "TAMs"
monocytic@meta.data$meta.cluster <- factor(monocytic@meta.data$meta.cluster,
    levels = c("Monocytic", "TAMs"), ordered = TRUE)
education.genes <- rownames(genes.to.project)[which(genes.to.project$avg_logFC >
    0)]

# Calculate ex-vivo education signature score (enrichment).
monocytic <- AddModuleScore(monocytic, list(education.genes),
    n.bin = 10, ctrl.size = 100, enrich.name = c("Ex-vivo"),
    seed = 123)
macrophage <- AddModuleScore(macrophage, list(education.genes),
    n.bin = 10, ctrl.size = 100, enrich.name = c("Ex-vivo"),
    seed = 123)
p1 <- DimPlot(monocytic, label = TRUE, group.by = "meta.cluster",
    reduction = "pca", cols.use = c("darkgrey", "orange"), pt.size = 1.5,
    label.size = 8, do.return = TRUE)
p2 <- PlotVln(monocytic, "Ex-vivo1", cols.use = c("darkgrey",
    "orange"), do.jitter = FALSE, plot.quantile = TRUE, group.by = "meta.cluster") +
    ylab("Module Score") + ggtitle("Ex vivo Education Signature")
cowplot::plot_grid(p1, p2, rel_widths = c(3, 2))
```

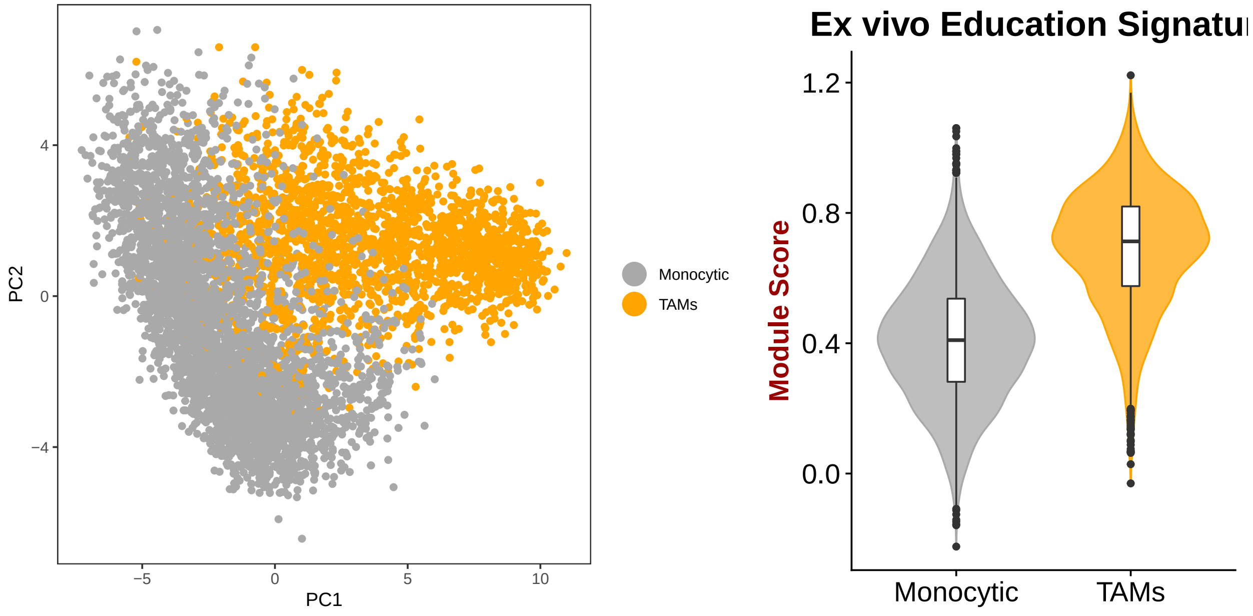


```
monocytic@ident <- factor(monocytic@ident, levels = c("Monocytic",
    "TAM_25", "TAM_23", "TAM_28"), ordered = TRUE)

# Human M2 signatures
human.signature <- FindMarkers(monocytic, c("TAM_23", "TAM_25",
    "TAM_28"), logfc.threshold = 0.75, min.pct = 0.4, print.bar = FALSE)

# Calculate human M2 signature score (enrichment).
monocytic <- AddModuleScore(monocytic, list(rownames(human.signature)[human.signature$avg_logFC >
    0]), n.bin = 10, ctrl.size = 100, enrich.name = c("human"),
    seed = 123)
p1 <- PlotVln(monocytic, "Ex-vivo1", cols.use = cols.use, do.jitter = FALSE,
    plot.quantile = TRUE, color.by.expr = TRUE, group.by = "ident") +
    ylab("Module Score") + ggtitle("Ex vivo Education Signature") +
    theme(legend.position = "right")
p2 <- ggplot(data = monocytic@meta.data, mapping = aes(human1,
    `Ex-vivo1`)) + geom_point(aes(color = identity)) + scale_colour_manual(values = c("darkblue",
    "lightblue2", "yellow2", "pink2")) + xlab("Ex-vivo Education Signature") +
    ylab("Human M2 Signature") + ggtitle(paste("Correlation =",
    round(cor(monocytic@meta.data[, "human1"], monocytic@meta.data[,
        "Ex-vivo1"]), digits = 2)))
cowplot::plot_grid(p1, p2, align = "hv")
```

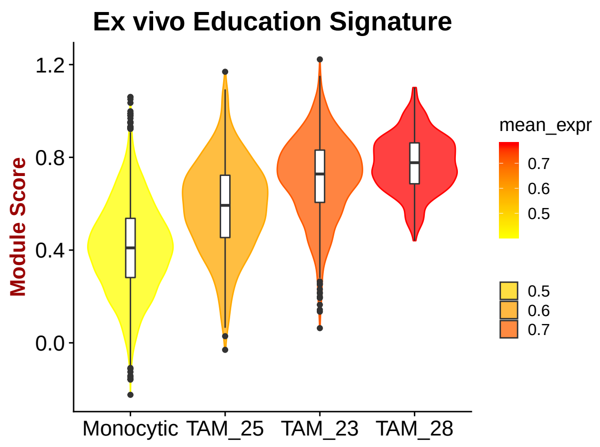


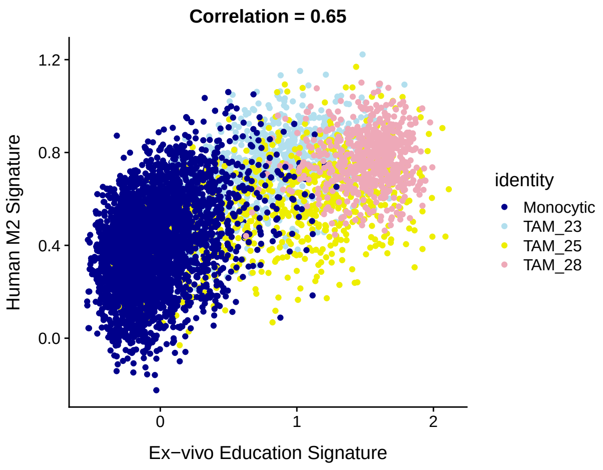


```r
# Session info
print(sessionInfo())
```

```
## R version 3.4.1 (2017-06-30)
## Platform: x86_64-redhat-linux-gnu (64-bit)
## Running under: CentOS Linux 7 (Core)
##
## Matrix products: default
## BLAS/LAPACK: /usr/lib64/R/lib/libRblas.so
##
## locale:
##  [1] LC_CTYPE=en_US.UTF-8       LC_NUMERIC=C
##  [3] LC_TIME=en_US.UTF-8        LC_COLLATE=en_US.UTF-8
##  [5] LC_MONETARY=en_US.UTF-8    LC_MESSAGES=en_US.UTF-8
##  [7] LC_PAPER=en_US.UTF-8       LC_NAME=C
##  [9] LC_ADDRESS=C               LC_TELEPHONE=C
## [11] LC_MEASUREMENT=en_US.UTF-8 LC_IDENTIFICATION=C
##
## attached base packages:
## [1] stats     graphics  grDevices utils     datasets  methods   base
##
## other attached packages:
##  [1] scales_1.0.0    dplyr_0.8.3     ggrepel_0.8.2   infotheo_1.2.0
##  [5] princurve_2.1.4 destiny_2.6.2   Seurat_2.3.4    Matrix_1.2-14
##  [9] cowplot_0.9.3   ggplot2_3.2.1
##
## loaded via a namespace (and not attached):
##   [1] readxl_1.3.1               snow_0.4-3
##   [3] backports_1.1.2            Hmisc_4.1-1
##   [5] RcppEigen_0.3.3.4.0        plyr_1.8.4
##   [7] igraph_1.2.2               lazyeval_0.2.2
##   [9] sp_1.3-1                   splines_3.4.1
##  [11] GenomeInfoDb_1.14.0        digest_0.6.18
##  [13] foreach_1.4.4              htmltools_0.3.6
##  [15] lars_1.2                   gdata_2.18.0
##  [17] magrittr_1.5               checkmate_1.8.5
##  [19] cluster_2.0.7-1            mixtools_1.1.0
##  [21] ROCR_1.0-7                 openxlsx_4.1.0
```

```
##  [23] matrixStats_0.54.0         R.utils_2.7.0
##  [25] xts_0.11-2                 colorspace_1.3-2
##  [27] haven_2.2.0                xfun_0.22
##  [29] RCurl_1.95-4.11            crayon_1.3.4
##  [31] jsonlite_1.6               survival_2.42-6
##  [33] zoo_1.8-4                  iterators_1.0.10
##  [35] ape_5.2                    glue_1.3.0
##  [37] gtable_0.2.0               XVector_0.18.0
##  [39] zlibbioc_1.24.0            DelayedArray_0.4.1
##  [41] car_3.0-2                  kernlab_0.9-27
##  [43] prabclus_2.2-6             BiocGenerics_0.24.0
##  [45] DEoptimR_1.0-8             abind_1.4-5
##  [47] VIM_4.8.0                  mvtnorm_1.0-8
##  [49] bibtex_0.4.2               Rcpp_1.0.1
##  [51] metap_1.0                  dtw_1.20-1
##  [53] laeken_0.5.0               htmlTable_1.12
##  [55] reticulate_1.10            foreign_0.8-71
##  [57] bit_1.1-14                 proxy_0.4-22
##  [59] mclust_5.4.1               SDMTools_1.1-221
##  [61] Formula_1.2-3              stats4_3.4.1
##  [63] tsne_0.1-3                 vcd_1.4-4
##  [65] htmlwidgets_1.3            httr_1.4.1
##  [67] gplots_3.0.1               RColorBrewer_1.1-2
##  [69] fpc_2.1-11.1               ellipsis_0.3.0
##  [71] acepack_1.4.1              modeltools_0.2-22
##  [73] ica_1.0-2                  pkgconfig_2.0.2
##  [75] R.methodsS3_1.7.1          flexmix_2.3-14
##  [77] nnet_7.3-12                labeling_0.3
##  [79] tidyselect_0.2.5           rlang_0.4.6
##  [81] reshape2_1.4.3             munsell_0.5.0
##  [83] cellranger_1.1.0           tools_3.4.1
##  [85] ranger_0.10.1              ggridges_0.5.0
##  [87] evaluate_0.14              stringr_1.4.0
##  [89] yaml_2.2.0                 npsurv_0.4-0
##  [91] knitr_1.25                 bit64_0.9-7
##  [93] fitdistrplus_1.0-11        zip_1.0.0
##  [95] robustbase_0.93-3          caTools_1.17.1.1
##  [97] purrr_0.3.3                RANN_2.6
##  [99] pbapply_1.3-4              nlme_3.1-137
## [101] formatR_1.5                R.oo_1.22.0
## [103] hdf5r_1.0.0                compiler_3.4.1
## [105] rstudioapi_0.10            curl_3.2
## [107] png_0.1-7                  e1071_1.7-0
## [109] lsei_1.2-0                 smoother_1.1
## [111] tibble_2.1.3               stringi_1.2.4
## [113] forcats_0.4.0              lattice_0.20-35
## [115] trimcluster_0.1-2.1        vctrs_0.3.0
## [117] pillar_1.4.3               lifecycle_0.1.0
## [119] Rdpack_0.9-0               lmtest_0.9-36
## [121] data.table_1.12.0          bitops_1.0-6
## [123] irlba_2.3.2                gbRd_0.4-11
## [125] GenomicRanges_1.30.3       R6_2.2.2
## [127] latticeExtra_0.6-28        KernSmooth_2.23-15
## [129] gridExtra_2.3              rio_0.5.10
```

```
## [131] IRanges_2.12.0              codetools_0.2-15
## [133] boot_1.3-20                 MASS_7.3-50
## [135] gtools_3.8.1                assertthat_0.2.0
## [137] SummarizedExperiment_1.8.1 MAST_1.4.1
## [139] rhdf5_2.22.0                withr_2.1.2
## [141] GenomeInfoDbData_1.0.0      S4Vectors_0.16.0
## [143] diptest_0.75-7              parallel_3.4.1
## [145] doSNOW_1.0.16               hms_0.5.2
## [147] grid_3.4.1                  rpart_4.1-13
## [149] tidyr_1.0.0                 class_7.3-14
## [151] rmarkdown_2.7               segmented_0.5-3.0
## [153] carData_3.0-1               Rtsne_0.13
## [155] TTR_0.23-4                  Biobase_2.38.0
## [157] base64enc_0.1-3
```
